# Polysaccharide utilization loci in *Bacteroides* determine population fitness and community-level interactions

**DOI:** 10.1101/2021.06.12.448201

**Authors:** Jun Feng, Yili Qian, Zhichao Zhou, Sarah Ertmer, Eugenio Vivas, Freeman Lan, Federico E. Rey, Karthik Anantharaman, Ophelia S. Venturelli

## Abstract

Polysaccharide utilization loci (PULs) in the human gut microbiome have critical roles in shaping human health and ecological dynamics. We develop a CRISPR-FnCpf1-RecT genome-editing tool to study 23 PULs in the highly abundant species *B. uniformis* (BU). We identify the glycan-degrading functions of multiple PULs and elucidate transcriptional coordination between PULs that enables the population to adapt to the loss of PULs. Exploiting a pooled BU mutant barcoding strategy, we demonstrate that the *in vitro* fitness and the colonization ability of BU in the murine gut is enhanced by deletion of specific PULs and modulated by glycan availability. We show that BU PULs can mediate complex glycan-dependent interactions with butyrate producers that depend on the mechanism of degradation and the butyrate producer glycan utilizing ability. In sum, PULs are major determinants of community dynamics and butyrate production and can provide a selective advantage or disadvantage depending on the nutritional landscape.

## INTRODUCTION

The human gut microbiome is a dynamic ecosystem shaped by a myriad of abiotic, microbial and host interactions. A core functionality of the human gut microbiome is to transform dietary and host derived polysaccharides (i.e. glycans) into metabolites that regulate our energy balance, provide colonization resistance to intestinal pathogens and maintain gut homeostasis^1-4^. While human cells lack the capability of utilizing complex dietary glycans, bacteria that inhabit the human gut harbor a broad repertoire of carbohydrate active enzymes (CAZymes) that can degrade these chemically diverse molecules, providing a unique metabolic function for the host^5-7^. Members of the *Bacteroides* genus are primary degraders of glycans due to the large number of CAZymes harbored within their genomes, which enables this genus to thrive as one of the most abundant and stable groups of organisms in the human gut^6^.

In *Bacteroides*, utilization of complex glycans is frequently mediated by polysaccharide utilization loci (PULs), which consist of sets of co-regulated genes for sensing nutrient availability (sensor-regulators), glycan capture (glycan binding proteins), uptake (oligosaccharide transporters) and digestion (CAZymes)^8^. PULs can have activity for a single glycan or a set of chemically similar glycans^9-12^. In some cases, more than one PUL contributes to the utilization of a chemically complex glycan^13,14^. The abundance of genome sequencing data has enabled the identification of many PULs in gut bacteria and facilitated their detailed biochemical characterization^10,15-19^. However, there remain a large number of PULs with unknown and uncharacterized biochemical functions^20^. Further, we have limited knowledge of the impact of a given PUL on microbial fitness in the context of the gut ecosystem.

The utilization of glycans by gut bacteria is a major driver of the ecological dynamics of gut microbiota. Competition for a given glycan can occur among species that are capable of utilizing the glycan, generating negative inter-species interactions^21^. By contrast, extracellular glycan digestion can lead to the release of polysaccharide breakdown products (PBPs) that can be utilized by specific members of the community, leading to a net positive outgoing interaction from the glycan utilizer to the recipient organism^22,23^. However, glycans can also be degraded via a selfish mechanism that does not release PBPs into the environment^22,24^. The potential release of PBPs as a public good for the community depends on the mechanism of degradation of the glycan and utilization ability of the recipient species.

While *Bacteroides* genomes contain many PULs enabling metabolic flexibility for using diverse glycans, other key beneficial bacteria such as butyrate producers primarily belonging to Firmicutes have a narrower range of glycan degrading capabilities^6,25^. Therefore, the release of PBPs by digestion of diverse glycans could promote species coexistence by the creation of new metabolic niches that can be exploited by other species including butyrate producers. However, we do not fully understand how glycan utilization via PULs in *Bacteroides* modulates inter-species interactions.

*Bacteroides uniformis* (BU) is one of the most abundant and prevalent species in the gut microbiome and is predicted to have 55 PULs^26-28^. However, we have limited understanding of the contributions of each PUL to the fitness of BU in response to diverse glycans. Equipped with a novel CRISPR genome editing system for *Bacteroides*, we investigate the contribution of 23 PULs in *B. uniformis* DSM 6597 using a pooled mutant barcoding strategy (**Fig. 1**). We discover glycan utilization functions of multiple PULs and the transcriptional coordination of PULs in response to the complex plant polysaccharide xyloglucan. Notably, while the presence of a given PUL can provide a fitness advantage in certain environments, we show that PULs can negatively impact fitness both *in vitro* and in the murine gut environment. These results provide key insights into the potential fitness benefit and cost of PULs in response to nutrient availability. Finally, we demonstrate that PULs in BU can shape ecological dynamics and the production of the beneficial metabolite butyrate in synthetic human gut communities through three major mechanisms. Due to the high abundance of BU in the human gut microbiome^26,27^, a deeper understanding of the molecular and ecological interactions mediated by PULs could inform precise microbiome interventions to benefit human health.

**Figure 1.**
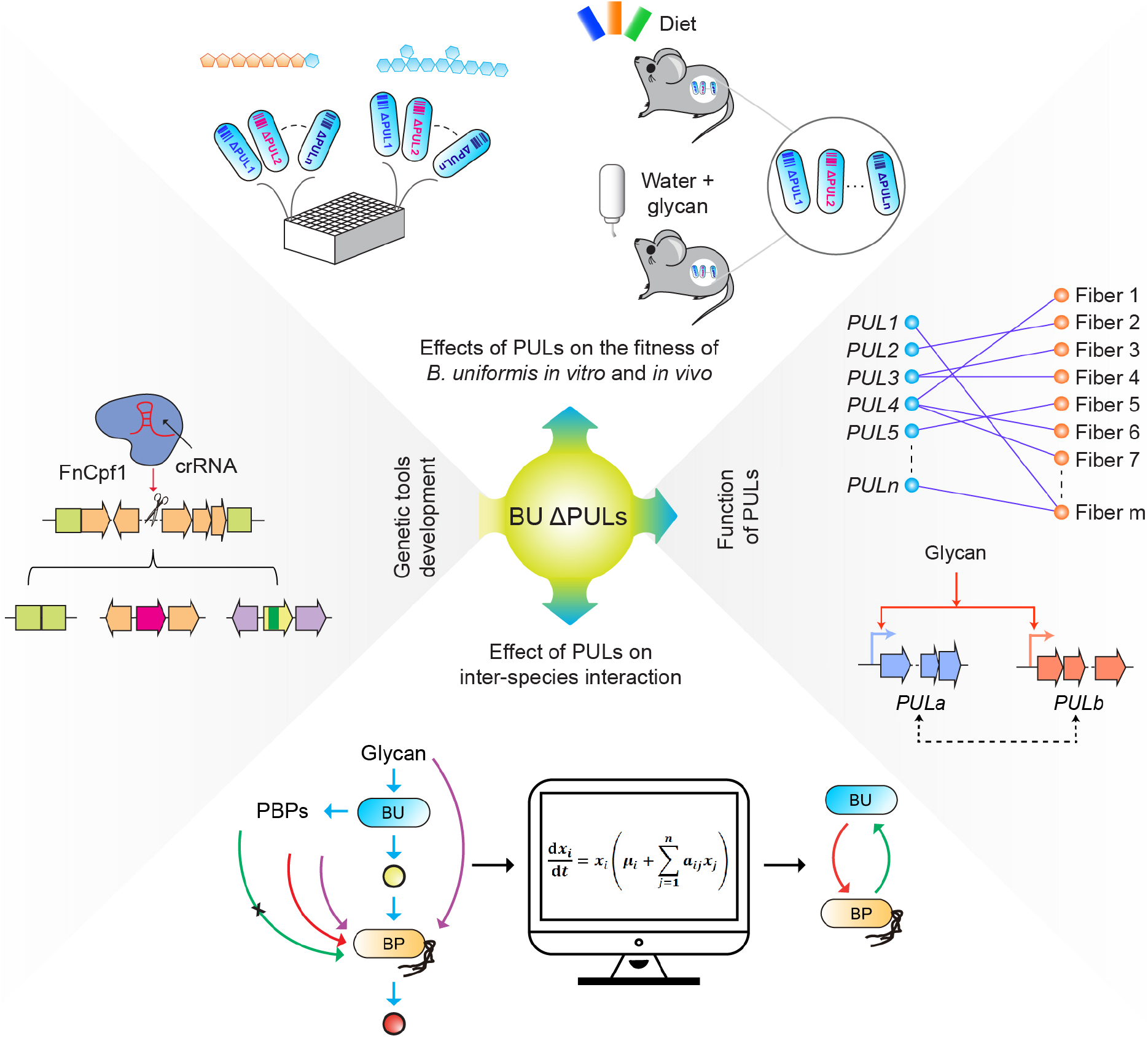
Effects of polysaccharide utilization loci (PULs) in *B. uniformis* (BU) on fitness and community-level interactions. Schematic highlighting the systematic characterization of 23 PULs in BU using a CRISPR-FnCpf1 genome editing tool. We investigated the contribution of PULs to the fitness of BU in media with a broad range of single glycans. We used transcriptional profiling to study the co-regulation between PULs in the presence of xyloglucan. We studied the impact of PULs on the fitness of BU *in vitro* and in germ-free mice in different nutrient environments. Finally, we examined the effects of PULs on community dynamics and butyrate production in the presence of diverse butyrate producers.

## RESULTS

### Development of genetic tools for construction of PUL mutants in BU

To investigate the contributions of PULs to fitness of BU, we developed new genome editing tool for *Bacteroides*. The current gene manipulation methods for *Bacteroides* are frequently based on two-step selections and counterselection^29-31^, thus limiting their generalizability to diverse *Bacteroides* isolates. The CRISPR-Cas system has been demonstrated as the most versatile genome editing tool thus far, and has been used for genome engineering of mammals, plants and diverse prokaryotes^32-34^. Developing CRISPR-Cas based genome editing tools in *Bacteroides* could expand our ability to understand and engineer health-relevant functions. Therefore, we sought to exploit CRISPR-Cas as a genome editing tool in *Bacteroides* that does not rely on a modified genetic background and genome-integrated selectable markers. To this end, we focused on the type V CRISPR-Cas protein from *Francisella novicida* U112 (FnCpf1) due to several unique features for genome editing, including its small size (i.e. lower fitness burden), endoribonuclease domain and functionality without requiring RNase activity^35^. To construct the CRISPR-FnCpf1 system, we characterized ribosome binding sites (RBSs), constitutive and inducible promoters and shuttle plasmids in BU^36-38^ (**Fig. S1, S2, Supplementary Note**).

Equipped with these tools, we demonstrated that the CRISPR-FnCpf1 *Bacteroides* genome editing system could successfully delete 23 PULs (efficiency of 3-100%) in BU that were selected based on predicted glycan degrading activities and cluster length (**Supplementary Data 1, Fig. S3, S4**). We found that expression of *E. coli* RecT improved the efficiency (∼3-fold higher in the presence of RecT) of the CRISPR-FnCpf1 mediated gene deletions in BU and the construction of ∼50% PUL mutants required RecT (**Fig. S3f**). We created two double deletions of *PUL22* and *PUL23* due to their proximity or *PUL11* and *PUL43* based on bioinformatic prediction of their functions^10^ (**Supplementary Data 1**). To quantify the abundance of each strain when combined into a single culture, we introduced a unique 4-base pair DNA barcode downstream of the *tyrP* locus (**Fig. 2a**). Finally, we demonstrated that the CRISPR-FnCpf1 method can be used to delete genes in diverse *Bacteroides* species and for gene insertions in BU (**Fig. S3, Supplementary Note**). In sum, these data show that CRISPR-FnCpf1 coupled with the expression of RecT can be used for efficient genome engineering in *Bacteroides*.

**Figure 2.**
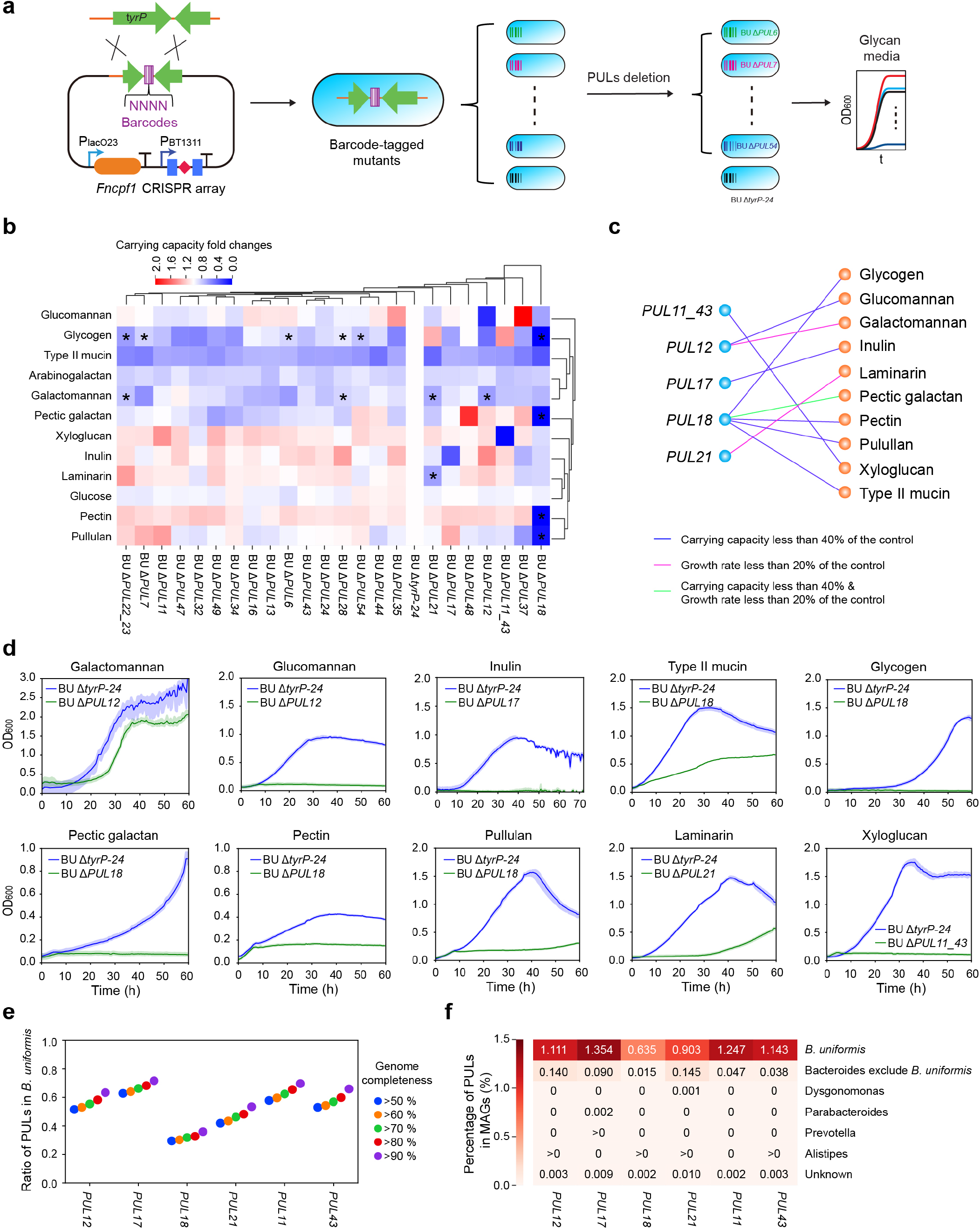
Effects of polysaccharide utilization loci (PULs) on *B. uniformis* (BU) monoculture growth in response to a range of glycans. **(a)** Schematic of the construction and characterization of barcoded BU mutants using the CRISPR-FnCpf1 genome editing tool. The growth of each mutant was characterized in media supplemented with diverse glycans. **(b)** Biclustering heatmap of the fold changes of inferred carry capacity based on a logistic growth model for each PUL mutant to Δ*tyrP-24* in media with single carbon sources. Asterisks represent carry capacities with coefficient of variation > 0.2, indicating low confidence in the inferred parameter value (**Supplementary Data 2**). In these conditions, the carry capacity was set to the maximum OD_600_. **(c)** Bipartite network of PULs and glycans generated based on thresholds in the fold changes of inferred carry capacity and/or growth rate. **(d)** Time-series measurements of OD_600_ of a set of PUL mutants in media with specific glycans based on the bipartite network in **(c)**. Lines denote the mean and the shaded regions represent 95% confidence interval of 3 biological replicates. **(e)** Categorial scatter plot of the ratio of PULs in BU metagenome-assembled genomes (MAGs) with varying genome completeness. **(f)** Heatmap of the percentage of PULs in MAGs with genome completeness >50%. The ‘>0’ denotes percentage of PULs greater than zero and less than 0.001.

### Effects of PUL deletions on the growth response of BU

BU can utilize diverse dietary and host-derived glycans in the human gut microbiome^4,9,14,17,39-41^. To determine how each PUL contributed to BU growth in environments with single glycans, we grew the BU wild-type strain (BU WT) in *Bacteroides* minimal media^42^ supplemented with 22 individual glycans, some of which were chosen based on previous studies^4,9,14,17,39-41^. Our results showed that 50% of the tested glycans could support the growth of BU (**Fig. S5**), indicating that BU provides key metabolic functions for the host by degrading a wide range of chemically diverse glycans.

To quantify the effects of glycans on the growth responses of PUL deletion mutants in different environments, we fit time-series measurements of absorbance at 600 nm (OD_600_) to a logistic growth model^43^. This model captures microbial growth as a function of exponential growth rate and carrying capacity (**Figs. 2b, S6, S7, Supplementary Data 2**). To determine potential effects of the genome integrated barcode on fitness, Δ*tyrP-24* (barcoded BU WT) was used as control. The growth responses of BU WT and Δ*tyrP-24* were similar in media supplemented with glucose, demonstrating that the barcode did not impact the growth response of BU (**Fig. S8**).

To determine how each PUL contributed to the fitness of BU in media with single glycans, we examined the fold change of the carrying capacity or growth rate for each PUL mutant compared to Δ*tyrP-24*. In the presence of glucose, the inferred growth parameters of the PUL mutants were similar to Δ*tyrP-24*, demonstrating that PUL deletions did not impact the growth of BU in media with simple carbon sources such as glucose (**Figs. 2b, S6**). However, deletion of the majority of PULs reduced the carrying capacity of BU in the presence of glycogen, type II mucin, arabinogalactan, or galactomannan. By contrast, the majority of PUL deletion mutants exhibited higher carrying capacities than Δ*tyrP-24* in xyloglucan, inulin, laminarin or pectin. The carrying capacity of Δ*PUL18* was the most significantly reduced across the majority of glycans, demonstrating that Δ*PUL18* contributed to the utilization of several glycans.

Based on these data, we highlighted a subset of PUL mutants that were determinants of BU fitness on a given glycan by applying a threshold in the percent change of the inferred growth parameters compared to Δ*tyrP-24* (<40% carrying capacity or <20% growth rate) (**Fig. 2c**). The bipartite network indicated that *PUL18* had multiple glycan degradation activities, *PUL12* contributes to glucomannan and galactomannan utilization, *PUL17* contributes to inulin utilization and both *PUL11* and *PUL43* contribute to growth on xyloglucan. These results were further validated using frequent time-series growth measurements (**Fig. 2d**). Whereas some mutants were unable to grow on a given glycan, the growth of other mutants were reduced (Δ*PUL12*-galactomannan, Δ*PUL18*-type II mucin and Δ*PUL21*-laminarin), suggesting that other genes could contribute to the utilization of these glycans.

The sequence and glucomannan/galactomannan utilization functions of *PUL12* have similarity to a previously reported PUL (*BACOVA_02087-97*) in *Bacteroides ovatus* (BO)^44^. While both PULs contain an outer membrane anchored glycoside hydrolase GH26, other glycoside hydrolases differed between *PUL12* and *BACOVA_02087-97*. Consistent with these differences, *PUL12* had greater specificity for glucomannan, whereas *BACOVA_02087-97* showed preference for galactomannan, indicating a potential trade-off in utilization of these chemically related glycans (**Fig. 2d**). The promiscuity of *PUL18* for starch analogues (glycogen and pullulan) and plant cell wall polysaccharides (pectin and pectic galactan) shares some similarity with previous reported *Bacteroides thetaiotaomicron* (BT) starch and pullulan utilization loci, as both PULs contain multiple GH13^30,45^. However, *PUL18* also contains an endo-β-1,4-galactanase GH53 (*BACUNI_RS06505*), which may enable *PUL18* to access multiple glycans (**Fig. 2d**). Finally, *PUL17* has the unique capability to utilize both inulin (β2-1 fructan) and levan (β2-6 fructan)^9,19^, which has not been previously observed in *Bacteroides* (**Fig. 2d, S9**). Based on protein sequence homology, predicted protein structure and sub-cellular localization, and previous studies, we propose biochemical models for glycan utilization (**Fig. S10-S13, Supplementary Note, Supplementary Data 1**)^28,46-50^.

To determine the abundance and prevalence of the PULs in the human gut microbiome, we performed bioinformatic analysis of two large metagenomic sequencing datasets^51,52^ (**Methods, Fig. 2e, 2f, S14**). Whereas *PUL12, 17, 21, 11* and *43* were found in over 53% of BU MAGs, *PUL18* was observed in ∼36%, indicating that *PUL18* was less conserved across BU strains than the other PULs (genome completeness >90%, **Fig. 2e**). Although these PULs were found in *Bacteroides* genomes beyond BU, they were infrequently observed in gut species excluding *Bacteroides* (**Fig. 2f**). In sum, these PULs were ubiquitous in BU but rarely found in other gut species, highlighting the unique role of BU in glycan utilization in the human gut microbiome.

### Xyloglucan utilization in BU is due to the coordination of PUL11 and PUL43

A xyloglucan utilization pathway (XyGULs) was previously reported in BO^10,53^, which shares high similarity to *PUL11* in BU. In addition, a second xyloglucan utilization pathway was predicted in BU^10^ (*PUL43* and *PUL44* in this work) (**Fig. 3a**). To investigate the functions of these PULs, we constructed double deletion strains Δ*PUL11_43* and Δ*PUL11_44*. We found that Δ*PUL11*, Δ*PUL43*, Δ*PUL44* and Δ*PUL11_44* were able to grow in the presence of xyloglucan, whereas Δ*PUL11_43* failed to grow. This implies that *PUL11* and *PUL43* have redundant roles in xyloglucan utilization (**Fig. 3b**). While Δ*PUL43* had a similar growth response to Δ*tyrP-24*, Δ*PUL11* and Δ*PUL11_44* exhibited a longer lag phase and higher carrying capacity than Δ*tyrP*-24 (**Fig. 3b**), suggesting that *PUL11* enables a faster response to xyloglucan but imposes a metabolic burden in stationary phase.

**Figure 3.**
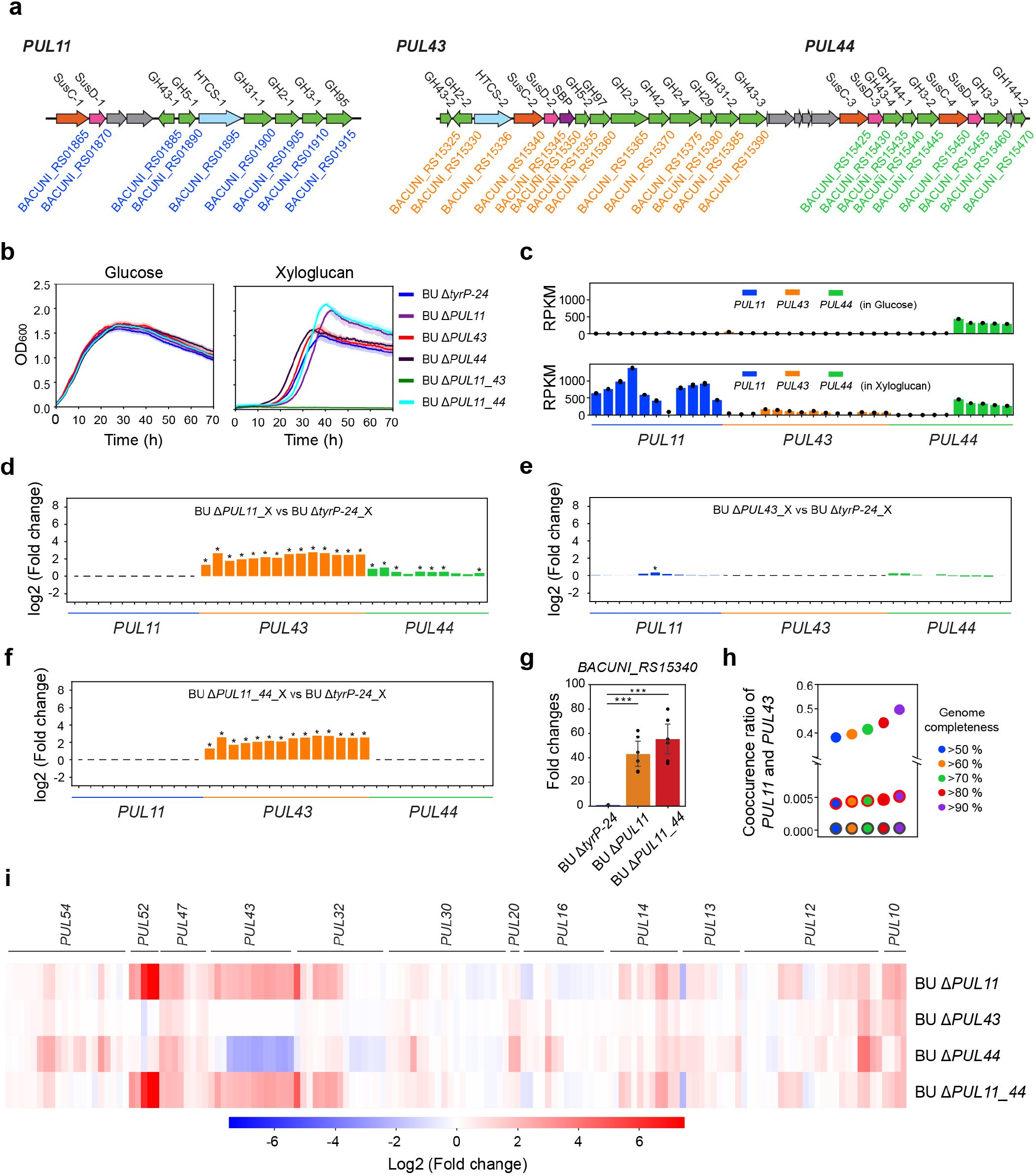
Co-regulation of polysaccharide utilization loci (PULs) for xyloglucan utilization in *B. uniformis* (BU). **(a)** Schematic of gene organization of *PUL11, PUL43* and *PUL44*. Colors represent predicted gene functions as described in **Fig. S16**. Genes with unknown functions are shaded in gray. GenBank locus tag numbers are shown for each gene. SusC: SusC-like TonB-dependent transporter; SusD: SusD-like cell-surface glycan-binding protein; SBP, sugar binding protein; HTCS, hybrid two-component system. **(b)** Time-series measurements of OD_600_ of PUL mutants and Δ*tyrP-24* in media with glucose (left) or xyloglucan (right). Lines denote the mean and shaded regions represent 95% confidence interval of 3 biological replicates. **(c)** Bar plots of the reads per kilobase per million mapped reads (RPKM) for each gene in *PUL11, PUL43* and *PUL44* in Δ*tyrP-24* in media with glucose or xyloglucan. Data points denote 2 biological replicates. The colored bars represent mean RPKM value of the genes shown in the same order as panel **(a)**. Bar plot of the log2 fold changes of RPKM of **(d)** Δ*PUL11* to Δ*tyrP-24*, **(e)** Δ*PUL43* to Δ*tyrP-24*, or **(f)** Δ*PUL11_44* to Δ*tyrP-24* in xyloglucan media (n=2, *p<0.05, unpaired t-test). **(g)** Bar plot of transcription fold changes of *BACUNI_RS15340* in *PUL43* to Δ*tyrP-24* in xyloglucan media using qRT-PCR (n = 6, ***p<0.001; Δ*PUL11* vs Δ*tyrP-24* p=2.9e-5; Δ*PUL11_44* vs Δ*tyrP-24* p=2.3e-5, unpaired t-test). All values indicate mean ± 1 s.d. **(h)** Categorial scatter plot of co-occurrence of *PUL11* and *PUL43* in metagenome-assembled genomes (MAGs). Data points outlined in black, red or without outlines indicate the ratio of co-occurrence in all MAGs, *Bacteroides* MAGs excluding BU or BU MAGs. **(i)** Heatmap of log2 fold changes of RPKM of PULs in PUL mutants that contained at least one gene displaying an absolute value of the log2 fold change greater than 2 compared to Δ*tyrP-24* in xyloglucan media.

We hypothesized that *PUL11* and *PUL43* could be transcriptionally co-regulated due to their redundant roles in xyloglucan utilization. Therefore, we performed genome-wide transcriptional profiling of Δ*tyrP-24*, Δ*PUL11*, Δ*PUL43*, Δ*PUL44* and Δ*PUL11_44* in the presence of xyloglucan or glucose as a control. The expression of genes in *PUL44* were not upregulated in xyloglucan compared to the glucose control, consistent with the negligible effect of the *PUL44* deletion on growth in media with xyloglucan (**Fig. 3b, c, S15**). Therefore, our results indicate that *PUL44* does not contribute to xyloglucan utilization and thus *PUL43* and *PUL44* have independent functions. In Δ*tyrP*-*24, PUL11* and *PUL43* were upregulated in media with xyloglucan compared to the glucose condition (**Fig. 2c, Fig. S15**), with *PUL11* exhibiting larger transcriptional fold changes on average than *PUL43*. The xyloglucan-dependent up-regulation of *PUL11* and *PUL43* corroborated their critical roles in xyloglucan utilization in BU (**Fig. 3b**).

To provide insight into potential transcriptional coordination, we evaluated how the expression of *PUL11* or *PUL43* were impacted by the deletions of *PUL43* or *PUL11* in media with xyloglucan, respectively. Notably, the expression of *PUL43* was significantly up-regulated in Δ*PUL11* and Δ*PUL11_44* compared to Δ*tyrP-24* in both the RNA-seq and qRT-PCR data, whereas the expression of *PUL11* did not depend on the presence of *PUL43* (**Fig. 3d-g, S15**). In sum, these data suggest that the regulatory coordination between *PUL11* and *PUL43* may enable BU to adapt to the loss of *PUL11* in xyloglucan (**Fig. 3b**). Guided by these data, we propose a biochemical model of xyloglucan utilization (**Fig. S16, Supplementary Note, Supplementary Data 1**).

We next examined the co-occurrence of *PUL11* and *PUL43* across BU MAGs in the human gut microbiome based on two metagenomic sequencing datasets^51,52^. We found that *PUL11* and *PUL43* were both present in ∼50% of BU MAGs (genome completeness >90%) and rarely found in MAGs from other gut organisms (**Fig. 3h**). The frequent co-occurrence of *PUL11* and *PUL43* in BU suggests that redundant PULs for xyloglucan utilization may be advantageous for BU.

### PULs can negatively or positively impact fitness in competition with other mutants

The presence of a given PUL can vary across isolates of the same gut species^54^ (**Fig. 2e**) and PULs have been shown to evolve within individuals, potentially due to the variable selection pressures of host diet^55^. Based on this observation, we hypothesized that PULs may provide a benefit or cost to microbial fitness depending on the nutritional landscape. To test this hypothesis, we combined the 22 PUL mutants and Δ*tyrP-24* into a single culture and characterized the mutant pool in the presence of single glycans (**Fig. 4a**). We determined the absolute abundance of each mutant based on relative abundance measured by barcode sequencing and OD_600_ (**Fig. 4a**). We characterized the fitness of the mutant pool in the presence of diverse glycans using serial dilution perturbations, which could represent variable transit and feeding patterns in the gut microbiome.

**Figure 4.**
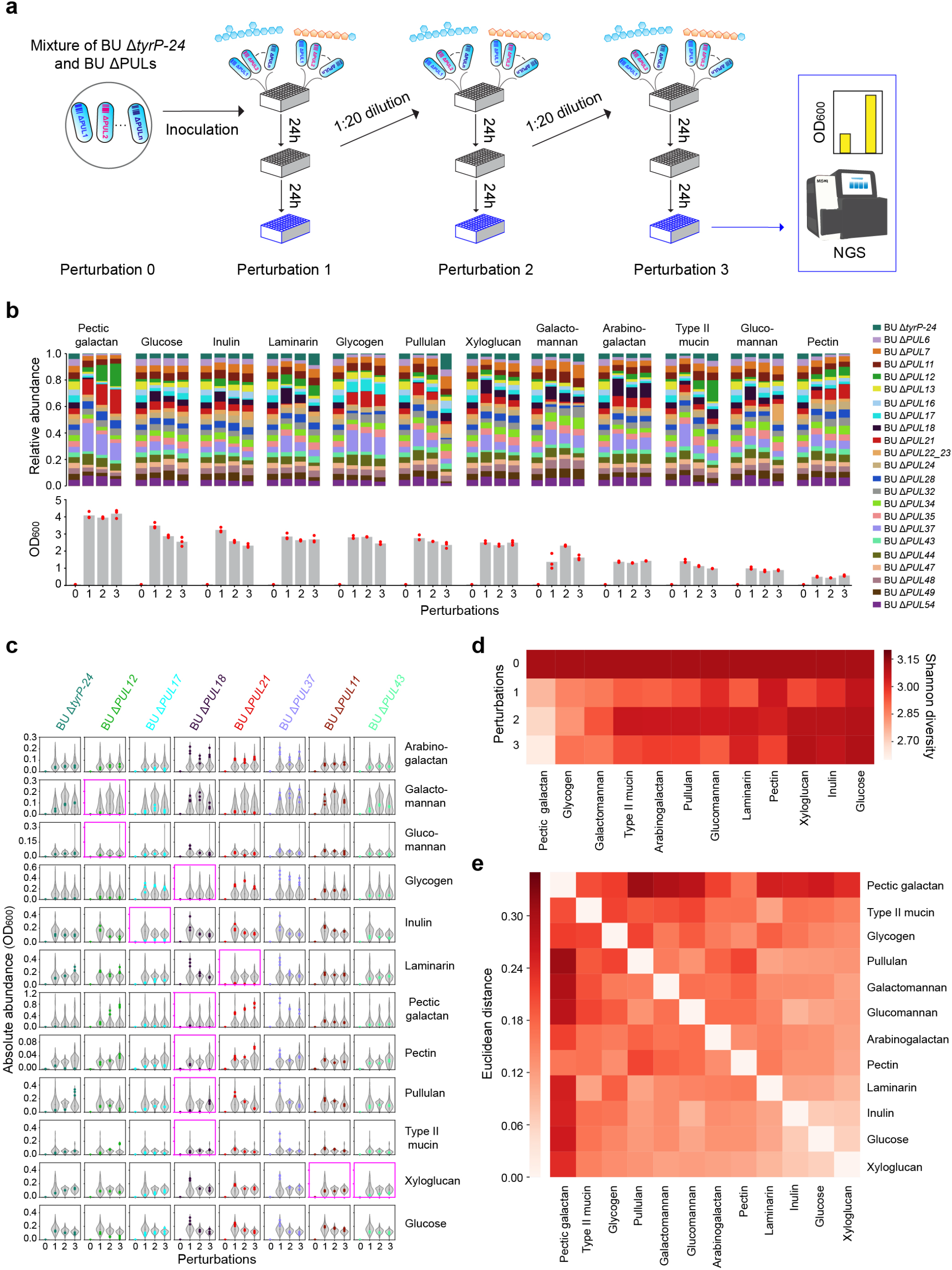
Impact of polysaccharide utilization loci (PULs) on *B. uniformis* (BU) pooled mutant fitness in media with different carbon sources. **(a)** Schematic representing experimental design of the pooled barcoded PUL mutants and Δ*tyrP-24* in media with different carbon sources. The mutant pool was passaged every 24 hr. The absolute abundance of each mutant was determined by next-generation sequencing and OD_600_ measurements. **(b)** Stacked bar plots represent the mean relative abundance of each mutant in media with a given carbon source (top). Bar plots denote the OD_600_ of each condition (bottom). Data points represent 3 biological replicates. **(c)** Violin plots represent the absolute mutant abundance in media with a given carbon source. Colored data points represent the absolute abundance of the indicated mutant (n = 3 biological replicates). The pink outlined subplots highlight the PUL-glycan pairs identified in **Fig. 2c. (d)** Heatmap of Shannon diversity of mutant pool across different perturbations. The values represent the mean of 3 biological replicates. **(e)** Heatmap of the Euclidean distance of the mutant pool composition (relative abundance) between different media for perturbation 3.

The OD_600_ of the mutant pool exhibited a wide variation across media, with pectic galactan and pectin exhibiting the highest and lowest total biomass across conditions, respectively (**Fig. 4b**). The growth impairment of certain PUL mutants (e.g. Δ*PUL12*-glucomannan or galactomannan; Δ*PUL17*-inulin, Δ*PUL18*-glycogen or pectic galactan or pectin; Δ*PUL21*-laminarin) were consistent both in monoculture and the mutant pool (highlighted subplots in **Fig. 4c**). However, there were also cases where the growth in monoculture and the mutant pool deviated in the presence of given glycan (e.g. Δ*PUL18*-type II mucin or pullulan) (**Fig. 4c**), suggesting that interactions between mutants (e.g. release of PBPs) could potentially rescue growth.

Our results showed that specific PUL deletions could enhance growth on a given glycan compared to Δ*tyrP-24* (e.g. Δ*PUL12*-laminarin or pectic galactan or pectin) (**Fig. 4c**). Furthermore, Δ*tyrP-24* was outcompeted by PUL mutants in media with glycogen, pectic galactan or pectin. These results indicate that PULs can provide a fitness cost in specific nutrient environments (**Fig. 4c, S17**). In addition, we found that Δ*PUL37* was highly abundant across many conditions, suggesting that *PUL37* was disadvantageous to BU (**Fig. 4c**). In sum, these results demonstrated that PULs can negatively impact fitness due to potential metabolic burden where the pathway is not required for growth^9,56^ (**Fig. 4c**).

We analyzed Shannon diversity of the mutant pool to provide insight into the strength of growth selection across different nutrient environments. The Shannon diversity in pectic galactan was substantially lower than in other glycan conditions, indicating pectic galatan provided a strong selection for the growth of certain BU mutants (**Fig. 4d**). By contrast, the Shannon diversity was high in glucose, inulin and xyloglucan, demonstrating that these carbon sources provided a weaker growth selection for BU mutants. The high Shannon diversity in these conditions was consistent with the robust monoculture growth of the majority of PUL mutants in media with glucose, inulin or xyloglucan (**Fig. 2b**).

To determine the impact of each glycan on the mutant pool composition, we examined the Euclidean distance of the relative abundance for Perturbation 3 (**Fig. 4e**). The mutant pool composition in pectic galactan exhibited the largest differences from the mutant pool compositions in other conditions, with the exception of pectin. These data suggest that the chemical similarity between pectic galactan and pectin selects for similar PUL mutant compositions (**Fig. 4e**). In addition, the mutant pool compositions were similar in the high Shannon diversity conditions (i.e. glucose, inulin and xyloglucan). In sum, our results demonstrated that PULs can have a positive or negative impact on fitness that depends on the nutrient environment, providing insight into the evolutionary selection for the presence or absence of PULs in the human gut microbiome.

### ΔPUL37 and ΔPUL18 exhibit high colonization ability in germ-free mice

To understand how PULs impact the fitness of BU in the mammalian gut environment in response to nutrient availability, we colonized gnotobiotic mice with the pooled PUL mutants and Δ*tyrP-24*, and fed groups of mice different diets that vary in the type and abundance of microbiota accessible carbohydrates (MAC) (**Supplementary Data 3**). Characterization of temporal changes in mutant abundance *in vivo* could provide insights into interactions between PULs in BU and the host as well as how the diet modulates these interactions. To this end, we colonized germ-free mice with the mutant pool on diet containing high MAC (high fiber diet) or MAC free (fiber free diet or FFD) (**Fig. 5a**). To understand the effects of specific diet-derived glycans on PUL mutant colonization, separate groups of mice fed the FFD received a given glycan (inulin, glucomannan or pectic galactan) in the drinking water (**Fig. 5a**). In addition, we included a high fat and low MAC diet (high fat diet) based on the hypothesis that BU would shift its metabolic niche towards utilization of host-derived glycans due to the limited availability of diet-derived glycans^57,58^.

**Figure 5.**
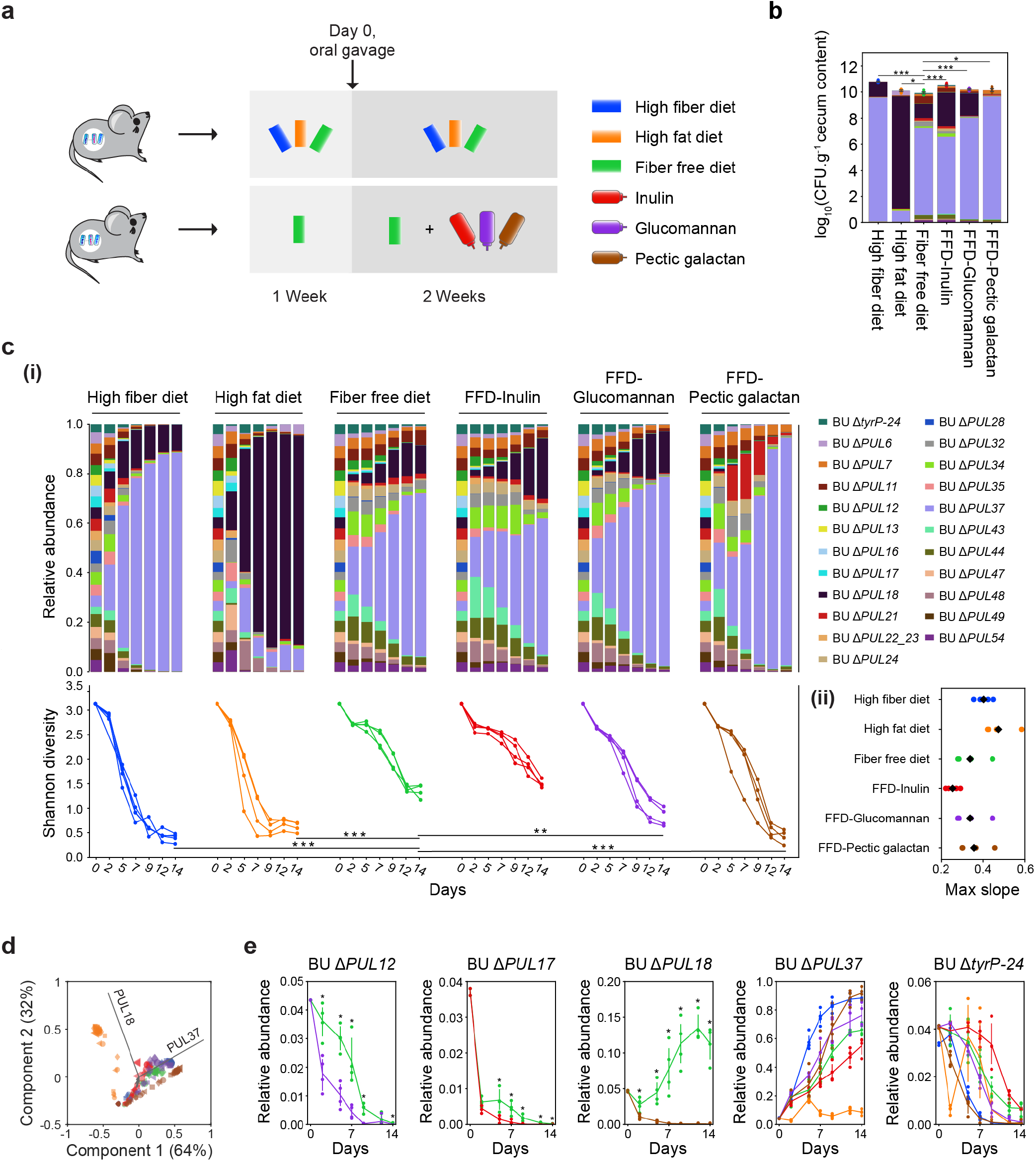
Impact of polysaccharide utilization loci (PULs) on the colonization ability of *B. uniformis* (BU) in germ-free mice fed different diets. **(a)** Schematic of experimental design to evaluate the effect of PULs on BU colonization ability in germ-free mice fed different diets. Top: mice were fed a high fiber diet, fiber free diet (FFD) or high fat diet a week prior to oral gavage and then maintained on the same diet for two weeks following oral gavage (n = 5 for high fiber group and n = 4 for other groups). Bottom: mice were fed the FFD a week prior to oral gavage and then provided with drinking water supplemented with inulin, pectic galactan or glucomannan (n = 4). On day 0, mice were orally gavaged with the BU mutant pool and Δ*tyrP-24*. Time-series measurements of fecal samples were performed. The cecal samples were collected at the end of the experiment. **(b)** Stacked bar plot of the absolute abundance of mutants in cecal samples (CFU g^-1^) in each group of mice. Data points denote 2 independent CFU measurements for each mouse. Asterisks denote a statistically significant difference in the CFU of each group compared to FFD group based on unpaired t-test (n = 8; *p<0.05, ***p<0.001; High fiber diet vs FFD, p=4.9e-9; High fat diet vs FFD, p=0.01271; FFD-Inulin vs FFD, p=2.8e-6; FFD-Glucomannan vs FFD, p=1.0e-4; FFD-Pectic galactan vs FFD, p=0.01393). **(c)** (i) Stacked bar plots of relative abundance of mutants in fecal samples as a function of time (top). Line plots of Shannon diversity of the mutant pool as a function of time (bottom). Asterisks indicate statistically significant difference of Shannon diversity of each group on day 14 compared to FFD group based on unpaired t-test (n=4-5; **p<0.01, ***p<0.001; High fiber diet vs FFD, p=3.9e-6; High fat diet vs FFD, p=1.4e-4; FFD-Glucomannan vs FFD, p=0.00425; FFD-Pectic galactan vs FFD, p=7.9e-5). (ii) Categorial scatter plot of the maximum slope of the Shannon diversity as a function of time. The colored data points represent individual mice and the black data point represents the mean. **(d)** Principal component analysis (PCA) as a function of time. Colors represent different diets based on the legend in **(a)**. The size of the data points is proportional to the time of measurement. The PCA loadings are denoted by the black lines. Symbols represent different mice. **(e)** Line plots of the relative abundance of PUL mutants or Δ*tyrP-24* in mice fed different diets as a function of time. The colors represent different diets based on the legend in **(a)**. Data points denote individual mice and lines represent the mean. The asterisks denote a statistically significant difference based on unpaired t-test (n=4-5, p<0.05).

To understand how variation in the diet impacted the overall fitness of the mutant pool in the murine gut, we characterized the absolute abundance of BU in the cecum at the end of the experiment. The colony forming units (CFU) g^-1^ of BU was significantly higher in all diets than the group fed with FFD and the CFU g^-1^ was highest in the high fiber diet (**Fig. 5b**), indicating that higher MAC enhanced the colonization ability of BU. In addition, *ΔPUL37* was high abundance in all conditions except the high fat diet whereas *ΔPUL18* was present at variable levels in all diets except FFD-pectin galactan. The temporal changes of the composition of the mutant pool varied across diets, indicating that the nutrients in the diet are a critical variable shaping the colonization ability of PUL mutants (**Fig. 5c**). We analyzed the maximum rate of change of Shannon diversity to quantify the strength of selection for colonization of specific PUL mutants across diets. The Shannon diversity of the mutant pool decreased most rapidly in the high fiber and high fat diets, whereas a gradual decrease in diversity was observed in the FFD and FFD supplemented with inulin (FFD-inulin). After the two-week period, the Shannon diversity was low in the high fiber, high fat diet and FFD supplemented with pectic galactan (FFD-pectic galactan), and converged to a higher steady-state value for mice fed FFD and FFD-inulin. Notably, the trends in Shannon diversity *in vivo* mirrored the high and low Shannon diversities in the *in vitro* experiment, with the diversity being lowest and highest in media with pectic galactan or inulin/glucose, respectively (**Figs. 4d, 5c**). These results suggest that diet composition was one of the major driving factors of Shannon diversity *in vivo*. Principal component analysis over all time points revealed that Δ*PUL18* and Δ*PUL37* contributed most significantly to the temporal changes in the variance of the mutant pool composition and distinguished the high fat diet from all other diets (**Fig. 5d**).

The abundance of most mutants decreased over time in FFD, FFD-inulin and FFD-glucomannan groups, potentially due to competition with the high fitness mutants Δ*PUL18* and Δ*PUL37* (**Figs. 5c, e**). However, Δ*PUL12* was higher abundance in FFD than FFD-glucomannan for a period of time and Δ*PUL17* exhibited moderately higher abundance in FFD than FFD-inulin, consistent with the *in vitro* characterization of these strains (**Figs. 2d, 5e**). Δ*PUL18* was unable to colonize mice in FFD-pectic galactan but persisted as a high fraction of the community in the mice fed all other diets, consistent with the critical role of this PUL in pectic galactan utilization identified *in vitro* (**Figs. 2d, 5e**). Together, these data demonstrate that the absence of a given PUL required for utilization of given glycan reduced the colonization ability of BU in response to a diet containing this glycan. Therefore, the glycan-utilization functions of PULs identified *in vitro* can be used to predict the colonization ability of BU in the mammalian gut.

The deletion of *PUL37* enhanced colonization ability across most diets, enabling this mutant to dominate the mutant pool. This is consistent with its high *in vitro* fitness in response to a wide range of glycans and demonstrates that the loss of *PUL37* provides a substantial fitness advantage to BU (**Figs. 4c, 5c**). In addition, Δ*PUL18* was able to stably colonize the murine gut in all diets except FFD-pectic galactan and dominated the community in the high fat diet (**Fig. 5c**). This suggests that the high colonization ability of Δ*PUL18* is dependent at least in part on the availability of diet-derived or host-derived nutrients in response to the high fat diet. Notably, Δ*tyrP-24* only colonized mice fed the FFD and FFD-inulin diets at a low abundance. The diminished colonization ability of Δ*tyrP-24* in competition with other mutants mirrored the low fitness of Δ*tyrP-24 in vitro* (**Fig. 4c, S17**). Therefore, the colonization ability of BU *in vivo* could be substantially enhanced by the loss of specific PULs (**Fig. 5e**).

Overall, our data show that the effects of PULs on BU colonization are complex: the presence of a given PUL could improve fitness in specific nutrient conditions, whereas it can also reduce the fitness in other conditions. We found that certain qualitative trends observed in our *in vitro* experiments could predict the qualitative trends in mice fed diets containing similar nutrient compositions, highlighting that diet-derived nutrient availability is a major factor shaping the colonization ability of the BU mutants. In addition, the rapid loss of diversity in the composition of mutant pool suggests a high strength of selection of PULs in the murine gut environment. Therefore, our data suggest that the presence of PULs is a key factor determining the ability of BU to colonize mice due to an interplay of nutrient availability and potential microbe-host interactions.

### PULs are major drivers of inter-species interactions with butyrate producers

Butyrate produced by a specialized group of gut bacteria is linked to numerous health benefits including maintaining homeostasis of the gut environment^59-61^. We hypothesized that glycans utilized by BU could impact the ecological dynamics of gut communities including the abundance of butyrate producers and thus butyrate production (**Fig. S5**). To this end, we characterized the growth of BU WT, Δ*PUL12*, Δ*PUL17*, Δ*PUL18*, Δ*PUL21* and Δ*PUL11_43* and four highly prevalent butyrate producers in the human gut microbiome, including *Anaerostipes caccae* (AC), *Coprococcus comes* (CC), *Eubacterium rectale* (ER) and *Roseburia intestinalis* (RI), in monoculture and in pairwise communities in media with single carbon sources^62^ (**Fig. 6a**).

**Figure 6.**
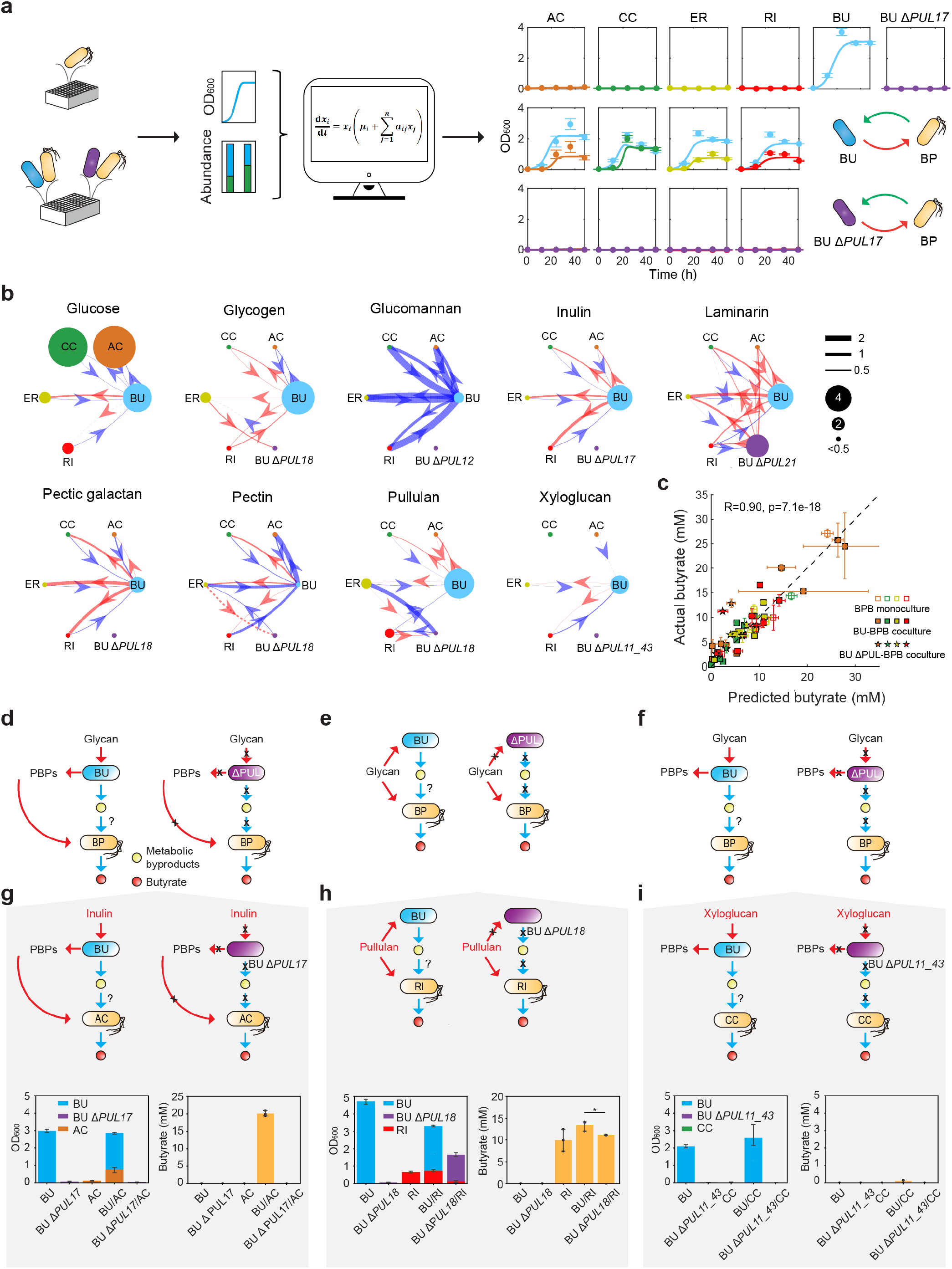
*B. uniformis* (BU) polysaccharide utilization loci (PULs) mediate glycan-dependent inter-species interactions influencing butyrate production. **(a)** Schematic representing the experimental design investigating PUL mediated inter-species interactions between BU and butyrate producers *A. caccae* (AC), *C. comes* (CC), *E. rectale* (ER) and *R. intestinalis* (RI). Inter-species interactions were deduced using a generalized Lotka-Volterra (gLV) model informed by time-series data of species absolute abundance. Representative data (right) shows time-series measurements of absolute abundance based on 16S rDNA sequencing and OD_600_ measurements in inulin media (**Fig. S18**). **(b)** Inferred networks of inter-species interactions between BU wild-type or a given PUL mutant and each butyrate producer in media with different carbon sources. Node size represents the maximum mean OD_600_ measured in monoculture in the indicated media. For species whose maximum mean OD_600_ is less than 0.5, their node sizes were set to OD_600_=0.5 for visibility in the network. The width of an edge connecting node *j* to node *i* represents the magnitude of the median of the inferred marginal distribution of their inter-species interaction coefficient (*a*_*ij*_). An edge is colored red (blue) if the interaction is positive (negative). If the 25% and the 75% quantiles of the *a*_*ij*_ marginal distribution have different signs, we represent the edge with a dashed line, indicating lack of certainty. Inter-species interactions where the magnitude of the median of the marginal *a*_*ij*_ distribution is less than 0.01 are not included in the network. **(c)** Scatter plot of predicted and measured butyrate concentrations in monoculture and coculture experiments. Predicted butyrate concentrations are computed according to the linear regression model (**Methods)**. Marker horizontal position represents predicted butyrate concentration based on mean end point butyrate producer abundance, and horizontal error bars represent 1 s.d. of predicted butyrate concentration given the uncertainty in butyrate producer abundance measurements. Schematic of proposed mechanism of PUL mediated interactions between BU and butyrate producers (**d-f**). **(d)** In Mechanism A, the butyrate producer is unable to utilize the given glycan but can utilize PBPs released by BU. **(e)** In Mechanism B, the butyrate producer can utilize the glycan and thus compete with BU. **(f)** In Mechanism C, butyrate producer lacks the capability to utilize both the given glycan and PBPs potentially released by BU. Representative data of **(g)** BU-AC in inulin media consistent with Mechanism A, **(h)** BU-RI in pullulan media consistent with Mechanism B and **(i)** BU-CC in xyloglucan media consistent with Mechanism C. Stacked bar plot of the absolute abundance of each strain in monoculture and co-culture (left). Bar plot denotes the butyrate concentration in each condition (right). All values are mean ± 1 s.d (n=3 biological replicates). The asterisk denotes a statistically significant difference based on unpaired t-test (p=0.031).

To decipher inter-species interactions, we fit a generalized Lotka-Volterra model (gLV) to time-series measurements of species absolute abundance based on 16S rDNA sequencing and OD_600_ (**Fig. 6a, S18, Supplementary Data 4**). The gLV model is a set of coupled differential equations that describe the growth dynamics due to each organism’s intrinsic growth rate and interactions with each community member^43^. We used a Markov-Chain Monte Carlo method to infer the parameters based on the data (**Fig. S18-S20**).

Visualizing the inferred inter-species interaction coefficients as a network highlighted that BU WT can substantially enhance the growth of butyrate producers in media with inulin, laminarin, pectic galactan, pectin or pullulan (**Fig. 6b, S18**). Many of these inferred interactions vanished in pairwise communities composed of a PUL mutant and butyrate producer, demonstrating the critical role of PULs in mediating inter-species interactions. For example, BU and each butyrate producer co-existed in media with inulin, but the growth of all butyrate producers was abolished in co-culture with Δ*PUL17* (**Fig. 6a, b, S18**). In cases where the butyrate producer could utilize the glycan, the inferred inter-species interactions with BU WT or the PUL mutant exhibited major differences in directionality and sign. For instance, ER could utilize pullulan and its growth was enhanced by BU WT and inhibited by Δ*PUL18* (**Fig. 6b, Fig. S18a**). Further, the growth of BU WT was inhibited by ER whereas the growth of Δ*PUL18* was enhanced by ER. Together, these data demonstrate the key role of PULs in shaping ecological networks.

Previous work has demonstrated that the abundance of butyrate producers can be used to predict butyrate production^62^. Based on this result, we constructed a linear regression model to predict the end-point butyrate concentration based on the abundance of butyrate producers (**Methods**). Using the trained regression models, the predicted butyrate concentration was highly correlated to the measured butyrate concentration (Pearson r=0.9, p=7.1e-18) (**Fig. 6c**). Therefore, these results demonstrate PUL-dependent BU glycan utilization impacts butyrate production via modulation of the growth of butyrate producers as opposed to growth uncoupled activities^62^. In sum, these data demonstrate the critical role of BU PULs in mediating inter-species interactions influencing butyrate production, and therefore could be potential engineering targets for controlling butyrate production in the human gut microbiome.

### PULs influence BU-butyrate producer interactions via three major mechanisms

We next sought to understand the molecular basis of the inferred inter-species interactions between BU and butyrate producers. We hypothesized that the inferred positive interactions supporting the growth of the butyrate producers in co-culture with the BU WT could be due to the release of metabolic byproducts or energy rich PBPs.

To determine if the released compounds from BU could impact butyrate producer growth, we characterized the growth of butyrate producers in BU conditioned glycan media. To eliminate the effect of environmental pH modification, the pH of the conditioned glycan media was adjusted to match fresh media (**Fig. S21**). By evaluating the ratio of total growth (area under the curve or AUC) of each butyrate producer in conditioned glycan media (AUC_CM_) to the corresponding fresh media (AUC_FM_), we found that the conditioned glycan media enhanced the growth of butyrate producers across many conditions. For instance, a growth benefit for all butyrate producers was observed in conditioned inulin or laminarin media, whereas a negligible growth benefit was observed for other conditions such as CC in conditioned xyloglucan media (**Fig. S21, Table S3**). Therefore, our results demonstrate that the effect of released compounds from BU on the growth of each butyrate producer was determined by the specific glycan, consistent with the glycan-dependent variation in the inferred inter-species interaction networks for the co-cultures (**Fig. 6b**).

We next investigated the effects of metabolic byproducts excluding PBPs on the growth of butyrate producers. Assuming that glucose was the limiting resource, we characterized the growth of butyrate producers in BU conditioned glucose media, where the glucose concentration and pH were adjusted to match the values of fresh media (**Fig. S22**). We observed a moderate increase in the growth of the butyrate producers (1.2-2.2-fold) in the conditioned glucose media compared to fresh media (**Fig. S22c, Table S3**), suggesting that metabolic byproducts can provide only a minor growth benefit for butyrate producers.

As *Bacteroides* have been shown to release acetate, propionate and succinate as metabolic byproducts^60^, we tested whether these molecules can support the growth of the butyrate producers. Butyrate producer growth was not observed in media with acetate, propionate and succinate as the primary carbon sources or in media with single glycans (**Fig. S23, S24**). Together, these results indicate that acetate, propionate and succinate released from BU do not support the growth of the butyrate producers.

Since our results suggested that metabolic byproducts released by BU did not provide a substantial growth benefit for the butyrate producers, we next characterized the effects of PBPs released by BU due to outer-membrane glycan degrading enzymes (**Fig. 6b, S10-13, S16, S18**)^9,14,23^. To test this possibility, we treated glycan media with BU cell membrane fractions (**Methods, Fig. S25a**) and determined the fold change in the AUC of the butyrate producer growth response in the cell membrane treated (AUC_MT_) to untreated media (AUC_FM_) (**Fig. S25, Table S3**). Our results suggest that BU releases PBPs that can be utilized by a butyrate producer if AUC_MT_ AUC_FM_^-1^ was much larger than 1 and the associated PULs were predicted to have only outer-membrane anchoring glycan degrading enzymes (e.g. *PUL17*-inulin) (**Fig. S10-13, S16, S25, Table S3**).

We used the combined results of these experiments to predict whether the PUL mechanism was selfish or unselfish (i.e. released PBPs). For instance, for inulin and laminarin, all butyrate producers displayed substantial growth benefits in co-culture with BU, BU conditioned glycan media and cell membrane treated glycan media compared to their respective controls. This suggests that *PUL17* and *PUL21* degrade their respective glycan via unselfish mechanisms (**Fig. S18, S21, S25, Table S3**). In addition, we observed growth enhancements in these experiments for certain fibers degraded by *PUL18* for specific butyrate producers, suggesting an unselfish mechanism of degradation of these glycans by *PUL18* (e.g AC or CC in pullulan; ER or RI in pectic galactan). For xyloglucan, we observed a growth enhancement of specific butyrate producers (e.g. AC, ER or RI) in BU conditioned xyloglucan media and in cell membrane treated xyloglucan media, but not in co-culture with BU (**Table S3**). These data suggest that BU can release xyloglucan breakdown products and AC, ER and RI have the capability to utilize these products, but the community interactions are more complex.

Based on our data, we propose three classes of mechanisms where PULs in BU can influence inter-species interactions (**Fig. 6d, e, f**). In Mechanism A (**Fig. 6d**), the butyrate producer can utilize PBPs released by BU but not the glycan. In this case, the growth of butyrate producer would be enhanced in co-culture with BU WT but not supported in co-culture with the PUL deletion mutant required for utilization of the given glycan (e.g. AC and BU in inulin media) (**Fig. 6g**). Whereas AC did not display growth or butyrate production in inulin media or in co-culture with Δ*PUL17*, significant growth and butyrate production were observed in the presence of BU WT (20.1 mM butyrate) (**Fig. 6g**). Consistent with Mechanism A, fructose was detected in BU conditioned inulin media and growth of AC was observed in media with fructose as the primary carbon source, indicating that the growth of AC was enhanced by breakdown products of inulin released by BU (**Fig. S26**).

In Mechanism B, BU and the butyrate producer can both utilize a given glycan and thus the butyrate producer exhibits growth in co-culture with BU WT or the PUL deletion mutant (**Fig. 6e**). However, the presence or absence of a PUL required for utilization of a certain glycan can shape the inter-species interaction network (**Fig. S18**). For example, RI displayed growth in pullulan media as well as in co-culture with BU WT or Δ*PUL18* (**Fig. 6h**). The growth of BU WT was inhibited by RI whereas RI enhanced the growth of *ΔPUL18*, which was captured by the inferred inter-species interaction network (**Fig. 6b, S18b, c**).

In Mechanism C (**Fig. 6f**), the butyrate producer lacks the capability to utilize both the glycan and corresponding PBPs. In this case, low energy metabolic byproducts released by BU could lead to a moderate or even negligible growth benefit (e.g. BU and CC in xyloglucan media) (**Fig. 6i**). CC did not exhibit significant growth or butyrate production in co-culture with BU (0.1 mM butyrate) or Δ*PUL11_43*, indicating that the released compounds from BU did not provide a major benefit to CC. In addition, the BU conditioned xyloglucan media or BU cell membrane treated xyloglucan media did not enhance the growth of CC (**Fig. S21, S25, Table S3**). Therefore, the specific mechanism of PUL degradation and glycan utilizing capabilities of the butyrate producer are critical variables shaping community-level interactions and dynamics.

## DISCUSSION

We developed a novel CRISPR-FnCpf1 genome editing tool to comprehensively study the role of 23 PULs on the fitness of BU in different environments, providing a comprehensive understanding of how diverse PULs contribute to glycan utilization, microbial interactions and colonization of the murine gut environment in response to nutrient availability. We show that PULs can provide a fitness benefit or cost (e.g. deletion of *PUL37* or *PUL18* enhanced the colonization ability of BU in the murine gut environment) depending on the nutrient landscape. We show that deletion of a given PUL impacts the gene expression patterns of other PULs across the BU genome (**Fig. 3i**), highlighting unknown mechanisms for transcriptional coordination. In addition, PULs can shape ecological networks via competition for glycans or release of PBPs, thus shaping the butyrate production capability of gut communities.

Deletion of the more highly expressed *PUL11* for xyloglucan utilization resulted in significant up-regulation of the second xyloglucan utilization pathway *PUL43*, but not the reciprocal. This observation suggests that the population compensates for the absence of *PUL11* by up-regulating *PUL43*, allowing the population to adapt to the loss of a major PUL for xyloglucan utilization (**Fig. 3c-3g**). This transcriptional coordination could be due to cross-regulation of *PUL43* by a sensor-regulator in *PUL11*, mirroring the transcriptional linkage observed between two xylan-degrading PULs in *Bacteroides xylanisolvens*^63^. Notably, we showed that PUL deletions can lead to major shifts in the gene expression patterns of other PULs distributed across the BU genome (**Fig. 3i**). The shifts in the gene expression patterns of PUL genes across the BU genome in response to a given PUL deletion could also arise due to cross-regulation of a sensor-regulator in *PUL11* or *PUL43*. Alternatively, differences in the composition of enzymes in *PUL11* and *PUL43* could potentially lead to variation in PBPs, which in turn could alter the activities of sensor-regulators that respond to similar PBPs in disparate PULs. Finally, global regulators that respond to carbon limitation could couple the expression of disparate PULs^64,65^.

In competition with other mutants *in vitro* and *in vivo*, BU strains that lacked specific PULs displayed enhanced fitness compared to the control strain that harbored all PULs. Therefore, while a given PUL can enhance fitness by enabling utilization of key nutrients, expression of PUL genes may impose substantial energetic costs when they are not needed for growth^66^. Supporting this hypothesis, BT has been shown to constitutively express certain PULs at a low-level^9,56,67^. This regulatory strategy may allow the cells to rapidly adapt to temporally changing nutrient conditions at the cost of unnecessary gene expression.

In the murine gut, Δ*PUL18* and Δ*PUL37* colonized the murine gut at a high level in all diets except for Δ*PUL18* in FFD-pectic galactan, due to the pectic-galactan degrading function of Δ*PUL18* (**Fig. 5c**). Deletion of *PUL18* resulted in a decrease in fitness in the presence of type II mucin *in vitro* (**Fig. 2b, d**), suggesting that *PUL18* can also contribute to the utilization of host-derived polysaccharides. The human hydrolases glucosylceramidase, sialidase and hexosaminidase target degradation of the carbohydrate head group of glycosphingolipids^68^. Bioinformatic analysis of *PUL37* revealed glucosylceramidases (*BACUNI_RS00220* and *BACUNI_RS00255*), an exo-α-sialidase (*BACUNI_RS00240*) and a β-hexosaminidase (*BACUNI_RS00225*), suggesting that *PUL37* could potentially contribute to the degradation of glycosphingolipids on host cells (**Supplementary Data 1**). Therefore, *PUL18* and *PUL37* may share a common function in utilization of host-derived polysaccharides. Immunoglobulin A (IgA) released in the gut regulates the growth and functional activities of gut microbiota, which was recently shown to target pectin and fructan utilization PULs in BT, leading to a decrease in PUL gene expression^69^. Based on the hypothesis that *PUL18* and *PUL37* may degrade host-derived glycans, IgA released by the host could regulate these activities by targeting proteins in *PUL18* and/or *PUL37*. Therefore, the loss of *PUL18* or *PUL37* could in turn promote the colonization ability of BU by eliminating interactions with IgA. However, we cannot exclude other possibilities including the role of *PUL18* or *PUL37* as receptors for prophage^70^ or their interactions with capsular polysaccharide synthesis, which provides protection from the host immune system and phage^56,67^. Therefore, the two PULs especially *PUL37* could be exploited as a potential engineering target for manipulating the colonization ability of BU in the mammalian gut.

BU enabled the growth of diverse butyrate producers in media with inulin, laminarin, pectic galactan, pectin and pullulan, suggesting that *PUL17, PUL21* and *PUL18* degrade certain glycans via unselfish mechanisms by releasing PBPs. Corroborating the key role of *Bacteroides* in modulating butyrate production, BT has been shown to enhance the growth and butyrate production of *E. ramulus* via starch breakdown products^71^, AC via human milk carbohydrate breakdown products^72^, and up-regulate the host butyrate transporter *Mct-1* in the presence of ER in the murine gut^58^. However, we found that interactions between BU and butyrate producers can also be more nuanced beyond the release of PBPs. For example, growth of AC, ER and RI were enhanced in BU conditioned xyloglucan media and cell membrane treated xyloglucan media but not in co-culture with BU in xyloglucan media. These data suggest that BU can degrade xyloglucan using an unselfish mechanism in monoculture and may either switch to a selfish xyloglucan-degrading mechanism in co-culture and/or release other compounds that inhibit butyrate producer growth. Further, we found that complex interactions occurred between BU and butyrate producers in cases where both species can utilize a certain glycan. For example, both species could compete for the available glycan and the butyrate producer could enhance the growth of BU via release of PBPs. In sum, these data demonstrate that inter-species interactions between *Bacteroides* and diverse butyrate producers, mediated by glycan competition and PBPs release, are critical determinants of ecological dynamics. Metabolic complementarity between Bacteroidetes and Firmicutes may promote coexistence and stability in the human gut microbiome.

PULs harbored by gut bacteria provide the host with the unique capability to transform chemically diverse glycans into molecules that serve as nutrients for the host and shape the ecological dynamics of the gut microbiome. Future work will investigate the predicted biochemical mechanisms of glycan degradation of the PULs identified in this study. To provide insights into the mechanisms of niche differentiation in the murine gut in response to nutrient availability, we could explore the impact of other gut species on the temporal changes in abundance of PUL mutants. A deeper understanding of the contributions of PULs to microbial fitness and community-level functions could guide the design of microbiome interventions to optimize nutrient extraction from food or restore homeostasis in the gut following a disturbance. For example, PULs could be harnessed as a control knob to modulate key inter-species interactions to enhance the production of beneficial metabolites produced by gut microbiota. Identifying such mechanistic control parameters that influence community-level functions performed by gut microbiota will enable us to harness the potential of the system to benefit human health.

## METHODS

### Microbial strains and growth conditions

Detailed information of the strains used in this study is provided in **Table S1**. All anaerobes were cultured in an anaerobic chamber with an atmosphere of 83% N_2_, 2% H_2_ and 15% CO_2_ at 37 °C. For most experiments, *Bacteroides* strains, AC, CC were grown in Anaerobe Basal Broth (ABB, Oxoid), ER was grown in ABB media with the addition of 3.3 mM sodium acetate (Sigma) and RI was grown in Brain Heart Infusion Broth (BHI, Sigma). We used *E. coli* pir2 (Invitrogen) for cloning and maintenance of plasmids with R6K origin (pNBU2-ermGb derivatives and pFW1000 derivatives, **Table S2**). *E. coli* DH5α (Thermo Fisher Scientific) was used for cloning and maintenance of plasmids with p15A, pSC101ts and ColE1 origins (pFW2000 derivatives, pFW3000 and pFW4000). We used *E. coli* BW29427 (*E. coli* Genetic Stock Center, CGSC) for *E. coli*-*Bacteroides* conjugations. All *E. coli* strains were grown aerobically in Luria Bertoni (LB) media. To support the growth of *E. coli* BW29427, we supplemented LB media with 25 nM of 2,6-Diaminopimelic acid (DAP, Sigma). We used the following antibiotics when required including 100 µg mL^-1^ carbenicillin (Carb, IBI Scientific), 25 µg mL^-1^ erythromycin (Em, Sigma) and 200 µg mL^-1^ gentamicin (Gm, Sigma). We used 1 mM of Isopropyl β-D-1-thiogalactopyranoside (IPTG, Gold Biotechnology) for the induction of FnCpf1.

### Plasmid construction

All plasmids used in this work are described in **Table S2** and sequences for genetic elements are in **Supplementary Data 5**. The P_BfP1E6_ promoter fused to 18 previously reported RBSs were cloned into pNBU2-ermGb at the Not I restriction site to characterize the strength of each RBS in BU^37,30,73^. We constructed plasmids with 20 BU native promoters as well as three previously reported strong promoters in BT (P_BT1311_, P_cfiA_ and P_BfP1E6_) by cloning each promoter into pNBU2-ermGb using the Not I restriction site. In addition, RBS8 was fused to the promoters P_BU18065_, P_BU18270_, P_BU15675_ and P_BfP1E6_. To construct the shuttle plasmid pFW1000, the *Bacteroides* replication origin PB8-51^74^ (synthesized by Twist Biosciences) was cloned using the Nsi I and Kpn I restriction sites in pNBU2-ermGb by replacing the NBU2 integrase gene and *attN2* site. To generate the shuttle plasmids pFW2000, pFW3000 and pFW4000, the R6K origin on pFW1000 was replaced by the *E. coli* origins p15A, pSC101ts or ColE1, respectively.

The three plasmids pFW1004, pFW2100 and pFW2500 were used for genetic manipulation in BU. The P_BT1311_-*lacI*-P_lacO23_ sequence amplified from pFW025 was cloned into pFW1000 and pFW2000 using the BamH I site, generating pFW1001 and pFW2001, respectively. Subsequently, the *Fncpf1* gene amplified from pT7FnCpf1 was cloned into pFW1001 and pFW2001 using the BamH I site, generating pFW1004 and pFW2100, respectively. Next, the IPTG inducible promoter P_lacO23_^36^ and *recT* cloned from *E. coli* DH5α were cloned into pFW2100 using the Sal I site, generating plasmid pFW2500.

To construct plasmids for deletion of genes or PULs, the promoters P_BT1311_ or P_BfP1E6_ controlling crRNA, *rrnB* T2 terminator as well as two homologous arms (∼500-bp for single gene deletion and ∼1000-bp for PUL deletion) were cloned into pFW1004, pFW2100 or pFW2500 using the BamH I site, yielding a set of plasmids for targeted gene or PUL deletions. For plasmids used for gene insertions, the promoter P_BT1311_ controlling the crRNA, *rrnB* T2 terminator as well as two ∼1000-bp homologous arms and gene fragments were cloned into pFW2100, yielding the final plasmids pFW2028 and pFW2059. The gene fragment replacement (for the generation of *rhaR’*) of inactive *rhaR* (BU) gene was designed based on the RhaR sequence from WP_005834782.1.

### Conjugation

All plasmids (pNBU2-ermGb derivatives and shuttle plasmids derivatives) were introduced into *Bacteroides* via conjugation with *E. coli* BW29427, which harbors the conjugative machinery integrated into the chromosome^75^. To perform the conjugation, *E. coli* BW29427 was grown until early stationary phase. Next, cell pellets were collected by centrifugation at 3,200 x g for 5 min and washed once with fresh LB media. The washed cell pellets were combined with *Bacteroides* cultures (OD_600_ approximately equal to 0.5-0.6) at a ratio of 1:10 (donor:recipient, v/v). The mixed culture was pelleted and resuspended in 0.2 mL BHI media and then spotted onto BHIAD (BHI + 10%ABB + DAP) plates prior to anaerobic incubation at 37 °C for 24 hr. Next, we collected the cells from the plate and resuspended into 1 mL BHI. The culture was plated on BHIAEG (BHI + 10%ABB + Em + Gm) plates using a range of dilutions and anaerobically incubated at 37 °C for 2 days. The pNBU2-ermGb plasmid derivatives harbor the IntN2 tyrosine integrase, which mediates the recombination between the *attN* site on the plasmid and one of the two *attBT* sites at the 3’ end of the tRNA^Ser^ gene (*BACUNI_RS18270* and *BACUNI_RS18350*) on the BU chromosome^73^. Thus, all pNBU2-ermGb plasmid derivatives were integrated onto the chromosome following conjugation^36^. As all the shuttle plasmids derivatives contain the *Bacteroides* replication origin pB8-51 and lack the *intN2* gene, these plasmids may continue to be replicated and potentially maintained over time.

### Characterization of shuttle plasmid stability

The BU strains harboring shuttle plasmids pFW1000, pFW2000, pFW3000 and pFW4000 were first grown at 37 °C in ABB media with erythromycin for 12-16 hr. Next, these strains were diluted 20-fold into fresh ABB media lacking antibiotics every 12 hr for 15 passages. The same volume of the diluted cell cultures was plated on BHIA (BHI + 10% ABB) and BHIAE (BHI + 10% ABB + Em) plates to determine the number of colonies containing the plasmid compared to the total number of colonies per unit volume. We evaluated plasmid stability based on the ratio of CFU on antibiotic selective plates to the total number of CFU on plates without antibiotic selection. Colony forming units (CFU) were determined at the following passages: 1, 3, 5, 7, 9, 11,13 and 15.

### Markerless gene deletion and insertion in Bacteroides

The plasmids pFW1004, pFW2100 and pFW2500 were used as the original vector for genome editing (**Table S2**). Plasmids for gene deletion and insertion were first transformed into *E. coli* BW29427 and then introduced into the *Bacteroides* strains using conjugation. The transconjugants were then grown anaerobically in ABB media supplemented with Em at 37 °C for 12-16 hr. The cell cultures were diluted 10-fold and plated on BHIAEI (BHI + 10% ABB + Em + IPTG) plates. The plates were anaerobically incubated at 37 °C for 2 days until the colonies were observed. We performed colony PCR using BioRed PCR mix (Bioline) to screen for genome modifications. Colonies with the correct genome modifications based on the colony PCR results were anaerobically cultured in ABB media and passaged three times every 12-16 h with a dilution factor of 20-fold. After this period, the cell cultures were diluted 10^−3^ to 10^−4^ and plated on BHIA agar plates and incubated at 37 °C for 1-2 days until the colonies were observed. Next, single colonies were picked and streaked on BHIA and BHIAE agar plates for replica plating. Colonies that could only grow on BHIA plates were selected as the final mutants that had lost the plasmids. We evaluated the efficiency of genome modification by computing the number of correct mutants divided by the total number of tested colonies.

### Construction of barcoded BU strains

To distinguish the PUL deletion mutants in the mutant pool, randomly generated DNA barcodes (4 bp) were introduced onto the chromosome of BU prior to gene deletion. We constructed Δ*PUL12*, Δ*PUL34* and Δ*PUL49* by first deleting the given PUL and then introducing the barcode into the PUL deletion background. In other cases, the barcoded strains were generated by introducing the 4 bp barcode onto the chromosome while simultaneously deleting the tryptophanase gene (*tyrP*) (**Fig. 2a**). To this end, a library of pFW2026 plasmids were constructed and introduced into the BU WT via conjugation. The barcoded plasmids were sequenced using Sanger Sequencing (Functional Biosciences) prior to gene deletion. The final set of *tyrP* deletion mutants were used as the barcode-tagged strains in this study.

### Luciferase assay to quantify gene expression in Bacteroides

All NanoLuc luciferase assays to quantify gene expression were performed with cell lysate according to the procedure described in the Nano-Glo Luciferase Assay System kit (Promega). For these experiments, *Bacteroides* strains were first anaerobically cultured at 37 °C in ABB media for 12-16 hr. Next, cultures were inoculated into BHI media and incubated at 37 °C anaerobically until the OD_600_ reached ∼0.6. To characterize the strengths of the RBS and promoter sequences, 4 mL of the cultures was centrifuged at 13,800 x g for 3 min. Cell pellets were lysed by resuspending into 400 µL of 1X BugBuster (Novagen) in 1X PBS (MP Biomedicals). To characterize the inducible promoters, 20 mL of cultures were harvested via centrifugation at 3,200 x g for 10 min. Cell pellets were resuspended into 400 µL of 1X BugBuster in 1X PBS containing 0.5 µL of rLysozyme™ Solution (EMD Millipore). Next, 10 µL of the cell lysate was mixed with an equal volume of NanoLuc Reaction buffer containing the substrate. The relative light units (RLU) were measured using a plate reader (Spark 10M, Tecan). The luminescence value was normalized to the OD_600_ value of the culture prior to lysis.

### Glycan utilization characterization of BU strains

The BU WT strain was incubated in ABB media at 37 °C anaerobically without shaking for 12-16 hr and then inoculated into ABB media again and incubated at 37 °C anaerobically until the culture reached exponential phase (OD_600_ ∼1.0). Next, the cell pellets were collected by centrifugation at 3,200 x g for 10 min, and then washed with BMM-C media (*Bacteroides* minimal media^42^ (BMM) without glucose, **Table S4**). The washed cell pellets were resuspended into BMM-C media to a final OD_600_ of approximately 1. These cultures were inoculated into a 96-well plate (Greiner Bio-One) containing BMM-glycan (BMM-C media supplemented with a given glycan, **Table S4**) to an initial OD_600_ of 0.05. These plates were incubated at 37 °C anaerobically. Cell growth determined by OD_600_ was monitored using a Tecan F200 plate reader every 12 hr or 30 min depending on the experimental design. All the glycans used in this study are listed in **Table S5**.

### Growth characterization of BU WT and Δtdk with 5-fluorodeoxyuridine

The BU WT and Δ*tdk* strains were grown in ABB media at 37°C anaerobically for 12-16 hours. Next, cultures were diluted by 20-fold into ABB media and then incubated at 37 °C anaerobically until the culture reached exponential phase (OD_600_ ∼1.0). Next, the cultures were diluted into ABB and ABB containing 200 µg/mL 5-fluorodeoxyuridine (FudR, Sigma) to an initial OD_600_ of 0.025. The growth of the strains was monitored based on OD_600_ on a Tecan F200 plate reader every 30 min for 24 hr.

### B. fragilis Δxyl xylose utilization assay

The *B. fragilis* DSM 2151 wild-type and *B. fragilis* Δ*xyl* strains were grown at 37°C anaerobically in ABB media for 12-16 hr. The cultures were then diluted by 20-fold into ABB media and then incubated at 37 °C anaerobically until the culture reached exponential phase (OD_600_ ∼1.0). The cell pellets were collected by centrifugation at 3,200 x g for 10 min and washed with BMM-C media. The cell pellets were resuspended into BMM-C at an OD_600_ of approximately 1 and then inoculated into BMM-xylose (5 g L^-1^ xylose, MP Biomedicals) to an OD_600_ of 0.025 in a 96-well plate (Greiner Bio-One). The growth of the strains was measured based on OD_600_ using a Tecan F200 plate reader and measurements were taken every 30 min for 54 hr.

### Characterization of B. thetaiotamicron Δlevan strain

The *B. thetaiotaomicron* ATCC 29148 (VPI-5482) wild-type and *B. thetaiotamicron* Δ*levan* (deletion of *BT1754*-*BT1765*) strains were grown at 37 °C anaerobically in ABB media for 12-16 hr. Cultures were diluted by 20-fold into ABB media and then incubated at 37 °C anaerobically until the culture reached exponential phase (OD_600_ ∼1.0). Cell pellets were collected by centrifugation at 3,200 x g for 10 min, and then washed with BMM-C media. The cell pellets were resuspended into BMM-C to an OD_600_ of approximately 1 and then inoculated into BMM-levan (5 g L^-1^ levan, Sigma) using an initial OD_600_ of 0.05 in a 96-well plate (Greiner Bio-One). Cell growth was determined based on OD_600_ using a Tecan F200 plate reader every 30 min for 60 hr.

### Whole-genome transcriptional profiling of BU

The BU WT and genome modified mutants were grown at 37 °C anaerobically in ABB media for 12-16 hr. Cultures were then diluted by 20-fold into ABB media and then grown at 37 °C anaerobically until the culture reached exponential phase (OD_600_ ∼1.0). The cell pellets were collected by centrifugation at 3,200 x g for 10 min and washed with BMM-C media. Next, the cell pellets were resuspended into BMM-C and then inoculated into 5 mL BMM or BMM-xyloglucan media (5 g L^-1^ xyloglucan, Megazyme) to an initial OD_600_ of 0.05. Cells were harvested for total RNA extraction when the OD_600_ reached 0.6-0.8. RNA was extracted using the RNeasy Mini Kit (Qiagen) and genomic DNA was digested using the RNase-Free DNase Set (Qiagen). The RNA integrity number (RIN, an algorithm for assigning integrity values to RNA measurements) was measured by Agilent TapeStation 4150 with the Agilent High Sensitivity RNA ScreenTape. The samples were then processed by GENEWIZ (NJ, USA) by performing rRNA depletion, cDNA library preparation and sequencing. rRNA depletion was performed by using Ribozero rRNA Removal Kit (Illumina). The NEBNext Ultra RNA Library Prep Kit (NEB) was used for RNA sequencing library preparation. The sequencing libraries were sequenced with an Illumina HiSeq instrument. Image analysis and base calling were conducted by the HiSeq Control Software (HCS). Raw sequence data generated from Illumina HiSeq was converted into FASTQ files and de-multiplexed using Illumina’s bcl2fastq 2.17 software. One mismatch was allowed for index sequence identification.

The compressed FASTQ files were quality checked using the FastQC tool v0.11.8^76^. Packages from the BBTools suite v38.42^77^ including BBDuk, BBSplit, and BBMap were used to filter high quality reads, trim adapters using built-in adapter reference file, remove rRNA reads, and map sequences to the reference genome (*B. uniformis* DSM 6597). The featureCounts package v1.6.4^78^ from the SubRead suite was used for read mapping to gene features and quantifying raw counts for each transcript. The DESeq2 Bioconductor library v4.0.3^79^ was used in R v4.0.4 to normalize read counts across samples and quantify differential gene expression using a negative binomial generalized linear models with apeglm shrinkage estimator^80^.

### Quantitative reverse transcription PCR (qRT-PCR)

The BU WT and genome modified mutants were grown at 37°C anaerobically in ABB media for 12-16 hr. Cultures were diluted by 20-fold into ABB media and then grown at 37°C anaerobically until the culture reached exponential phase (OD_600_ ∼1.0). Cell pellets were collected by centrifugation at 3,200 x g for 10 min and then washed with BMM-C media. The washed cell pellets were then resuspended into BMM-C and then inoculated into 5 mL BMM or BMM-xyloglucan (5 g L^-1^) media to an initial OD_600_ of 0.05. Cells were harvested for total RNA extraction when the OD_600_ reached 0.6-0.8. Total RNA was extracted with RNeasy Mini Kit (Qiagen) and treated with DNase I (Invitrogen) to remove the genomic DNA. We performed cDNA synthesis with 0.5-1 μg of total RNA using the iScript Select cDNA Synthesis Kit (Bio-Rad Laboratories). We performed Real Time quantitative PCR (RT-qPCR) on the Bio-Rad CFX connect Real-Time PCR instrument with SYBR™ Green PCR Master Mix (Thermo Fisher Scientific). We computed the fold changes of the target genes by normalizing to the two reference genes 16S rRNA gene and *gyrA* with against of their geometric mean^81^. We computed the fold change using the equation 2^*x*^ where 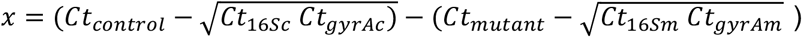 where *Ct*_*control*_, *Ct*_*mutant*_, *Ct*_16*Sc*_, *Ct*_*gyrAc*_, *Ct*_16*Sm*_, *Ct*_*gyrAm*_ denote the Ct value of the gene in Δ*tyrP-24*, 16S rRNA gene in Δ*tyrP-24, gyrA* in Δ*tyrP-24*, target gene of the PUL mutant strain, 16S rRNA gene of the PUL mutant strain and *gyrA* in the PUL mutant strain, respectively.

### Pooled barcoded BU mutant experiments

The barcoded BU control strain Δ*tyrP-24* and PUL deletion mutants were grown at 37 °C anaerobically in ABB media for 12-16 hr. Cultures were then diluted by 20-fold into ABB media and grown at 37 °C anaerobically until the culture reached exponential phase (OD_600_ ∼1.0). Next, all the strains were mixed in equal proportions based on OD_600_ and then washed with DM29 media (lacking a carbon source, **Table S6**). Next, the cells were resuspended in DM29 and diluted to a final OD_600_ of approximately 1. The mixture of strains was inoculated into 2 mL 96-deep-well plates (Nest Scientific) containing 1 mL DM29-glucose or DM29 supplemented with different glycans (**Table S6**) to an initial OD_600_ of 0.05 and incubated at 37 °C anaerobically for varying lengths of time and passaging was performed by diluting the cultures by 20-fold up to two times. After 48 hr of cultivation, OD_600_ was measured by Tecan F200 plate reader and cell pellets were collected for NGS sequencing (**Fig. 4a**).

### Gnotobiotic mouse experiments

All germ-free mouse experiments were performed following protocols approved by the University of Wisconsin-Madison Animal Care and Use Committee. Three diets were used in this experiment: high fiber diet (Chow diet, Purina, LabDiet 5021), high fat diet (Envigo, TD.88137) and fiber free diet (Envigo, TD.190849) (**Supplementary Data 3**). Note that the high fiber diet contains higher fiber compared to the High fat diet or Fiber free diet and was thus referred to as high fiber diet. The barcoded BU control strain Δ*tyrP-24* and PUL deletion mutants were grown at 37 °C anaerobically in ABB media for 12-16 hr. Cultures were diluted by 20-fold into ABB media and then grown at 37 °C anaerobically until the culture reached exponential phase (OD_600_ ∼1.0). All strains were mixed in equal proportions based on OD_600_ and transferred to Hungate tubes (Chemglass) on ice prior to oral gavage. We used 8-week old C57BL/6 gnotobiotic mice (wild-type) fed the specific diets a week prior to oral gavage (**Fig. 5a**). At this time, 0.2 mL of mutant pool was introduced into the mice by oral gavage inside a Biological Safety Cabinet (BSC) and the mice were housed in biocontainment cages (Allentown Inc.) for the duration of the experiment. Mice were maintained on the same experimental diets with autoclaved water or water supplemented with glycans for two weeks after colonization. The concentration of inulin and pectic galactan in the drinking water was 5 g L^-1^. 2 g L^-1^ of glucomannan solution were prepared and autoclaved. The autoclaved solution was centrifuged at 3,200 x g for 5 min, and the supernatant of the glucomannan solution was used for the mouse experiments. Groups of mice (4-5 mice) fed a given diet were co-housed in a single cage. Fecal samples were collected every 2-3 days after oral gavage. At the end of the experiment, mice were euthanized, and the cecum samples were collected for NGS sequencing and CFU plating.

### CFU counting for gnotobiotic mouse experiments

The cecum contents weighing 0.2-0.3 mg were collected into sterilized Eppendorf tubes and then resuspended into 1 mL anaerobic ABB media. The suspended contents were homogenized by two 3.2 mm stainless steel beads (BioSpec Products) with intermittent vortex (BR-2000, Bio-Rad) for 5 min. The homogenized contents were then transferred into 9 mL of ABB media. This mixture was diluted 10^3^ to 10^5^ times for CFU plating. We plated 100 µL of the diluted solutions on BHIA plates and incubated in an anaerobic chamber at 37 °C for 36-48 hr until colonies were visible on the plates. We computed the CFU and divided this number by the measured weight of each cecum content.

### DNA extraction from fecal and cecum samples

The DNA extraction for fecal and cecum samples was performed as described previously with some modifications^82^. Fecal samples (∼50 mg) were transferred into solvent-resistant screw-cap tubes (Sarstedt Inc) with 500 μL of 0.1 mm zirconia/silica beads (BioSpec Products) and one 3.2 mm stainless steel bead (BioSpec Products). The samples were resuspended in 500 μL of Buffer A (200 mM NaCl (DOT Scientific), 20 mM EDTA (Sigma) and 200 mM Tris·HCl pH 8.0 (Research Products International)), 210 μL 20% SDS (Alfa Aesar) and 500 μL phenol/chloroform/isoamyl alcohol (Invitrogen). Cells were lysed by mechanical disruption with a bead-beater (BioSpec Products) for 3 min twice to prevent overheating. Next, cells were centrifuged for 5 min at 8,000 x g at 4°C, and the supernatant was transferred to a Eppendof tube. We added 60 μL 3M sodium (Sigma) and 600 μL isopropanol (Koptec) to the supernatant and incubated on ice for 1 hr. Next, samples were centrifuged for 20 min at 18,000 x g at 4°C. The harvested DNA pellets were washed once with 500 μL of 100% ethanol (Koptec). The remaining trace ethanol was removed by air drying the samples. Finally, the DNA pellets were then resuspended into 200 μL of AE buffer (Qiagen). The crude DNA extracts were purified by Zymo DNA Clean & Concentrator™-5 kit (Zymo Research) for NGS sequencing.

### Butyrate producer community experiments and sample collection

All BU strains and butyrate producers (*A. caccae* DSM 14662, *C. comes* ATCC 27758, *E. rectale* ATCC 33656 and *R. intestinalis* DSMZ 14610) were inoculated into ABB media with the exception of *R. intestinalis* which was inoculated into BHI media and grown at 37 °C anaerobically. After 16-24 hr, the cultures were diluted by 20-fold into the same media until the culture reached exponential phase (OD_600_∼1.0). The cultures were centrifuged at 3,200 x g for 10 min, and then washed with DM29 media. The washed cells were then resuspended into the same DM29 media and the OD_600_ was diluted to 1. The BU and each butyrate producer cultures were mixed in equal proportions based on OD_600_ and inoculated into DM29-glucose (5 g L^-1^) or DM29 media supplemented with glycogen, glucomannan, inulin, laminarin, pectic galactan, pectin, pullulan or xyloglucan (**Table S6**). The pairwise communities and monocultures were introduced into 2 mL 96-deep-well plates (Nest Scientific) to an initial OD_600_ of 0.05. We have 4 plates for each experiment and each plate was taken out for sampling every 12 hr for a total of 48 hr. At each time point, OD_600_ was measured with Tecan F200 (with 5-10 dilution based on the density of the cell culture) for the monitor of cell growth and cell pellets were collected for NGS sequencing.

We performed butyrate measurements at a single time point. Specifically, 2 µL of H_2_SO_4_ (Sigma) was added to the supernatant of each sample to precipitate any components that was incompatible with the mobile phase. The samples were then centrifuged at 3,200 x g for 10 min and then filtered through a 0.2 µm filter (Pall Corporation) using a vacuum manifold (KNF Neuberger) before transferring to HPLC vials (Thermo Scientific). Butyrate concentrations were measured with an Agilent 1260 infinity HPLC system equipped with a quaternary pump, chilled (4°C) autosampler, vacuum degasser, refractive index detector, Aminex HPX-87H column and Cation-H guard column (300×7.8mm, BioRad). We used 0.02 N H_2_SO_4_ as the mobile phase with a flow rate of 0.5 mL min^-1^ at a column temperature of 50°C. The injection volume of all samples was 50 µL and the run time was 30 min. Data analysis was performed using the Chem Station Rev.C01.08 software (Agilent Technologies).

### Bacterial genome DNA extraction and next-generation sequencing

All the genomic DNA extraction and next-generation sequencing sample preparation was performed as described previously^62^. Briefly, bacterial genome DNA extraction was carried out using a modified version of the Qiagen DNeasy Blood and Tissue Kit protocol in 96-well plates^62^. Genomic DNA concentrations were measured using the SYBR Green fluorescence assay (Bio-Rad) and then normalized to 1 ng μL^-1^ or 2 ng μL^-1^ for genomic DNA extracted from fecal and cecal samples by diluting in molecular grade water (VWR International) using a Tecan Evo Liquid Handling Robot. We performed PCR using custom dual-indexed primers^43,62^ targeting the V3-V4 region of the 16S rRNA gene using the diluted genomic DNA samples as template. These libraries were purified using the DNA Clean & Concentrator™-5 kit (Zymo) and eluted in water. Sequencing was performed on an Illumina MiSeq using MiSeq Reagent Kit v3 (600-cycle) to generate 2×300 paired end reads. For the sequencing of barcoded BU strains, the 200 bp amplicon libraries containing 4-bp barcodes were generated using the procedure described above and PCR amplified with custom dual-indexed primers listed in **Table S7**. The obtained libraries were sequenced on an Illumina MiSeq using MiSeq Reagent Nano Kit v2 (500-cycles) to generate 2×250 paired end reads or 2×300 paired end reads.

### Bioinformatic analysis of species and barcoded strain abundances

For the 16S rDNA gene sequencing data analysis, we used previously described custom scripts in Python 3.7 and aligned to a reference database of V3-V4 16S rRNA gene sequences as previously described^43,83^. Relative abundance was calculated as the read count mapped to each species divided by the total number of reads for each condition. The absolute abundance of each species was calculated by multiplying the relative abundance determined by NGS sequencing by the OD_600_ measurement for each sample. For the sequencing analysis of barcode-tagged strains, paired end reads were first merged using PEAR (Paired-End reAd mergeR) v0.9.10^83^ after which barcodes were extracted by searching for exact matches of the immediate upstream and downstream sequences within the reads. Barcodes with less than 100% match were discarded. The relative abundance of each barcoded strain was calculated as the number of reads mapped to each barcode divided by the total reads that mapped to each condition. Absolute abundance of each mutant was calculated by multiplying the relative abundance by the OD_600_ measurement of each condition.

### Characterization of butyrate producer growth in BU conditioned media

BU was grown anaerobically at 37 °C in ABB media for 12-16 hr. Cells were harvest by centrifugation at 3,200 x g for 10 min, washed with DM29 and then inoculated into 12 mL DM29-glucose (5 g L^-1^) and DM29 media supplemented with glucose or different glycans and incubated at 37°C anaerobically for 16 hr. Next, cell pellets were collected with centrifugation at 3,200 x g for 10 min and washed twice with DM29. The washed cell pellets were resuspended into same volume of the same media (glucose or glycan media) and incubated at 37°C for 3 hr to allow cell growth and glycan utilization. The supernatants were collected by centrifugation at 3,200 x g at 4°C for 30 min and the pH was adjusted to the same value as fresh media 5N KOH (Alfa Aesar) with the Mettler Toledo InLab Micro pH electrode. For DM29-glucose conditioned media, the residual glucose was measured using the Amplex™ Red Glucose/Glucose Oxidase Assay Kit (Sigma). Based on the measurement, glucose was restored to the initial concentration of 5 g L^-1^ such that the fresh media and BU conditioned glucose media had the same glucose concentration. The pH adjusted conditioned media was filtered twice using a 0.2 μm filter (Whatman) to remove BU cells. Butyrate producers were grown in ABB (AC, CC and ER) or BHI (RI) at 37 °C anaerobically for 16-24 hr, and passaged in the same media with a dilution of 20-fold until they reached exponential phase (OD_600_∼1.0). Cells were harvested by centrifugation at 3,200 x g for 10 min and washed with DM29 and resuspended in DM29 to an OD_600_ of approximately 1. The cultures were inoculated into both fresh media and conditioned media with an initial OD_600_ of 0.05 and anaerobically grown at 37 °C in a 96-well plate (Greiner Bio-One) without shaking. Cell growth was monitored by plate reader (Tecan F200).

### Characterization of butyrate producer growth with fermentation end products as primary carbon source

Butyrate producers were grown in ABB (AC, CC and ER) or BHI (RI) at 37 °C anaerobically for 16-24 hr, and passaged in the same media with a dilution of 20-fold until they reached exponential phase (OD_600_∼1.0). Cells were harvested by centrifugation at 3,200 x g for 10 min and washed with DM29 and resuspended in DM29 to an OD_600_ of approximately 1. The cell cultures were then inoculated into DM29 with addition of 0, 10, 25 or 50 mM of acetate, propionate or succinate or the mixture of fermentation end products (25 mM of acetate, 25 mM of propionate and 50 mM of succinate). Cells were cultured in a 96-well plate (Greiner Bio-One) with an initial OD_600_ of 0.05. Cell growth was monitored using a Tecan F200 plate reader.

### Characterization of growth of butyrate producers in DM29-glucose or DM29-glycan media supplemented with fermentation end products

Butyrate producers were grown in ABB (AC, CC and ER) or BHI (RI) at 37 °C anaerobically for 16-24 hr and passaged in the same media with a dilution of 20-fold until they reached exponential phase (OD_600_∼1.0). Cells were harvested by centrifugation at 3,200 x g for 10 mins and washed with DM29 and resuspended in DM29 to an OD_600_ of approximately 1. The cultures were then inoculated into DM29-glucose (5 g L^-1^) and DM29 media supplemented with different glycans (**Table S6**) with or without the addition of 25 mM of acetate, 25 mM of propionate and 50 mM of succinate. Cells were cultured in a 96-well plate (Greiner Bio-One) using an initial OD_600_ of 0.05. Cell growth was monitored using a Tecan F200 plate reader.

### Characterization of butyrate producer growth in cell membrane treated glycan media

The preparation of the cell membrane fraction including inner and outer membrane anchoring enzymes was modified based on the procedures described from Millipore Sigma (https://www.sigmaaldrich.com/technical-documents/protocols/biology/purifying-challenging-proteins/cell-disruption-and-membrane-preparation.html). BU was grown anaerobically at 37 °C in ABB media for 12-16 hr. Cells were harvest by centrifuge at 3,200 x g for 10 min and washed with DM29 and inoculated into 20 mL of DM29-glycan media (**Table S6**) and grown anaerobically at 37°C for 16 hr. Cell pellets were collected and washed with DM29 at 3,200 x g, 4°C for 1 hr. The washed cell pellets were resuspended into 6 mL of DM29 with the addition of 60 μL of pefabloc (100 mM, Sigma) and 6 μL of DNase I (20 mg mL^-1^, Sigma). Cells were lysed via sonication (5 s on and 5 s off for 2.5 min, performed twice, Sonicator 3000, Misonix). The cell lysis was centrifuged at 3,200 x g at 4 °C for 10 min. The supernatants were collected and filtered twice with 0.45 μm filter (Whatman) to remove the remaining intact cells. The collected supernatants were centrifuged at 300,000 x g for 2 hr with Optima MAX-XP ultracentrifuge with SW 55 Ti Swinging-Bucket Rotor (Beckman Coulter). The pellets consisting of the cell membrane fractions were resuspended into 10 mL of the same glycan containing media and incubated in the anaerobic chamber at 37°C for 16 hr. The cell membrane treated glycan media as well as the respective fresh glycan containing media were filtered with 0.2 μm filter (Whatman). Butyrate producers were grown in ABB (AC, CC and ER) or BHI (RI) at 37°C anaerobically for 16-24 hr, and passaged in the same media with a dilution of 20-fold until they reached exponential phase (OD_600_∼1.0). Cells were harvested by centrifugation at 3,200 x g for 10 min and washed with DM29 and resuspended in DM29 to an OD_600_ of approximately 1. Cultures were then inoculated into fresh media supplemented with different glycans and cell membrane treated glycan media to an initial OD_600_ of 0.05 and incubated at 37 °C without shaking in a 96-well plate (Greiner Bio-One). Cell growth was monitored by plate reader (Tecan F200).

### Measurements of fructose

The fructose in BU conditioned inulin media was measured using a Fructose assay kit (Sigma). BU was grown anaerobically at 37°C in ABB media for 12-16 hr. Cells were harvest by centrifuge at 3,200 x g for 10 min and washed with DM29 and inoculated into DM29 media supplemented with 5 g L^-1^ inulin for 16 hr. Next, cell pellets were collected with centrifugation at 3,200 x g for 10 mins and washed twice with DM29. The washed cell pellets were resuspended into same volume of DM29-inulin and incubated at 37°C for 3 hr. The supernatants were collected with centrifugation at 3,200 x g for 10 min and filtered using a 0.2 μm filter (Whatman) prior to fructose measurement. The fructose concentrations in control solutions were also measured (5 g L^-1^ inulin solution and DM29-inulin media).

### Growth characterization of AC with fructose as the primary carbon source

AC was grown anaerobically in ABB media at 37 °C for 16-24 hr, and passaged in the same media with a dilution of 20-fold until they reached exponential phase (OD_600_∼1.0). Cells were harvested with centrifugation at 3,200 x g for 10 min and washed with DM29 and resuspended into DM29 again to OD_600_ of approximately 1. The cultures were then inoculated into DM29-fructose (5 g L^- 1^) and DM29-glucose (5 g L^-1^) media in a 96-well plate (Greiner Bio-One) to an initial OD_600_ of 0.05. Cell growth was monitored using a Tecan F200 plate reader.

### Bioinformatic analysis of PULs in human gut microbiome metagenome datasets

Two human gut microbiome datasets, which contained 154,723 metagenome-assembled genomes (MAGs) from 9,428 human gut microbiomes^51^ and 92,143 MAGs from 11,850 human gut microbiomes^52^, were used to find PULs (including *PUL11, PUL12, PUL17, PUL18, PUL2, PUL37* and *PUL43*) of BU. All MAGs were annotated by Prodigal v2.6.3^84^ and DIAMOND BLASTP v0.9.28.129^85^ was used to find hits to reference proteins within each PUL with settings of “-k 1 -e 1e-5 --query-cover 25 --id 50”. Annotation by dbCAN2^86^ was used to identify glycoside hydrolases (GHs) from the DIAMOND BLASTP hits. We then used three criteria to assign PUL positive hits: 1) satisfies the requirement of essential genes for the function of each PUL; 2) GH annotation of the essential gene was consistent with the PUL reference; 3) essential genes of each PUL were within a gene array of size less than 30 genes (**Fig. S14**)

For all MAGs, we used CheckM v1.0.11 to evaluate genome completeness and to assign the total MAG dataset into subsets with > 50%, > 60%, > 70%, > 80%, and > 90% genome completeness. The finer taxonomic information of MAGs within Bacteroidetes was parsed by GTDB-Tk v0.1.3 with default settings. The abundance ratios of PULs in all MAGs, only BU MAGs, and other non-BU MAGs were calculated from the resulted PUL positive hit table. The cooccurrence ratios of PULs in all MAGs was also parsed out accordingly. Results of abundance ratios and cooccurrence ratios that were calculated from a series of genome completeness subsets were combined and visualized together.

### gLV modeling and parameter inference

We use the generalized Lotka-Volterra (gLV) model to describe growth dynamics and inter-species interactions. Specifically, the gLV model can be written as the following ordinary differential equation:

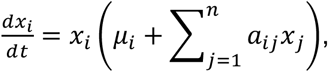

where *x*_*i*_ and *x*_*j*_ are the absolute abundance of species *i* and *j*, respectively, the non-negative parameter *μ*_*i*_ describes the basal growth rate of species *i*, integer *n* is the total number of species in an experiment, and *a*_*ij*_ is a parameter that quantifies how the abundance of species *j* modifies the growth rate of species *i*. When *j*≠*i*, the parameter *a*_ij_ is called the inter-species interaction coefficient, and it is called the intra-species interaction coefficient when *j*=*i*. For a monoculture experiment (i.e., *n*=1), the gLV model simplifies to the logistic growth equation. The gLV model has been used before to describe inter-species interactions in complex microbial communities and to predict their emerging community dynamics^43^.

To determine the gLV parameters θ = (*μ*_1_, …, *μ*_*n*_, *a*_11_, …, *a*_*nn*_) in each carbon source, we performed a set of *p* experiments that include monoculture of all species and some co-culture experiments (e.g., BU and butyrate producer pairs). For the *q*-th experiment, we took *m* time-series abundance measurements for three biological replicates with mean *x*_*q*_ = (*x*_*q*1_, … *x*_*qm*_) and standard deviation σ_*q*_ = (σ_*q*1_, …, σ_*qm*_). Given these observations in all *p* experiments: **x** = (*x*_1_, … *x*_*p*_) and **σ** = (σ_1_, …, σ_*p*_), the posterior distribution of θ, which we denote by *P*(θ |**x, σ**), is found using an adaptive Markov Chain Monte Carlo (MCMC) method. In particular, we assume that uncertainty for the *k-*th measurement in the *q*-th experiment is modeled by an additive and independent noise, which is distributed according to 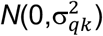. Given a fixed θ, we first simulate the gLV model for each experiment *q* to obtain the model predicted abundance 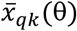 at every instant *k*. The likelihood to observe the sequence of abundance measurements x can then be computed as:

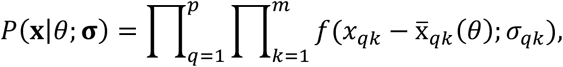

where *f*(∙ ; σ_*qk*_) is the probability density function for the normal distribution 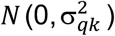. The posterior probability can then be described according to Bayes rule as *P*(θ|**x**) ∝ *P*(**x**|θ; **σ**)*P*(θ), where *P*(θ) is the prior parameter distribution. In this paper, for all cases except xyloglucan co-culture, we chose uniform priors for the parameters. Specifically, the priors for all growth rates *μ*_*i*_ are *U*(0,2), the priors for all inter-species interaction coefficients *a*_ij_ are *U*(−2.5,2.5), and the priors for all intra-species interaction coeffects *a*_ii_ are *U*(−2.5,0). The boundaries for these distributions are chosen to be sufficiently large to contain similar gLV parameters identified in the literature^43^ and to ensure positive, bounded growth when simulating monoculture experiments. A normal prior distribution was used to infer parameters in the xyloglucan co-culture experiments, and parameters of this distribution are listed in **Supplementary Data 4**. Since gLV models cannot capture cell growth in death phase, if the OD_600_ of the *k*-th measurement in a monoculture experiment *q* drops more than 20% from that of the (*k*-1)-th measurement, then *x*_qk_ is not used for parameter inference. Data points excluded for parameter inference were indicated with an empty circle in **Fig. S18**. In monoculture, many Δ*PUL* strains did not grow (OD_600_<0.08 for all time) in media supplemented with glycans. The growth rates (*μ*_*i*_) for these Δ*PUL* strains in the respective media are set to 0. Similarly, for Δ*PUL*/butyrate producer coculture experiments, no inference was made on the interaction coefficients between the two species in conditions that did not display growth, and thus the coefficients were set to 0.

An adaptive, symmetric, random-walk Metropolis MCMC algorithm^87^ is then used to draw samples from the posterior distribution *P*(θ|**x**). Specifically, given the current sample θ^(*l*)^ at step *l* of the Markov chain, the proposed sample for step (*l*+1) is θ^(*l*+1)^= θ^(*l*)^+δ^(*l*)^, where δ^(*l*)^ is drawn randomly from a normal distribution. The algorithm is adaptive in the sense that the covariance of this normal distribution is given by 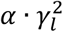, where 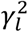 is the covariance of θ^(1)^,…, θ^(*l*)^ and *α* is a positive parameter. In this paper, depending on the carbon source, parameter *α* is either chosen to be 0.1or 0.5. The posterior probability *P*(θ^(*l*+1)^|**x**) of the proposed sample θ^(*l*+1)^ is then computed, and the proposed sample is accepted with probably 1 if 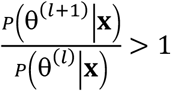, and it is accepted with probability 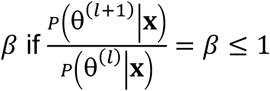.

The algorithm described above was implemented using custom code in MATLAB R2016a (The MathWorks, Inc., Natick, MA, USA), where the gLV models are solved using variable step solver ode23s. For each carbon source, we collected at least 300,000 MCMC samples after a burn-in period of at least 100,000 samples. The Gelman-Rubin potential scale reduction factor (PSRF) was used to evaluate convergence of the posterior distributions, where a PSRF closer to 1 indicates better convergence. We found that out of 186 parameters, 82% of them have PSRFs less than 1.2, and the median of PSRF is 1.03, indicating that the parameters drawn from MCMC had converged to the posterior parameter distribution. The marginal posterior distributions of the identified parameters are shown in Supplementary **Figs. S19 & S20**. The medians and the coefficient of variations (CVs) of these marginal distributions are summarized in **Supplementary Data 4**. The parameter medians were used to simulate the temporal abundance trajectory in **Fig. 6** and **Fig. S18**.

### Regression model for butyrate concentration

A previous study^62^ has shown that end point butyrate concentration in a microbial community in batch culture can be predicted by the absolute abundance of butyrate producers. Inspired by this, we propose the following linear regression model for end point butyrate concentration:

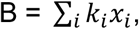

where B is the predicted butyrate concentration, *x*_i_ represents end point butyrate producer abundances, where *i*=AC, CC, ER, or RI, and *k*_*i*_ is a constant parameter. A major assumption in this model is that the parameter *k*_*i*_ is independent of carbon source in growth media, the absence/presence of BU, and the absence/presence of ΔPUL strains. The inferred parameters *k*_*i*_ for AC, CC, ER and RI were 19.2, 3.99, 8.31 and 19.35 mM OD_600_^-1^ respectively. The parameters *k*_*i*_ were obtained by performing least square regression on all monoculture and coculture experiments with a butyrate producer. Linear regression is performed using MATLAB function fitlm with intercept constrained to be 0.

## ACKNOWLEDGEMENTS

We would like to thank Yu-Yu Cheng, Bryce M. Connors, Ryan L. Clark, Erin Ostrem Loss, Susan E. Hromada for helpful advice about the design of experiments and data analysis. We would like to thank Michael Fischbach and Anthony Rush for providing plasmid pNBU2-bla-ermGb and Justin Sonnenburg and Michelle St Onge for providing the pWW3452 plasmid. We would like to thank Mcsean Mcgee, David Stevenson and Joshua Coon’s lab for helping with SCFA measurements. We would like to thank Aaron Hoskins and Xingyang Fu for providing access to key equipment used in study. Research was sponsored by the National Institutes of Health and was accomplished under Grant Number R35GM124774, National Institute of Biomedical Imaging and Bioengineering under grant number R01EB030340, Multi University Research Initiative (MURI) from under grant number W911NF-19-1-0269 and Department of Energy under grant number DE-FC02-07ER64494.

## AUTHOR CONTRIBUTIONS

O.S.V. and J.F. designed the research. J.F. carried out the experiments. N.V. and J.F. performed the mice experiments and F.E.R. provided guidance on the design of the mouse experiment. Y.Q. performed computational modeling. C.Z. performed bioinformatic analyses and K.A. helped with the design of bioinformatic analyses. S.E. and J.F. performed RNA-seq data analysis. J.F., O.S.V. and Y.Q. performed data analyses. F.L. wrote code for data analysis. J.F., O.S.V. and Y.Q. wrote the manuscript. All authors provided feedback on the manuscript. O.S.V. secured funding.

## CODE AVAILABILITY

The code used for computational modeling and data analysis is available upon request.

## REFERENCES

1 Koh, A., De Vadder, F., Kovatcheva-Datchary, P. & Backhed, F. From dietary fiber to host physiology: short-chain fatty acids as key bacterial metabolites. Cell 165, 1332–1345 (2016).

2 Fan, Y. & Pedersen, O. Gut microbiota in human metabolic health and disease. Nature Reviews Microbiology 19, 55–71 (2021).

3 Lloyd-Price, J., Abu-Ali, G. & Huttenhower, C. The healthy human microbiome. Genome Medicine 8, 51 (2016).

4 Desai, M. S. et al. A dietary fiber-deprived gut microbiota degrades the colonic mucus barrier and enhances pathogen susceptibility. Cell 167, 1339–1353 (2016).

5 Porter, N. T. & Martens, E. C. The critical roles of polysaccharides in gut microbial ecology and physiology. Annual Review of Microbiology 71, 349–369 (2017).

6 El Kaoutari, A., Armougom, F., Gordon, J. I., Raoult, D. & Henrissat, B. The abundance and variety of carbohydrate-active enzymes in the human gut microbiota. Nature Reviews Microbiology 11, 497–504 (2013).

7 Cantarel, B. L., Lombard, V. & Henrissat, B. Complex carbohydrate utilization by the healthy human microbiome. PloS one 7, e28742 (2012).

8 Wexler, A. G. & Goodman, A. L. An insider’s perspective: Bacteroides as a window into the microbiome. Nature Microbiology 2, 17026 (2017).

9 Sonnenburg, E. D. et al. Specificity of polysaccharide use in intestinal Bacteroides species determines diet-induced microbiota alterations. Cell 141, 1241–1252 (2010).

10 Larsbrink, J. et al. A discrete genetic locus confers xyloglucan metabolism in select human gut Bacteroidetes. Nature 506, 498–502 (2014).

11 Ndeh, D. et al. Metabolism of multiple glycosaminoglycans by Bacteroides thetaiotaomicron is orchestrated by a versatile core genetic locus. Nature Communications 11, 1–12 (2020).

12 Déjean, G. et al. Synergy between cell surface glycosidases and glycan-binding proteins dictates the utilization of specific Beta(1,3)-glucans by human gut Bacteroides. mBio 11, e00095 (2020).

13 Rogowski, A. et al. Glycan complexity dictates microbial resource allocation in the large intestine. Nature Communications 6, 7481 (2015).

14 Cartmell, A. et al. A surface endogalactanase in Bacteroides thetaiotaomicron confers keystone status for arabinogalactan degradation. Nature Microbiology 3, 1314–1326 (2018).

15 Ndeh, D. et al. Complex pectin metabolism by gut bacteria reveals novel catalytic functions. Nature 544, 65–70 (2017).

16 Briliute, J. et al. Complex N-glycan breakdown by gut Bacteroides involves an extensive enzymatic apparatus encoded by multiple co-regulated genetic loci. Nature Microbiology 4, 1571–1581 (2019).

17 Luis, A. S. et al. Dietary pectic glycans are degraded by coordinated enzyme pathways in human colonic Bacteroides. Nature Microbiology 3, 210–219 (2018).

18 Martens, E. C., Chiang, H. C. & Gordon, J. I. Mucosal glycan foraging enhances fitness and transmission of a saccharolytic human gut bacterial symbiont. Cell Host Microbe 4, 447–457 (2008).

19 Martens, E. C. et al. Recognition and degradation of plant cell wall polysaccharides by two human gut symbionts. PLoS Biology 9, e1001221 (2011).

20 Lapébie, P., Lombard, V., Drula, E., Terrapon, N. & Henrissat, B. Bacteroidetes use thousands of enzyme combinations to break down glycans. Nature Communications 10, 2043 (2019).

21 Tuncil, Y. E. et al. Reciprocal prioritization to dietary glycans by gut bacteria in a competitive environment promotes stable coexistence. mBio 8, e01068 (2017).

22 Rakoff-Nahoum, S., Foster, K. R. & Comstock, L. E. The evolution of cooperation within the gut microbiota. Nature 533, 255 (2016).

23 Rakoff-Nahoum, S., Coyne, M. J. & Comstock, L. E. An ecological network of polysaccharide utilization among human intestinal symbionts. Current Biology 24, 40–49 (2014).

24 Cuskin, F. et al. Human gut Bacteroidetes can utilize yeast mannan through a selfish mechanism. Nature 517, 165–169 (2015).

25 Sheridan, P. O. et al. Polysaccharide utilization loci and nutritional specialization in a dominant group of butyrate-producing human colonic Firmicutes. Microbial Genomics 2, e000043 (2016).

26 Qin, J. et al. A human gut microbial gene catalogue established by metagenomic sequencing. Nature 464, 59 (2010).

27 Almeida, A. et al. A unified catalog of 204,938 reference genomes from the human gut microbiome. Nature Biotechnology 39, 105–114 (2021).

28 Terrapon, N. et al. PULDB: the expanded database of polysaccharide utilization Loci. Nucleic Acids Research 46, D677–D683 (2017).

29 Baughn, A. D. & Malamy, M. H. A mitochondrial-like aconitase in the bacterium Bacteroides fragilis: implications for the evolution of the mitochondrial Krebs cycle. Proceedings of the National Academy of Sciences 99, 4662–4667 (2002).

30 Koropatkin, N. M., Martens, E. C., Gordon, J. I. & Smith, T. J. Starch catabolism by a prominent human gut symbiont is directed by the recognition of amylose helices. Structure 16, 1105–1115 (2008).

31 Garcia-Bayona, L. & Comstock, L. E. Streamlined genetic manipulation of diverse Bacteroides and Parabacteroides isolates from the human gut microbiota. mBio 10, e01762 (2019).

32 Hsu, P. D., Lander, E. S. & Zhang, F. Development and applications of CRISPR-Cas9 for genome engineering. Cell 157, 1262–1278 (2014).

33 Mali, P. et al. RNA-guided human genome engineering via Cas9. Science 339, 823–826 (2013).

34 Bortesi, L. & Fischer, R. The CRISPR/Cas9 system for plant genome editing and beyond. Biotechnology Advances 33, 41–52 (2015).

35 Zetsche, B. et al. Cpf1 is a single RNA-guided endonuclease of a class 2 CRISPR-Cas system. Cell 163, 759–771 (2015).

36 Mimee, M., Tucker, A. C., Voigt, C. A. & Lu, T. K. Programming a human commensal bacterium, Bacteroides thetaiotaomicron, to sense and respond to stimuli in the murine gut microbiota. Cell systems 1, 62–71 (2015).

37 Whitaker, W. R., Shepherd, E. S. & Sonnenburg, J. L. Tunable expression tools enable single-cell strain distinction in the gut microbiome. Cell 169, 538–546 (2017).

38 Lim, B., Zimmermann, M., Barry, N. A. & Goodman, A. L. Engineered regulatory systems modulate gene expression of human commensals in the gut. Cell 169, 547–558 (2017).

39 Benitez-Paez, A., Gomez Del Pulgar, E. M. & Sanz, Y. The glycolytic versatility of Bacteroides uniformis CECT 7771 and its genome response to oligo and polysaccharides. Frontiers in Cellular and Infection Microbiology 7, 383 (2017).

40 Li, M., Shang, Q., Li, G., Wang, X. & Yu, G. Degradation of marine algae-derived carbohydrates by Bacteroidetes isolated from human gut microbiota. Marine Drugs 15, 92 (2017).

41 Pluvinage, B. et al. Molecular basis of an agarose metabolic pathway acquired by a human intestinal symbiont. Nature Communications 9, 1043 (2018).

42 Bacic, M. K. & Smith, C. J. Laboratory maintenance and cultivation of Bacteroides species. Current Protocols in Microbiology 9, 13C–11 (2008).

43 Venturelli, O. S. et al. Deciphering microbial interactions in synthetic human gut microbiome communities. Molecular Systems Biology 14 (2018).

44 Bågenholm, V. et al. Galactomannan catabolism conferred by a polysaccharide utilization locus of Bacteroides ovatus. Journal of Biological Chemistry 292, 229–243 (2017).

45 Martens, E. C., Koropatkin, N. M., Smith, T. J. & Gordon, J. I. Complex glycan catabolism by the human gut microbiota: the Bacteroidetes Sus-like paradigm. Journal of Biological Chemistry 284, 24673–24677 (2009).

46 Almagro Armenteros, J. J. et al. SignalP 5.0 improves signal peptide predictions using deep neural networks. Nature Biotechnology 37, 420–423 (2019).

47 Yu, C. S. et al. CELLO2GO: a web server for protein subCELlular LOcalization prediction with functional gene ontology annotation. PloS one 9, e99368 (2014).

48 Yu, N. Y. et al. PSORTb 3.0: improved protein subcellular localization prediction with refined localization subcategories and predictive capabilities for all prokaryotes. Bioinformatics 26, 1608–1615 (2010).

49 Bhasin, M., Garg, A. & Raghava, G. P. PSLpred: prediction of subcellular localization of bacterial proteins. Bioinformatics 21, 2522–2524 (2005).

50 Kelley, L. A., Mezulis, S., Yates, C. M., Wass, M. N. & Sternberg, M. J. The Phyre2 web portal for protein modeling, prediction and analysis. Nature Protocols 10, 845–858 (2015).

51 Pasolli, E. et al. Extensive unexplored human microbiome diversity revealed by over 150,000 genomes from metagenomes spanning age, geography, and lifestyle. Cell 176, 649–662 (2019).

52 Almeida, A. et al. A new genomic blueprint of the human gut microbiota. Nature 568, 499-504, doi:10.1038/s41586-019-0965-1 (2019).

53 Hemsworth, G. R. et al. Structural dissection of a complex Bacteroides ovatus gene locus conferring xyloglucan metabolism in the human gut. Open Biology 6, 160142 (2016).

54 Fehlner-Peach, H. et al. Distinct polysaccharide utilization profiles of human intestinal Prevotella copri Isolates. Cell Host Microbe 26, 680–690 (2019).

55 Zhao, S. et al. Adaptive evolution within gut microbiomes of healthy people. Cell Host Microbe 25, 656–667 (2019).

56 Martens, E. C., Roth, R., Heuser, J. E. & Gordon, J. I. Coordinate regulation of glycan degradation and polysaccharide capsule biosynthesis by a prominent human gut symbiont. Journal of Biological Chemistry 284, 18445–18457 (2009).

57 Koropatkin, N. M., Cameron, E. A. & Martens, E. C. How glycan metabolism shapes the human gut microbiota. Nature Reviews Microbiology 10, 323–335 (2012).

58 Mahowald, M. A. et al. Characterizing a model human gut microbiota composed of members of its two dominant bacterial phyla. Proceedings of the National Academy of Sciences of the United States of America 106, 5859–5864 (2009).

59 Duncan, S. H. et al. Reduced dietary intake of carbohydrates by obese subjects results in decreased concentrations of butyrate and butyrate-producing bacteria in feces. Applied and Environmental Microbiology 73, 1073–1078 (2007).

60 Fischbach, M. A. & Sonnenburg, J. L. Eating for two: how metabolism establishes interspecies interactions in the gut. Cell Host Microbe 10, 336–347 (2011).

61 Makki, K., Deehan, E. C., Walter, J. & Backhed, F. The impact of dietary fiber on gut microbiota in host health and disease. Cell Host Microbe 23, 705–715 (2018).

62 Clark, R. L. et al. Design of synthetic human gut microbiome assembly and function. bioRxiv (2020).

63 Despres, J. et al. Xylan degradation by the human gut Bacteroides xylanisolvens XB1A(T) involves two distinct gene clusters that are linked at the transcriptional level. BMC Genomics 17, 326 (2016).

64 Schwalm, N. D., Townsend, G. E. & Groisman, E. A. Multiple signals govern utilization of a polysaccharide in the gut bacterium Bacteroides thetaiotaomicron. mBio 7, e01342 (2016).

65 Townsend, G. E. et al. A master regulator of Bacteroides thetaiotaomicron gut colonization controls carbohydrate utilization and an alternative protein synthesis factor. mBio 11, e03221 (2020).

66 Venturelli, O. S., Zuleta, I., Murray, R. M. & El-Samad, H. Population diversification in a yeast metabolic program promotes anticipation of environmental shifts. PLoS Biology 13, e1002042 (2015).

67 Sonnenburg, J. L. et al. Glycan foraging in vivo by an intestine-adapted bacterial symbiont. Science 307, 1955–1959 (2005).

68 Ryckman, A. E., Brockhausen, I. & Walia, J. S. Metabolism of glycosphingolipids and their role in the pathophysiology of lysosomal storage disorders. International Journal of Molecular Sciences 21, 6881 (2020).

69 Joglekar, P. et al. Intestinal IgA regulates expression of a fructan polysaccharide utilization locus in colonizing gut commensal Bacteroides thetaiotaomicron. mBio 10, e02324 (2019).

70 Reyes, A., Wu, M., McNulty, N. P., Rohwer, F. L. & Gordon, J. I. Gnotobiotic mouse model of phage-bacterial host dynamics in the human gut. Proceedings of the National Academy of Sciences of the United States of America 110, 20236–20241 (2013).

71 Rodriguez-Castano, G. P. et al. Bacteroides thetaiotaomicron starch utilization promotes quercetin degradation and butyrate production by Eubacterium ramulus. Frontiers in Microbiology 10, 1145 (2019).

72 Chia, L. W. et al. Bacteroides thetaiotaomicron fosters the growth of butyrate-producing Anaerostipes caccae in the presence of lactose and total human milk carbohydrates. Microorganisms 8, 1513 (2020).

73 Wang, J., Shoemaker, N. B., Wang, G.-R. & Salyers, A. A. Characterization of a Bacteroides mobilizable transposon, NBU2, which carries a functional lincomycin resistance gene. Journal of Bacteriology 182, 3559–3571 (2000).

74 Tagawa, J. et al. Development of a novel plasmid vector pTIO-1 adapted for electrotransformation of Porphyromonas gingivalis. Journal of Microbiological Methods 105, 174–179 (2014).

75 Datsenko & A. K. One-step inactivation of chromosomal genes in Escherichia coli K-12 using PCR products. Proceedings of the National Academy of Sciences of the United States of America 97, 6640–6645 (2000).

76 S., A. FastQC: a quality control tool for high throughput sequence data. Available online at: http://www.bioinformatics.babraham.ac.uk/projects/fastqc (2010).

77 B., B. BBMap. sourceforge.net/projects/bbmap/.

78 Liao, Y., Smyth, G. K. & Shi, W. featureCounts: an efficient general purpose program for assigning sequence reads to genomic features. Bioinformatics 30, 923–930 (2014).

79 Love, M. I., Huber, W. & Anders, S. Moderated estimation of fold change and dispersion for RNA-seq data with DESeq2. Genome Biology 15, 550 (2014).

80 Zhu, A., Ibrahim, J. G. & Love, M. I. Heavy-tailed prior distributions for sequence count data: removing the noise and preserving large differences. Bioinformatics 35, 2084–2092 (2019).

81 Jo Vandesompele, K. D. P., Filip Pattyn, Bruce Poppe, Nadine Van Roy, Anne De Paepe, Frank Speleman. Accurate normalization of real-time quantitative RT-PCR data by geometric averaging of multiple internal control genes. Genome Biology 3, 1–12 (2002).

82 Goodman, A. L. et al. Extensive personal human gut microbiota culture collections characterized and manipulated in gnotobiotic mice. Proceedings of the National Academy of Sciences of the United States of America 108, 6252–6257 (2011).

83 Zhang, J., Kobert, K., Flouri, T. & Stamatakis, A. PEAR: a fast and accurate Illumina Paired-End reAd mergeR. Bioinformatics 30, 614–620 (2014).

84 Hyatt, D. et al. Prodigal: prokaryotic gene recognition and translation initiation site identification. BMC Bioinformatics 11, 119 (2010).

85 Buchfink, B., Xie, C. & Huson, D. H. Fast and sensitive protein alignment using DIAMOND. Nature Methods 12, 59–60 (2015).

86 Zhang, H. et al. dbCAN2: a meta server for automated carbohydrate-active enzyme annotation. Nucleic Acids Res. 46, W95–W101 (2018).

87 Haario, H., Saksman, E. & Tamminen, J. An adaptive Metropolis algorithm. Bernoulli 7, 223–242 (2001).

