## Supplementary Information for "Polysaccharide utilization loci in *Bacteroides* determine population fitness and community-level interactions"

### Supplementary Note

*Development and characterization of genetic tools in B. uniformis*

To study the effects of *B. uniformis* PULs on the fitness, colonization ability and microbial interactions impacting butyrate producer growth and thus butyrate production, we needed to expand the genetic tools available for *B. uniformis*. In contrast to *E. coli*, *Bacteroides* species have a unique promoter and ribosome binding site (RBS) architectures, which could limit the generalizability of previously developed genetic tools from other bacteria in *B. uniformis*[1-4]. The genetic tools of *Bacteroides* were primarily developed in *B. thetaiotaomicron* VPI-5482[3,5], which may function differently in *B. uniformis* even though these species share similar mechanisms of gene regulation. Therefore, we systematically characterized genetic tools for manipulating gene expression in *B. uniformis* to enable predictable control of gene expression.

We first characterized a set of 18 reported RBSs from *Bacteroides* in *B. uniformis* using the strong promoter $P_{BfP1E6}$[5] driving the NanoLuc luciferase reporter. Using this promoter, our results demonstrated that RC500 and rpiL* were the weakest and strongest RBS sequences (**Fig. S1a**). Specifically, rpiL* (56,738±5267 RLU) was 203-fold higher than RC500 (279±123 RLU).

Next, we characterized 20 native *B. uniformis* promoters selected based on the -33/-7 consensus sequence (TTTG/TAnnTTTG) of *Bacteroides* promoters as well as three previously reported strong promoters in *B. thetaiotaomicron* $P_{cfiA}$, $P_{BT1311}$ and $P_{BfP1E6}$ in *B. uniformis*[2,6,7]. In addition, we characterized the promoter activities of three native tRNA promoters from *B. uniformis* ($P_{BU18065}$, $P_{BU18270}$ and $P_{BU15675}$) fused to RBS8. The $P_{BfP1E6}$ promoter and tRNA promoters $P_{BU18065}$ and $P_{BU15675}$ displayed higher promoter activities compared to other tested promoters (**Fig. S1b**). These results demonstrated that the strong promoter $P_{BfP1E6}$ previously characterized in *B. thetaiotaomicron* also has high promoter activity in *B. uniformis*. In addition, the tRNA promoters are also good candidates to achieve high expression levels in *B. uniformis*. Specifically, $P_{BfP1E6}$ (19,741±1669 RLU) was 256-fold higher than $P_{BU08215}$ (77±61 RLU).

Inducible gene expression systems are useful to optimize a target function or temporally control gene expression. To build inducible gene expression systems in *B. uniformis*, we characterized IPTG, aTc and rhamnose inducible systems[3,7]. All the inducible systems were constructed using pNBU2-ermGb and thus are integrated onto the chromosome (**Method**). To evaluate the basal and maximum expression from each inducible system, we used an empty plasmid pNBU2-ermGb and the $P_{BT1311}$ promoter driving NanoLuc as negative and positive controls, respectively. The IPTG, aTc and rhamnose inducible systems previously characterized in *B. thetaiotamicron* had undetectable basal expression in the absence of the inducers and exhibited dynamic ranges of 17 to 25-fold (**Fig. S1c-e**). As a reference, the maximum expression levels were 14-21% of the strength of the $P_{BT1311}$ promoter.

Next, we characterized a set of predicted inducible promoters from *B. uniformis* including rhamnose, inulin and xyloglucan. The function of the *B. uniformis* native rhamnose inducible system was restored with a modification to the sequence of the *rhaR* gene (**Methods**). The modified rhamnose inducible system consisting of the *rhaR'* mutant showed a dynamic range of 7-fold using the $P_{rha(BU)}$ promoter (**Fig. S1f, Table S1**). We found that $P_{BU06285}$ derived from *PUL17* was up-regulated in the presence of inulin or fructose regardless of the presence of glucose in the BMM media (**Fig. S1g-h**) and exhibited a dynamic range of 16 to 20-fold in inulin and fructose, respectively. Dissimilar to the other characterized promoters, $P_{BU06285}$ displayed basal expression in the absence of inducers and the maximum induction of $P_{BU06285}$ was 3-4 fold higher than $P_{BT1311}$ promoter as a reference (**Fig. S1h**). The xyloglucan inducible promoter $P_{BU01890}$ was up-regulated by 7-fold in the presence of xyloglucan and was inhibited by glucose (**Fig. S1i-j**). The maximum expression of $P_{BU01890}$ was 21% of the expression level of the $P_{BT1311}$ promoter as a reference.

We constructed plasmid tools for delivery of the CRIPSR-FnCpf1 system. To this end, we built four shuttle plasmids (pFW1000, pFW2000, pFW3000 and pFW4000) which each contain the *Bacteroides* replication origin pB8-51[8] and a unique *E. coli* replication origin (R6K, P15A, pSC101ts or ColE1) (**Fig. S2a**). While the four plasmids could be replicated in *B. uniformis*, these plasmids were not maintained in the absence of antibiotic selection (**Fig. S2b**). Our results showed that 40% of the cells lost the plasmid following three passages and more than 90% of the cells had lost the plasmid following nine passages (**Fig. S2b**). Therefore, the plasmids could be used for delivery of the CRIPSR-FnCpf1 based genome editing system since they were maintained in the presence of the antibiotic and then rapidly eliminated in the absence of the antibiotic.

In addition, to the deletion of diverse PULs in *B. uniformis* and the *tdk* gene in *B. uniformis* (**Fig. S3b**), we used the CRISPR-FnCpf1 method for gene fragment insertions by modifying the inactive *rhaR* gene to rescue its function with the functional sequence *rhaR'* (**Fig. S1f**). In addition, we introduced *gfp* into *PUL13* to yield a fluorescently labeled strain (**Fig. S3c**). Further, CRISPR-FnCpf1 was used to delete the xylulose kinase (*xylK*) in *Bacteroides fragilis* (**Fig. S3d**) and the levan utilization pathway (BT1754-BT1765) in *B. thetaiotaomicron* (**Fig. S3e**), demonstrating that this tool can be used for genome engineering of diverse *Bacteroides* species.

*Predicted biochemical models of glycan utilization*

Based on our experimental data demonstrating the effect of PUL deletions on the fitness of *B. uniformis* in response to different glycans (**Fig. 2b, c, d**), we propose biochemical models of glycan utilization based on bioinformatic tools and previously published studies (**Supplementary Data 1**)[9-14].

The gene cluster *PUL12* encodes 23 genes including two pairs of SusC/SusD homologs, two sugar binding proteins (SBP-1 and SBP-2), an acetyl xylan esterase-like enzyme (CE7), a cellobiose phosphorylase (CBP) and glycoside hydrolases (GH26, GH5-1, GH5-2, GH2, GH97, GH3-1 and GH3-2)[13]. GH26 is predicted to be an endo-β-1,4-mannanase and shares protein structure similarity with a surface-exposed GH26 β-mannanase in *Bacteroides ovatus*[15]. In addition, GH26 contains a lipoprotein signal peptide, thus it is predicted to be anchored on the outer cell membrane. GH5-1 is predicted to be an endo-β-1,4-glucanase, which shares protein structure similarity with a periplasmic *Cytophaga hutchinsonii* endoglucanases[16]. In addition, GH5-1 contains a lipoprotein signal peptide and thus is predicted to reside in the inner membrane and release cellobiose from oligosaccharides. GH5-2 is predicted to encode a β-1,4-mannosidase[17] and contains a lipoprotein signal peptide. Therefore, GH5-2 is predicted to reside in the inner membrane and releases mannose from the oligosaccharides. CE7 is predicted to encode an acetyl xylan esterase-like enzyme based on protein structure[18,19]. Although no signal peptide is predicted, we predict that CE7 releases acetyl groups from the side chains of oligosaccharides into the periplasm. GH3-1 and GH3-2 are predicted to encode β-glucosidases and both contain a secretory signal peptide cleaved by signal peptidase I. GH3-1 and GH3-2 share protein structure similarity with the *B. ovatus* xyloglucan PUL GH3B[20], and thus are predicted to release glucose from oligosaccharides in the periplasm. GH2 is predicted to encode β-galactosidase and contains a secretory signal peptide cleaved by signal peptidase I. GH97 is predicted to encode a α-glucosidase and contains a secretory signal peptide cleaved by signal peptidase I. GH97 shares protein structure similarity with BT1871 from *B. thetaiotaomicron*[21]. While we predict that GH2 and GH97 function in the periplasm based on their signal peptides, their functional activities are unknown.

We propose a biochemical model for glucomannan utilization (**Fig. S10**). The long chain of glucomannan is first degraded into oligosaccharides by a surface-exposed endo-1,4-mannanase (GH26, *BACUNI_RS03125*), facilitated by two sugar binding proteins (SBP-1 and SBP-2). The generated oligosaccharides are transported into the periplasm through SusC/SusD. The acetyl groups on the oligosaccharides are released by an acetyl xylan esterase-like enzyme CE7 (*BACUNI_RS03160*). Two β-glucosidases GH3-1 (*BACUNI_RS03200*) and GH3-2 (*BACUNI_RS03205*) release glucose from the oligosaccharides fragment. The endo-1,4-β-glucanase (GH5-1, *BACUNI_RS03150*) and β-1,4-mannosidase (GH5-2, *BACUNI_RS03155*) release cellobiose and mannose from the oligosaccharides and then imported into the cytoplasm. Cellobiose can be further degraded by a cellobiose phosphorylase (CBP, *BACUNI_RS03175*).

*PUL17* shares high similarity with the reported levan utilization loci in *B. thetaiotamicron* (BT1754-BT1765)[22]. *PUL17* encodes 11 genes including a SusC/SusD homolog pair, a hybrid two-component system (HTCS), a fructokinase, an inner membrane monosaccharide importer and glycoside hydrolases (GH32-1, GH32-2, GH32-3 and GH32-4)[13]. GH32-1 has protein structure similarity to β-fructofuranosidase from *Bifidobacterium longum*, which can liberate β-D-fructofuranose from non-reducing terminus of sucrose, fructooligosaccharides or inulin[23]. GH32-2 and GH32-3 share protein structure similarity features to exo-inulinase from *Aspergillus awamori*[24], which can liberate fructose from the end of the polysaccharide chain. The glycoside hydrolases GH32-1, GH32-2 and GH32-3 all contain secretory signal peptides cleaved by signal peptidase I, and thus are predicted to function in the periplasm. GH32-4 shares a similar protein structure to an endo-inulinase from *Aspergillus ficuum*[25] and BT1760 from *B. thetaiotamicron*[22], and contains a lipoprotein signal peptide suggesting that it resides on the outer cell membrane. We showed that *B. uniformis* can use both inulin and levan (**Fig. S9**) and thus assumed that GH32-4 can cleave both inulin and levan outside the cell.

A predicted biochemical model for inulin and levan utilization is shown in **Fig. S11**. In this model, a versatile outer membrane anchoring endo-inulinase GH32-4 (*BACUNI_RS06315*) can bind and degrade both inulin and levan into oligosaccharides and fructose (**Fig. S26**). The oligosaccharides are transported into periplasm through a SusC/SusD pair and are degraded by exo-inulinases GH32-2 (*BACUNI_RS06300*) and GH32-3 (*BACUNI_RS06310*) to release fructose. The remaining sucrose will be hydrolyzed by β-fructofuranosidase (GH32-1, *BACUNI_RS06295*) into fructose and glucose. A hybrid two-component system (HTCS) located on the inner membrane can sense fructose and upregulate *PUL17* for glycan utilization, consistent with our results showing that the $P_{BU06285}$ inducible promoter was up-regulated in the presence of both fructose and inulin (**Fig. S1h**).

*PUL18* shares similarity with the previously reported *B. thetaiotamicron* starch and pullulan utilization loci (both PULs contain multiple GH13)[26,27], but *PUL18* also contains a unique GH53 (*BACUNI_RS06505*, endo-β-1,4-galactanase). Therefore, we propose that the presence of this enzyme enables *PUL18* to degrade multiple glycans including glycogen, pullulan, pectic galactan and pectin (and potentially type II mucin) (**Fig. 2d**). *PUL18* encodes 10 genes including a pair of SusC/SusD, a sugar binding protein (SBP), a sugar transporter and glycoside hydrolases (GH65, GH53, GH2, GH13-1 and GH13-2)[13]. GH65 is predicted to encode a maltose phosphorylase, which shares protein structure similarity with the maltose phosphorylase from *Lactobacillus brevis*[28]. GH65 does not contain a known signal peptide, and thus is predicted to function in the cytoplasm. GH53 is predicted to be an endo-β-1,4-

galactanase, which shares high structure similarity with BT4668 from *B. thetaiotaomicron* encoding an outer membrane associated endo-β-1,4-galactanase[29,30]. GH53 contains a lipoprotein signal peptide, and thus is predicted to be anchored on the outer membrane and generate oligogalactan from the side chain of pectic galactan or pectin. GH2 is predicted to encode a β-galactosidase, which shares structure similarity with BUGH2Awt from *B. uniformis* NP1[31] and BT4668 from *B. thetaiotaomicron*[30]. GH2 contains a secretory signal peptide cleaved by signal peptidase I and is thus predicted to function in the periplasm and release galactose from the oligosaccharides. GH13-1 is predicted to be a α-amylase and shares protein structure similarity with the maltogenic amylase from a *Thermus* species[32]. GH13-1 contains a lipoprotein signal peptide. GH13-2 is predicted to be a pullulanase, and shares protein structure similarity with pullulanase from *Bacillus acidopullulyticus*[33]. GH13-2 contains a lipoprotein signal peptide. But, the sub-cellular locations of GH13-1 and GH13-2 are unknown.

A predicted model for glycogen, pullulan and pectic galactan utilization is shown in **Fig. S12**. This proposed biochemical model shows that GH13-2 (*BACUNI_RS06535*, pullulanase) and GH53 are key enzymes for utilization of diverse glycans. Specifically, GH13-2 targets glycogen and pullulan to generate oligosaccharides whereas GH53 targets the galactan side chain of pectic galactan (or pectin) to generate oligogalactan. All the oligosaccharides are transported into periplasm through a SusC/SusD pair. The oligosaccharides generated by GH13-2 are further hydrolyzed by α-amylase (GH13-1, *BACUNI_RS06530*) to generate glucose and maltose. Maltose will be transported into cytoplasm and further degraded by maltose phosphorylase (GH65, *BACUNI_RS06490*) into glucose and glucose 1-phosphate. The oligogalactan generated by GH53 will be transported into the periplasm and hydrolyzed by β-galactosidase (GH2, *BACUNI_RS06510*) into galactose.

We found that *PUL11* and *PUL43* can each support the growth of *B. uniformis* on xyloglucan, indicating that these PULs are functionally redundant for xyloglucan utilization (**Fig. 3b**). The two PULs contain 25 genes including 2 SusC/SusD pairs, 2 hybrid two-component systems (HTCS), a sugar binding protein (SBP) and 16 glycoside hydrolases[13]. GH43-1 is predicted as a non-reducing end α-L-arabinofuranosidase, which has high protein structure similarity to GH43A from *B. ovatus*[20,34]. Therefore, GH43-1 is predicted to release arabinose from oligosaccharides in the periplasm. GH5-1 is predicted to be a xyloglucanase, which has high protein structure similarity with the outer membrane anchoring GH5 from *B. ovatus*[20,34]. Thus, GH5-1 is predicted to cleave xyloglucan into oligosaccharides on the outer membrane. GH31-1 is predicted to be a α-xylosidase and shares high protein structure similarity to GH31 from *B. ovatus*[20,34]. GH31-1 is predicted to release xylose from the side chain of the xyloglucan fragment on the inner membrane. GH2-1 is predicted to be an exo-acting β-D-galactosidase. Similar to GH2 in *B. ovatus*[13,26], GH2-1 releases galactose from the side chain of the xyloglucan on the inner membrane. GH3-1 is predicted to be an isoprimeverose-producing enzyme and shares high protein structure similarity to the isoprimeverose-producing enzyme from *Aspergillus oryzae*[35]. GH3-1 contains a secretory signal peptide cleaved by signal peptidase I. Thus, GH3-1 is predicted to release isoprimeverose from xyloglucan breakdown products in the periplasm. The isoprimeverose can be further processed by GH31-1 to generate glucose and xylose. GH95 is predicted to be a α-L-galactosidase and shares similarity in protein structure to the xylan debranching enzyme GH95 in *B. ovatus*[36] but its function is not known. GH43-2 is predicted to be a non-reducing end α-L-arabinofuranosidase and shares similarity in protein structure to GH43B in the xyloglucan PUL of *B. ovatus*[20]. GH43-2 contains a lipoprotein signal peptide and is thus predicted to release arabinose from the side chain of xyloglucan breakdown products on the inner membrane. GH2-2 is predicted to be a β-galactosidase and contains a secretory signal peptide cleaved by signal peptidase I. GH2-2 is predicted to release galactose from the side chain of the xyloglucan breakdown products in the periplasm. GH5-2 is predicted to be a xyloglucanase and shares similarity to the outer membrane anchoring enzyme GH5 from *B. ovatus*[20,34]. GH5-2 is predicted to cleave xyloglucan into oligosaccharides on the outer membrane. GH97 is predicted to be a α-glucosidase and shares similarity to GH97 from *B. thetaiotaomicron*[21,37] but has unknown function. GH2-3 is predicted to be a β-galactosidase and shares similarity in protein structure with the same enzyme in *E. coli*[38]. GH2-3 contains a secretory signal peptide cleaved by signal peptidase I and thus is predicted to release galactose from the side chain of the xyloglucan breakdown products in the periplasm. GH42 is predicted to be a β-1,6-galactosidase and shares similarity in protein structure to β-galactosidase from *Bifidobacterium bifidum*[39] but has unknown function. GH2-4 is predicted to be a β-galactosidase and shares similarity in protein structure to β-galactosidase from *Thermotoga maritima*[40]. GH2-4 contains a secretory signal peptide cleaved by signal peptidase I and is thus predicted to release galactose from the side chain of the xyloglucan breakdown products in the periplasm. GH29 is predicted to be a α-fucosidase and shares similarity in protein structure to α-L-fucosidase from *Fusarium graminearum*[41]. GH29 contains a secretory signal peptide cleaved by signal peptidase I and is predicted to release fucose from the side chain of the xyloglucan breakdown products in the periplasm. GH31-2 is predicted to be a α-xylosidase and shares similarity in protein structure to GH31 from the xyloglucan PUL in *B. ovatus*[20]. GH31-2 contains a secretory signal peptide cleaved by signal peptidase I and is predicted to release xylose from the side chain of the xyloglucan breakdown products in the periplasm. GH43-3 is predicted to be a α-L-arabinofuranosidase and has similarity to arabinoxylan arabinofuranohydrolase from *Bacillus* *subtilis*[42]. GH43-3 is predicted to release arabinose from the side chain of the xyloglucan breakdown products in the periplasm.

Based on this information and a previously published paper, we propose a biochemical model for xyloglucan utilization by *PUL11* and *PUL43*[34] (**Supplementary Data 1**). For *PUL11*, xyloglucan is first degraded by GH5-1 (*BACUNI_RS01890*, xyloglucanase) into oligosaccharides. The oligosaccharides are transported into the periplasm and further degraded by GH2-1 (*BACUNI_RS01905*, exo-acting β-D-galactosidase), GH3-1 (*BACUNI_RS01910*, isoprimeverose-producing enzyme), GH31-1 (*BACUNI_RS01900*, α-xylosidase) and GH43-1 (*BACUNI_RS01885*, α-L-arabinofuranosidase) to release galactose, isoprimeverose, xylose and arabinose.

For *PUL43*, GH5-2 (*BACUNI_RS15355*, xyloglucanase) cleaves xyloglucan into oligosaccharides on the outer membrane. The oligosaccharides are transported into periplasm through SusC/SusD. The enzymes GH2-2 (*BACUNI_RS15330*, β-galactosidase), GH2-3 (*BACUNI_RS15365*, β-galactosidase) and GH2-4 (*BACUNI_RS15375*, β-galactosidase) release galactose from the side chain of the imported oligosaccharides. The enzymes GH29 (*BACUNI_RS15380*, α-fucosidase) and GH31-2 (*BACUNI_RS15385*, α-xylosidase) release fucose and xylose from oligosaccharides, respectively. The enzymes GH43-2 (*BACUNI_RS15325*, arabinanase) and GH43-3 (*BACUNI_RS15390*, α-L-arabinofuranosidases) release arabinose from the side chain of the oligosaccharides.

| **Strains** | **Genotype and description** | **Source** |
|---|---|---|
| *Bacteroides uniformis* DSM 6597 | Wild-type strain | DSMZ |
| *Bacteroides fragilis* DSM 2151 | Wild-type strain | DSMZ |
| *Bacteroides thetaiotaomicron* ATCC 29148 (VPI-5482) | Wild-type strain | ATCC |
| *Anaerostipes caccae* DSM 14662 | wild-type strain | DSMZ |
| *Coprococcus comes* ATCC 27758 | Wild-type strain | ATCC |
| *Eubacterium rectale* ATCC 33656 | Wild-type strain | ATCC |
| *Roseburia intestinalis* DSMZ 14610 | Wild-type strain | DSMZ |
| *E. coli* DH5α | $F^-$, φ80d*lac*ZΔM1, Δ(*lacZYA-argF*)U169, *deoR*, *recA*1, *endA*1, *hsdR*17($r_k^-$, $m_k^+$), *phoA*, *supE*44, $\lambda^-$*thi*-1, *gyrA*96, *relA*1 | Lab stock |
| *E. coli* pir2 | F- Δ*lac169 rpoS*(*Am*) *robA1 creC510 hsdR514 endA recA1 uidA*(Δ*MluI*)::*pir* | Invitrogen |
| *E. coli* BW29427 | *RP4-2 (TetS,kan1360::FRT*), *thrB1004*, *ΔlacZ 58*(M15), *ΔdapA1341::*[*erm pir*$^+$], *rpsL*(strR), *thi-*, *hsdS-*, *pro-* | The Coli Genetic Stock Center (CGSC) |
| BU (ermGb) | *B. uniformis* harboring plasmid pNBU2-ermGb | This study |
| BU (001) | *B. uniformis* harboring plasmid pFW001, IPTG inducible expression of NanoLuc luciferase reporter | This study |
| BU (002) | *B. uniformis* harboring plasmid pFW002, NanoLuc expression controlled by P_BfP1E6 promoter with RBS8 | This study |
| BU (003) | *B. uniformis* harboring plasmid spFW003, NanoLuc expression controlled by the $P_{cfiA}$ promoter | This study |
| BU (553) | *B. uniformis* harboring plasmid pMM553, NanoLuc expression controlled by the $P_{BT1311}$ promoter | This study |
| BU RhaR' (022) | *B. unformis* harboring RhaR' on plasmid | This study |

| | | |
|---|---|---|
| | pFW022, NanoLuc expression controlled by rhamnose inducible promoter $P_{rha(BU)}$ | |
| BU (023) | *B. uniformis* harboring plasmid pFW023, NanoLuc expression controlled by rhamnose inducible promoter $P_{rha(BT)}$ | This study |
| BU (027) | *B. uniformis* harboring plasmid pFW027, NanoLuc expression controlled by P_BfP1E6 promoter with RBS GH022 | This study |
| BU (028) | *B. uniformis* harboring plasmid pFW028, NanoLuc expression controlled by $P_{BfP1E6}$ promoter with RBS GH023 | This study |
| BU (029) | *B. uniformis* harboring plasmid pFW029, NanoLuc expression controlled by $P_{BfP1E6}$ promoter with RBS GH078 | This study |
| BU (030) | *B. uniformis* harboring plasmid pFW030, NanoLuc expression controlled by $P_{BfP1E6}$ promoter with RBS RC500 | This study |
| BU (031) | *B. uniformis* harboring plasmid pFW031, NanoLuc expression controlled by $P_{BfP1E6}$ promoter with RBS rpiL* | This study |
| BU (032) | *B. uniformis* harboring plasmid pFW032, NanoLuc expression controlled by $P_{BfP1E6}$ promoter with RBS A21 | This study |
| BU (033) | *B. uniformis* harboring plasmid pFW033, NanoLuc expression controlled by $P_{BfP1E6}$ promoter with RBS B1 | This study |
| BU (034) | *B. uniformis* harboring plasmid pFW034, NanoLuc expression controlled by $P_{BfP1E6}$ promoter with RBS B41 | This study |
| BU (035) | *B. uniformis* harboring plasmid pFW035, NanoLuc expression controlled by $P_{BfP1E6}$ promoter with RBS B40 | This study |
| BU (036) | *B. uniformis* harboring plasmid | This study |

| | | |
|---|---|---|
| | pFW036, NanoLuc expression controlled by $P_{BfP1E6}$ promoter with RBS C56 | |
| BU (037) | *B. uniformis* harboring plasmid pFW037, NanoLuc expression controlled by $P_{BfP1E6}$ promoter with RBS1 | This study |
| BU (038) | *B. uniformis* harboring plasmid pFW038, NanoLuc expression controlled by $P_{BfP1E6}$ promoter with RBS2 | This study |
| BU (039) | *B. uniformis* harboring plasmid pFW039, NanoLuc expression controlled by $P_{BfP1E6}$ promoter with RBS3 | This study |
| BU (040) | *B. uniformis* harboring plasmid pFW040, NanoLuc expression controlled by $P_{BfP1E6}$ promoter with RBS4 | This study |
| BU (041) | *B. uniformis* harboring plasmid pFW041, NanoLuc expression controlled by $P_{BfP1E6}$ promoter with RBS5 | This study |
| BU (042) | *B. uniformis* harboring plasmid pFW042, NanoLuc expression controlled by $P_{BfP1E6}$ promoter with RBS6 | This study |
| BU (043) | *B. uniformis* harboring plasmid pFW043, NanoLuc expression controlled by $P_{BfP1E6}$ promoter with RBS7 | This study |
| BU (044) | *B. uniformis* harboring plasmid pFW044, NanoLuc expression controlled by $P_{BU18075}$ promoter | This study |
| BU (045) | *B. uniformis* harboring plasmid pFW045, NanoLuc expression controlled by $P_{BU18080}$ promoter | This study |
| BU (046) | *B. uniformis* harboring plasmid pFW046, NanoLuc expression controlled by $P_{BU15585}$ promoter | This study |
| BU (047) | *B. uniformis* harboring plasmid pFW047, NanoLuc expression controlled by $P_{recO}$ promoter | This study |
| BU (048) | *B. uniformis* harboring plasmid | This study |

| | | |
|---|---|---|
| | pFW048, NanoLuc expression controlled by $P_{BU12125}$ promoter | |
| BU (049) | *B. uniformis* harboring plasmid pFW049, NanoLuc expression controlled by $P_{BU16445}$ promoter | This study |
| BU (050) | *B. uniformis* harboring plasmid pFW050, NanoLuc expression controlled by $P_{gyrB}$ promoter | This study |
| BU (051) | *B. uniformis* harboring plasmid pFW051, NanoLuc expression controlled by $P_{BU13840}$ promoter | This study |
| BU (052) | *B. uniformis* harboring plasmid pFW052, NanoLuc expression controlled by $P_{BU09950}$ promoter | This study |
| BU (053) | *B. uniformis* harboring plasmid pFW053, NanoLuc expression controlled by $P_{BU10945}$ promoter | This study |
| BU (054) | *B. uniformis* harboring plasmid pFW054, NanoLuc expression controlled by $P_{BU02520}$ promoter | This study |
| BU (055) | *B. uniformis* harboring plasmid pFW055, NanoLuc expression controlled by $P_{BU05295}$ promoter | This study |
| BU (056) | *B. uniformis* harboring plasmid pFW056, NanoLuc expression controlled by $P_{BU08215}$ promoter | This study |
| BU (057) | *B. uniformis* harboring plasmid pFW057, NanoLuc expression controlled by $P_{BU08545}$ promoter | This study |
| BU (058) | *B. uniformis* harboring plasmid pFW058, NanoLuc expression controlled by $P_{BU11495}$ promoter | This study |
| BU (059) | *B. uniformis* harboring plasmid pFW059, NanoLuc expression controlled by $P_{BU00025}$ promoter | This study |
| BU (060) | *B. uniformis* harboring plasmid pFW060, NanoLuc expression controlled by $P_{BU02380}$ promoter | This study |
| BU (061) | *B. uniformis* harboring plasmid pFW061, NanoLuc expression controlled by $P_{BU18065}$ promoter with RBS8 | This study |
| BU (062) | *B. uniformis* harboring plasmid pFW062, NanoLuc expression | This study |

| | | |
|---|---|---|
| | controlled by $P_{BU18270}$ promoter with RBS8 | |
| BU (063) | *B. uniformis* harboring plasmid pFW063, NanoLuc expression controlled by $P_{BU15675}$ promoter with RBS8 | This study |
| BU (064) | *B. uniformis* harboring plasmid pFW064, aTc inducible expression of NanoLuc | This study |
| BU (104) | *B. uniformis* harboring plasmid pFW104, NanoLuc expression controlled by $P_{BU06285}$ promoter | This study |
| BU (106) | *B. uniformis* harboring plasmid pFW106, NanoLuc expression controlled by $P_{BU01890}$ promoter | This study |
| BU Δ*tdk* | *B. uniformis* with the deletion of *tdk* gene | This study |
| BU Δ*tyrP-1* | *B. uniformis*, a barcode was integrated onto the chromosome while deleting the *tyrP* gene. Barcode: CGTG | This study |
| BU Δ*tyrP-2* | *B. uniformis*, a barcode was integrated onto the chromosome while deleting the *tyrP* gene. Barcode: GGGG | This study |
| BU Δ*tyrP-3* | *B. uniformis*, a barcode was integrated onto the chromosome while deleting the *tyrP* gene. Barcode: AATG | This study |
| BU Δ*tyrP-4* | *B. uniformis*, a barcode was integrated onto the chromosome while deleting the *tyrP* gene. Barcode: AGCC | This study |
| BU Δ*tyrP-5* | *B. uniformis*, a barcode was integrated onto the chromosome while deleting the *tyrP* gene. Barcode: GAGT | This study |
| BU Δ*tyrP-6* | *B. uniformis*, a barcode was integrated onto the chromosome while deleting the *tyrP* gene. Barcode: GGTG | This study |
| BU Δ*tyrP-7* | *B. uniformis*, a barcode was integrated onto the chromosome while deleting the *tyrP* gene. Barcode: AGTG | This study |
| BU Δ*tyrP-8* | *B. uniformis*, a barcode was integrated onto the chromosome while deleting the *tyrP* gene. Barcode: TGGC | This study |
| BU Δ*tyrP-11* | *B. uniformis*, a barcode was integrated onto the chromosome while deleting the *tyrP* gene. Barcode: GAGG | This study |

| | | |
|---|---|---|
| BU Δ*tyrP-12* | *B. uniformis*, a barcode was integrated onto the chromosome while deleting the *tyrP* gene. Barcode: CAGG | This study |
| BU Δ*tyrP-13* | *B. uniformis*, a barcode was integrated onto the chromosome while deleting the *tyrP* gene. Barcode: GCAG | This study |
| BU Δ*tyrP-14* | *B. uniformis*, a barcode was integrated onto the chromosome while deleting the *tyrP* gene. Barcode: GAGC | This study |
| BU Δ*tyrP-16* | *B. uniformis*, a barcode was integrated onto the chromosome while deleting the *tyrP* gene. Barcode: AGGG | This study |
| BU Δ*tyrP-17* | *B. uniformis*, a barcode was integrated onto the chromosome while deleting the *tyrP* gene. Barcode: TTAA | This study |
| BU Δ*tyrP-18* | *B. uniformis*, a barcode was integrated onto the chromosome while deleting the *tyrP* gene. Barcode: TAAG | This study |
| BU Δ*tyrP-19* | *B. uniformis*, a barcode was integrated onto the chromosome while deleting the *tyrP* gene. Barcode: GGTC | This study |
| BU Δ*tyrP-20* | *B. uniformis*, a barcode was integrated onto the chromosome while deleting the *tyrP* gene. Barcode: TTTG | This study |
| BU Δ*tyrP-21* | *B. uniformis*, a barcode was integrated onto the chromosome while deleting the *tyrP* gene. Barcode: TAGC | This study |
| BU Δ*tyrP-22* | *B. uniformis*, a barcode was integrated onto the chromosome while deleting the *tyrP* gene. Barcode: TCTA | This study |
| BU Δ*tyrP-24* | *B. uniformis*, a barcode was integrated onto the chromosome while deleting the *tyrP* gene. Barcode: CGGG | This study |
| BU Δ*PUL6* | *B. uniformis* Δ*tyrP-1* combined with deletion of *PUL6*, deletion region: *BACUNI_RS02580-BACUNI_RS02620* | This study |
| BU Δ*PUL7* | *B. uniformis* Δ*tyrP-2* combined with deletion of *PUL7*, deletion region: *BACUNI_RS02680-BACUNI_RS02750* | This study |
| BU Δ*PUL11* | *B. uniformis* Δ*tyrP-3* combined with deletion of *PUL11*, deletion region: *BACUNI_RS01865-BACUNI_RS01915* | This study |
| BU Δ*PUL18* | *B. uniformis* Δ*tyrP-4* combined with deletion of *PUL18*, deletion region: | This study |

| | | |
|---|---|---|
| | *BACUNI_RS06490-BACUNI_RS06535* | |
| BU Δ*PUL13* | *B. uniformis* Δ*tyrP-5* combined with deletion of *PUL13*, deletion region: *BACUNI_RS05300-BACUNI_RS05330* | This study |
| BU Δ*PUL16* | *B. uniformis* Δ*tyrP-6* combined with deletion of *PUL16*, deletion region: *BACUNI_RS05980-BACUNI_RS06030* | This study |
| BU Δ*PUL17* | *B. uniformis* Δ*tyrP-7* combined with deletion of *PUL17*, deletion region: *BACUNI_RS06285-BACUNI_RS06315* | This study |
| BU Δ*PUL54* | *B. uniformis* Δ*tyrP-8* combined with deletion of *PUL54*, deletion region: *BACUNI_RS18165-BACUNI_RS18230* | This study |
| BU Δ*PUL34* | *B. uniformis PUL34* deletion with barcode TGAC inserted onto the chromosome, deletion region: *BACUNI_RS14295-BACUNI_RS14325* | This study |
| BU Δ*PUL49* | *B. uniformis PUL49* deletion with barcode TATG inserted onto the chromosome, deletion region: *BACUNI_RS16190-BACUNI_RS16225* | This study |
| BU Δ*PUL21* | *B. uniformis* Δ*tyrP-11* with the deletion of *PUL21*, deletion region: *BACUNI_RS07655-BACUNI_RS07680* | This study |
| BU Δ*PUL22_23* | *B. uniformis* Δ*tyrP-12* with the deletion of *PUL22* and *PUL23*, deletion region: *BACUNI_RS08140-BACUNI_RS08190* | This study |
| BU Δ*PUL24* | *B. uniformis* Δ*tyrP-13* combined with deletion of *PUL24*, deletion region: *BACUNI_RS09510-BACUNI_RS09520* | This study |
| BU Δ*PUL28* | *B. uniformis* Δ*tyrP-14* combined with deletion of *PUL28*, deletion region: *BACUNI_RS12235-BACUNI_RS12255* | This study |
| BU Δ*PUL32* | *B. uniformis* Δ*tyrP-16* combined with deletion of *PUL32*, deletion region: *BACUNI_RS14085-BACUNI_RS14125* | This study |
| BU Δ*PUL35* | *B. uniformis* Δ*tyrP-17* combined with deletion of *PUL35*, deletion region: *BACUNI_RS14825-BACUNI_RS14860* | This study |
| BU Δ*PUL37* | *B. uniformis* Δ*tyrP-18* combined with deletion of *PUL37*, deletion region: *BACUNI_RS00220-BACUNI_RS00250* | This study |
| BU Δ*PUL43* | *B. uniformis* Δ*tyrP-19* combined with deletion of *PUL43*, deletion region: | This study |

| | | |
|---|---|---|
| | *BACUNI_RS15325-BACUNI_RS15390* | |
| BU Δ*PUL48* | *B. uniformis* Δ*tyrP-20* combined with deletion of *PUL48*, deletion region: *BACUNI_RS15960-BACUNI_RS15995* | This study |
| BU Δ*PUL44* | *B. uniformis* Δ*tyrP-21* combined with deletion of *PUL44*, deletion region: *BACUNI_RS15425-BACUNI_RS15460* | This study |
| BU Δ*PUL47* | *B. uniformis* Δ*tyrP-22* combined with deletion of *PUL47*, deletion region: *BACUNI_RS15760-BACUNI_RS15795* | This study |
| BU Δ*PUL12* | *B. uniformis PUL12* deletion with barcode TCGG inserted onto the chromosome, deletion region: *BACUNI_RS03125-BACUNI_RS03185* | This study |
| BU Δ*PUL11_43* | *B. uniformis* Δ*tyrP-3* combined with the double deletion of *PUL11* and *PUL43* | This study |
| BU Δ*PUL11_44* | *B. uniformis* Δ*tyrP-3* combined with double deletion of *PUL11* and *PUL44* | This study |
| BU RhaR' | *B. uniformis* with the modification of its inactive *rhaR* gene | This study |
| BU Δ*PUL13*::g*fp* | *B. uniformis*, *gfp* insertion onto the chromosome combined with deletion of *PUL13* | This study |
| BU (1000) | *B. uniformis* harboring shuttle plasmid pFW1000 | This study |
| BU (2000) | *B. uniformis* harboring shuttle plasmid pFW2000 | This study |
| BU (3000) | *B. uniformis* harboring shuttle plasmid pFW3000 | This study |
| BU (4000) | *B. uniformis* harboring shuttle plasmid pFW4000 | This study |
| BF Δ*xyl* | *B. fragilis* with the deletion of xylulose kinase gene | This study |
| BT Δ*levan* | *B. thetaiotamicron* with the deletion of levan utilization cluster, deletion region: *BT1754-BT1765* | This study |

**Table S1. Strains used in this study.**

| Plasmid | Description | Relevant features | References |
|---|---|---|---|
| pNBU2-ermGb | pNBU2-bla-ermGb | *Bacteriodes* plasmid, $Erm^r$, $Amp^r$ | [26] |
| pMM553 | pNBU2-$P_{BT1311}$-NL | NanoLuc constitutively from $P_{BT1311}$, pNBU2-ermGb backbone | Addgene (#68543)[3] |
| pMM656 | pNBU2-$P_{rha\ (BT)}$-NL | Rhamnose inducible NanoLuc expression, pNBU2-ermGb backbone | Addgene (#68887)[3] |
| pMM733 | pNBU2-BtdCas9-NL4 | IPTG inducible CRISPRi vector targeting NanoLuc (NL4), constitutive NanoLuc, pNBU2-ermGb backbone | Addgene (#68898)[3] |
| pNBU2_erm-TetR-P1T_DP-GH023 | | aTc inducible gene expression cassette, pNBU2-ermGb backbone | Addgene[7] (#90324) |
| pWW3452 | AmpR-ermG-RP4/R6K -[P_BfP1E6-RBSlp-LP-GFPtag-Term]-NBU2 | sfGFP constitutive expression plasmid, pNBU2-ermGb backbone | [5] |
| pT7FnCpf1 | | Fncpf1 controlled by T7 promoter | Lab stock |
| pFW001 | pNBU2-$P_{lacO23}$-NL-$P_{BT1311}$-*lacI* | IPTG inducible NanoLuc expression, pNBU2-ermGb backbone | This study |
| pFW002 | pNBU2-P_BfP1E6-NL | NanoLuc expressed constitutively from $P_{BfP1E6}$ with RBS8, pNBU2-ermGb backbone | This study |
| pFW003 | pNBU2-$P_{cfiA}$-NL | NanoLuc expressed constitutively from $P_{cfiA}$, pNBU2-ermGb backbone | This study |
| pFW018 | pNBU2-$P_{BT1311}$-*lacI* | $P_{BT1311}$-lacI cassette, pNBU2-ermGb backbone | This study |
| pFW022 | pNBU2-$P_{rha(BU)}$-NL | NanoLuc expressed from rhamnose inducible promoter $P_{rha(BU)}$ from BU, pNBU2-ermGb backbone | This study |
| pFW023 | pNBU2-$P_{BT1311}$-*rhaR*-$P_{rha(BT)}$-NL | NanoLuc expressed from rhamnose inducible promoter $P_{rha(BT)}$ and *rhaR* regulator from BT, pMM656 derivatives | This study |
| pFW025 | pNBU2-$P_{BT1311}$-*lacI*-$P_{lacO23}$ | pNBU2-ermGb with IPTG induction cassette | This study |
| pFW027 | pNBU2-P_BfP1E6-GH022-NL | NanoLuc expressed constitutively from $P_{BfP1E6}$ promoter with RBS GH022, pNBU2-ermGb backbone | This study |
| pFW028 | pNBU2-P_BfP1E6-GH023-NL | NanoLuc expressed constitutively from $P_{BfP1E6}$ with RBS GH023, pNBU2-ermGb backbone | This study |

| | | | |
|---|---|---|---|
| pFW029 | pNBU2-P_BfP1E6-GH078-NL | NanoLuc expressed constitutively from $P_{BfP1E6}$ with RBS GH078, pNBU2-ermGb backbone | This study |
| pFW030 | pNBU2-P_BfP1E6-RC500-NL | NanoLuc expressed constitutively from $P_{BfP1E6}$ with RBS RC500, pNBU2-ermGb backbone | This study |
| pFW031 | pNBU2-P_BfP1E6-rpiL*-NL | NanoLuc expressed constitutively from $P_{BfP1E6}$ with RBS rpiL*, pNBU2-ermGb backbone | This study |
| pFW032 | pNBU2-P_BfP1E6-A21-NL | NanoLuc expressed constitutively from $P_{BfP1E6}$ with RBS A21, pNBU2-ermGb backbone | This study |
| pFW033 | pNBU2-P_BfP1E6-B1-NL | NanoLuc expressed constitutively from $P_{BfP1E6}$ with RBS B1, pNBU2-ermGb backbone | This study |
| pFW034 | pNBU2-P_BfP1E6-B41-NL | NanoLuc expressed constitutively from $P_{BfP1E6}$ with RBS B41, pNBU2-ermGb backbone | This study |
| pFW035 | pNBU2-P_BfP1E6-B40-NL | NanoLuc expressed constitutively from $P_{BfP1E6}$ with RBS B40, pNBU2-ermGb backbone | This study |
| pFW036 | pNBU2-P_BfP1E6-C56-NL | NanoLuc expressed constitutively from $P_{BfP1E6}$ with RBS C56, pNBU2-ermGb backbone | This study |
| pFW037 | pNBU2-P_BfP1E6-RBS1-NL | NanoLuc expressed constitutively from $P_{BfP1E6}$ with RBS1, pNBU2-ermGb backbone | This study |
| pFW038 | pNBU2-P_BfP1E6-RBS2-NL | NanoLuc expressed constitutively from $P_{BfP1E6}$ with RBS2, pNBU2-ermGb backbone | This study |
| pFW039 | pNBU2-P_BfP1E6-RBS3-NL | NanoLuc expressed constitutively from $P_{BfP1E6}$ with RBS3, pNBU2-ermGb backbone | This study |
| pFW040 | pNBU2-P_BfP1E6-RBS4-NL | NanoLuc expressed constitutively from $P_{BfP1E6}$ with RBS4, pNBU2-ermGb backbone | This study |
| pFW041 | pNBU2-P_BfP1E6-RBS5-NL | NanoLuc expressed constitutively from $P_{BfP1E6}$ with RBS5, pNBU2-ermGb backbone | This study |
| pFW042 | pNBU2-P_BfP1E6-RBS6-NL | NanoLuc expressed constitutively from $P_{BfP1E6}$ with RBS6, pNBU2-ermGb backbone | This study |

| | | | |
|---|---|---|---|
| pFW043 | pNBU2-P_BfP1E6-RBS7-NL | NanoLuc expressed constitutively from $P_{BfP1E6}$ with RBS7, pNBU2-ermGb backbone | This study |
| pFW044 | pNBU2-$P_{BU18075}$-NL | NanoLuc expressed from $P_{BU18075}$, pNBU2-ermGb backbone | This study |
| pFW045 | pNBU2-$P_{BU18080}$-NL | NanoLuc expressed from $P_{BU18080}$, pNBU2 backbone | This study |
| pFW046 | pNBU2-$P_{BU15585}$-NL | NanoLuc expressed from $P_{BU15585}$, pNBU2-ermGb backbone | This study |
| pFW047 | pNBU2-$P_{recO}$-NL | NanoLuc expressed from $P_{recO}$, pNBU2-ermGb backbone | This study |
| pFW048 | pNBU2-$P_{BU12125}$-NL | NanoLuc expressed from $P_{BU12125}$, pNBU2-ermGb backbone | This study |
| pFW049 | pNBU2-$P_{BU16445}$-NL | NanoLuc expressed from $P_{BU16445}$, pNBU2-ermGb backbone | This study |
| pFW050 | pNBU2-$P_{gyrB}$-NL | NanoLuc expressed from $P_{gyrB}$, pNBU2-ermGb backbone | This study |
| pFW051 | pNBU2-$P_{BU13840}$-NL | NanoLuc expressed from $P_{BU13840}$, pNBU2-ermGb backbone | This study |
| pFW052 | pNBU2-$P_{BU09950}$-NL | NanoLuc expressed from $P_{BU09950}$, pNBU2-ermGb backbone | This study |
| pFW053 | pNBU2-$P_{BU10945}$-NL | NanoLuc expressed from $P_{BU10945}$, pNBU2-ermGb backbone | This study |
| pFW054 | pNBU2-$P_{BU02520}$-NL | NanoLuc expressed from $P_{BU02520}$, pNBU2-ermGb backbone | This study |
| pFW055 | pNBU2-$P_{BU05295}$-NL | NanoLuc expressed from $P_{BU05295}$, pNBU2-ermGb backbone | This study |
| pFW056 | pNBU2-$P_{BU08215}$-NL | NanoLuc expressed from $P_{BU08215}$, pNBU2-ermGb backbone | This study |
| pFW057 | pNBU2-$P_{BU08545}$-NL | NanoLuc expressed from $P_{BU08545}$, pNBU2-ermGb backbone | This study |
| pFW058 | pNBU2-$P_{BU11495}$-NL | NanoLuc expressed from $P_{BU11495}$, pNBU2-ermGb backbone | This study |
| pFW059 | pNBU2-$P_{BU00025}$-NL | NanoLuc expressed from $P_{BU00025}$, pNBU2-ermGb backbone | This study |
| pFW060 | pNBU2-$P_{BU02380}$-NL | NanoLuc expressed from $P_{BU02380}$, pNBU2-ermGb backbone | This study |
| pFW061 | pNBU2-$P_{BU18065}$-RBS8-NL | NanoLuc expressed from $P_{BU18065}$-RBS8, pNBU2-ermGb backbone | This study |
| pFW062 | pNBU2-$P_{BU18270}$-RBS8-NL | NanoLuc expressed from $P_{BU18270}$-RBS8, pNBU2-ermGb backbone | This study |
| pFW063 | pNBU2-$P_{BU15675}$-RBS8-NL | NanoLuc expressed from $P_{BU15675}$-RBS8, pNBU2-ermGb backbone | This study |

| | | | |
|---|---|---|---|
| pFW064 | pNBU2-TetR-P1T-NL | aTc inducible expression of NanoLuc, pNBU2_erm-TetR-P1T_DP-GH023 derivative | This study |
| pFW104 | pNBU2-$P_{BU06285}$-NL | NanoLuc expressed from $P_{BU06285}$, pNBU2-ermGb backbone | This study |
| pFW106 | pNBU2-$P_{BU01890}$-NL | NanoLuc expressed from $P_{BU01890}$, pNBU2-ermGb backbone | This study |
| pFW1000 | *Bacteriodes-E. coli* shuttle plasmid | *Bacteroides-E. coli* shuttle plasmid, Erm$^r$, Amp$^r$, R6K origin, minimal pB8-51 origin (1651 bp)[8] | This study |
| pFW2000 | *Bacteriodes-E. coli* shuttle plasmid | pFW1000 derivative with R6K origin replaced by p15A | This study |
| pFW3000 | *Bacteriodes-E. coli* shuttle plasmid | pFW1000 derivative with R6K origin replaced by pSC101ts | This study |
| pFW4000 | *Bacteriodes-E. coli* shuttle plasmid | pFW1000 derivative with R6K origin replaced by ColE1 | This study |
| pFW1001 | pFW1000-$P_{BT1311}$-*lacI*-$P_{lacO23}$ | pFW1000 with IPTG inducible genetic cassette | This study |
| pFW1004 | pFW1000-$P_{BT1311}$-*lacI*-$P_{lacO23}$-*Fncpf1* | *Fncpf1* expressed from the IPTG inducible promoter, pFW1000 backbone | This study |
| pFW2001 | pFW2000- $P_{BT1311}$-*lacI*-$P_{lacO23}$ | pFW2000 with IPTG inducible cassette | This study |
| pFW2100 | pFW2000-$P_{BT1311}$-*lacI*-$P_{lacO23}$-*Fncpf1* | *Fncpf1* expressed by the IPTG inducible promoter, pFW2000 backbone | This study |
| pFW2500 | pFW2100-$P_{lacO23}$-*recT* | *Fncpf1* and *recT* expressed from the IPTG inducible promoter, pFW2000 backbone | This study |
| pFW1007 | pFW1004-P_BfP1E6-crRNA-*xyl*-UD | *B. fragilis* xylulose kinase gene deletion from the $P_{BfP1E6}$ controlled crRNA, pFW1004 backbone | This study |
| pFW1008 | pFW1004-$P_{BT1311}$-crRNA -*tdk*-UD | *B. uniformis tdk* gene deletion with $P_{BT1311}$ controlled crRNA expression, pFW1004 backbone | This study |
| pFW2026 | pFW2100-$P_{BT1311}$-crRNA-*tryP*-UD-Barcode | 4-bp barcode (NNNN) on the chromosome while deleting the *tryP* gene. $P_{BT1311}$ controlled crRNA expression, pFW2100 backbone | This study |
| pFW2028 | pFW2100-$P_{BT1311}$-crRNA-*RhaR'*-UD | *B. uniformis RhaR* modification. $P_{BT1311}$ controlled crRNA expression, pFW2100 backbone | This study |
| pFW2029 | pFW2100-$P_{BT1311}$-crRNA -*PUL6*-UD | *B. uniformis PUL6* deletion with $P_{BT1311}$ controlled crRNA expression, pFW2100 backbone | This study |

| | | | |
|---|---|---|---|
| pFW2030 | pFW2100-$P_{BT1311}$-crRNA -*PUL7*-UD | *B. uniformis PUL7* deletion with $P_{BT1311}$ controlled crRNA expression, pFW2100 backbone | This study |
| pFW2031 | pFW2100-$P_{BT1311}$-crRNA -*PUL11*-UD | *B. uniformis PUL11* deletion with $P_{BT1311}$ controlled crRNA expression, pFW2100 backbone | This study |
| pFW2057 | pFW2100-$P_{BT1311}$-crRNA -*PUL12*-UD | *B. uniformis PUL12* deletion with $P_{BT1311}$ controlled crRNA expression, pFW2100 backbone | This study |
| pFW2033 | pFW2100-$P_{BT1311}$-crRNA -*PUL13*-UD | *B. uniformis PUL13* deletion with $P_{BT1311}$ controlled crRNA expression, pFW2100 backbone | This study |
| pFW2034 | pFW2100-$P_{BT1311}$-crRNA -*PUL16*-UD | *B. uniformis PUL16* deletion with $P_{BT1311}$ controlled crRNA expression, pFW2100 backbone | This study |
| pFW2035 | pFW2100-$P_{BT1311}$-crRNA -*PUL17*-UD | *B. uniformis PUL17* deletion with $P_{BT1311}$ controlled crRNA expression, pFW2100 backbone | This study |
| pFW2073 | pFW2500-$P_{BT1311}$-crRNA -*PUL18*-UD | *B. uniformis PUL18* deletion with $P_{BT1311}$ controlled crRNA expression, pFW2500 backbone | This study |
| pFW2061 | pFW2100-$P_{BT1311}$-2crRNAs-*PUL21*-UD | *B. uniformis PUL21* deletion with $P_{BT1311}$ controlled crRNAs expression, pFW2100 backbone | This study |
| pFW2074 | pFW2500-$P_{BT1311}$-crRNA -*PUL22-23*-UD | *B. uniformis PUL22* and *PUL23* deletion with $P_{BT1311}$ controlled crRNA expression, pFW2500 backbone | This study |
| pFW2075 | pFW2500-$P_{BT1311}$-2crRNAs-*PUL24*-UD | *B. uniformis PUL24* deletion with $P_{BT1311}$ controlled crRNAs expression, pFW2500 backbone | This study |
| pFW2076 | pFW2500-$P_{BT1311}$-2crRNAs-*PUL28*-UD | *B. uniformis PUL28* deletion with $P_{BT1311}$ controlled crRNAs expression, pFW2500 backbone | This study |
| pFW2066 | pFW2500-$P_{BT1311}$-2crRNAs-*PUL32*-UD | *B. uniformis PUL32* deletion with $P_{BT1311}$ controlled crRNAs expression, pFW2500 backbone | This study |
| pFW2043 | pFW2100-$P_{BT1311}$-crRNA -*PUL34*-UD | *B. uniformis PUL34* deletion with $P_{BT1311}$ controlled crRNA expression, pFW2100 backbone | This study |
| pFW2067 | pFW2500-$P_{BT1311}$-2crRNAs-*PUL35*-UD | *B. uniformis PUL35* deletion with $P_{BT1311}$ controlled crRNAs expression, pFW2500 backbone | This study |

| | | | |
|---|---|---|---|
| pFW2068 | pFW2500-$P_{BT1311}$-2crRNAs-*PUL37*-UD | *B. uniformis PUL37* deletion with $P_{BT1311}$ controlled crRNAs expression, pFW2500 backbone | This study |
| pFW2069 | pFW2500-$P_{BT1311}$-2crRNAs-*PUL43*-UD | *B. uniformis PUL43* deletion with $P_{BT1311}$ controlled crRNAs expression, pFW2500 backbone | This study |
| pFW2070 | pFW2500-$P_{BT1311}$-2crRNAs-*PUL44*-UD | *B. uniformis PUL44* deletion with $P_{BT1311}$ controlled crRNAs expression, pFW2500 backbone | This study |
| pFW2071 | pFW2500-$P_{BT1311}$-2crRNAs-*PUL47*-UD | *B. uniformis PUL47* deletion with $P_{BT1311}$ controlled crRNAs expression, pFW2500 backbone | This study |
| pFW2072 | pFW2500-$P_{BT1311}$-2crRNAs-*PUL48*-UD | *B. uniformis PUL48* deletion with $P_{BT1311}$ controlled crRNAs expression, pFW2500 backbone | This study |
| pFW2050 | pFW2100-$P_{BT1311}$-crRNA -*PUL49*-UD | *B. uniformis PUL49* deletion with $P_{BT1311}$ controlled crRNA expression, pFW2100 backbone | This study |
| pFW2051 | pFW2100-$P_{BT1311}$-crRNA -*PUL54*-UD | *B. uniformis PUL54* deletion with $P_{BT1311}$ controlled crRNA expression, pFW2100 backbone | This study |
| pFW2059 | pFW2100-$P_{BT1311}$-crRNA -*PUL13::gfp*-UD | *gfp* insertion with *PUL13* been deleted. $P_{BT1311}$ controlled crRNA expression, pFW2100 backbone | This study |
| pFW2078 | pFW2500-$P_{BT1311}$-crRNA -*Levan*-UD | *B. thetaiotamicron levan* utilization cluster deletion with $P_{BT1311}$ controlled crRNA expression, pFW2500 backbone | This study |
| pFW2100-*tyrP*-24 | pFW2100-$P_{BT1311}$-crRNA -*tyrP-24*-UD | *B. uniformis tyrP-24* deletion with $P_{BT1311}$ controlled crRNA expression, barcode (CGGG), pFW2100 backbone | This study |
| pFW2500-*tyrP*-24 | pFW2500-$P_{BT1311}$-crRNA -*tyrP-24*-UD | *B. uniformis tyrP-24* deletion with $P_{BT1311}$ controlled crRNA expression, barcode (CGGG), pFW2500 backbone | This study |

**Table S2. Plasmids used in this study.**

| **Strains** | **Carbon source** | **Co-culture of butyrate producer and *B. uniformis*** | ***B. uniformis* conditioned media** | ***B. uniformis* cell membrane treated glycan media** |
|---|---|---|---|---|
| *A. caccae* | Glucose | 0.90±0.08 | 1.25±0.06 | NA |
| | Glycogen | 0.67±0.29 | 1.72±0.08 | 6.82±0.07 |
| | Glucomannan | 0.48±0.24 | 3.66±0.04 | 5.02±0.28 |
| | Inulin | 9.69±0.66 | 21.26±3.37 | 57.70±5.72 |
| | Laminarin | 8.24±1.67 | 7.17±1.51 | 45.10±8.01 |
| | Pectic galactan | 1.08±0.37 | 2.31±0.02 | 1.92±0.39 |
| | Pectin | 0.37±0.19 | 1.19±0.09 | 4.78±0.56 |
| | Pullulan | 16.73±2.63 | 13.91±0.28 | 61.76±3.30 |
| | Xyloglucan | 0.68±0.41 | 4.72±0.11 | 51.03±6.38 |
| *C. comes* | Glucose | 0.57±0.03 | 1.74±0.13 | NA |
| | Glycogen | 1.97±0.15 | 1.00±0.00 | 1.23±0.58 |
| | Glucomannan | 0.45±0.02 | 1.06±0.06 | 0.70±0.14 |
| | Inulin | 20.55±1.21 | 18.47±0.49 | 10.62±1.54 |
| | Laminarin | 13.64±1.19 | 3.39±0.07 | 13.50±0.27 |
| | Pectic galactan | 3.42±0.38 | 0.99±0.00 | 0.67±0.07 |
| | Pectin | 1.90±0.18 | 0.97±0.06 | 1.02±0.16 |
| | Pullulan | 18.77±2.37 | 10.76±0.09 | 12.29±1.14 |
| | Xyloglucan | 0.57±0.11 | 1.31±0.06 | 1.01±0.35 |
| *E. rectale* | Glucose | 1.73±0.21 | 2.18±0.40 | NA |
| | Glycogen | 1.06±0.07 | 1.24±0.08 | 1.00±0.01 |
| | Glucomannan | 0.33±0.08 | 4.36±0.30 | 2.10±0.16 |
| | Inulin | 9.55±1.01 | 7.28±0.15 | 8.46±0.42 |
| | Laminarin | 20.09±4.07 | 4.80±0.19 | 14.37±2.09 |
| | Pectic galactan | 5.87±1.44 | 10.20±0.97 | 6.51±0.45 |
| | Pectin | 1.64±0.21 | 1.07±0.15 | 1.00±0.11 |
| | Pullulan | 1.80±0.08 | 2.93±0.09 | 2.80±0.04 |
| | Xyloglucan | 0.60±0.11 | 2.94±0.06 | 13.47±1.90 |
| *R. intestinalis* | Glucose | 1.26±0.28 | 1.23±0.02 | NA |
| | Glycogen | 0.82±0.17 | 3.54±0.09 | 4.47±0.26 |
| | Glucomannan | 0.51±0.05 | 6.03±1.28 | 14.21±0.62 |
| | Inulin | 12.80±0.60 | 10.27±0.52 | 13.17±0.67 |
| | Laminarin | 8.70±2.50 | 4.04±0.13 | 34.38±0.65 |

| | | | | |
|---|---|---|---|---|
| | Pectic galactan | 3.95±0.28 | 18.25±0.02 | 9.76±1.48 |
| | Pectin | 1.08±0.14 | 1.14±0.10 | 3.09±0.53 |
| | Pullulan | 1.24±0.15 | 1.86±0.03 | 1.91±0.13 |
| | Xyloglucan | 0.81±0.13 | 18.92±0.41 | 33.25±1.43 |

**Table S3. Fold changes of butyrate producer growth modified by *B. uniformis* compared to butyrate producer growth in the corresponding fresh media.** Fold changes of the total growth were computed by dividing the area under $OD_{600}$ curve (AUC) of each butyrate producer in *B. uniformis*-butyrate producer coculture, *B. uniformis* conditioned media or *B. uniformis* cell membrane treated glycan media with the AUC of the $OD_{600}$ curve of the butyrate producer in the corresponding fresh media. For AUC calculation, when the measured $OD_{600}$ values were lower than the initial $OD_{600}$, we set the $OD_{600}$ data to the initial $OD_{600}$. All the values shown represent the mean ± 1 s.d (4-9 values from 2-3 biological replicates). For conditioned glucose media experiments, the concentration of glucose was measured and then adjusted to the concentration of glucose in the fresh media (**Methods**). NA indicates that this condition was not measured.

| Component | Concentration | Note |
| --- | --- | --- |
| **BMM-C (100 mL)** | | |
| Hemin | 200 µL | 2.5 mg $mL^{-1}$ solution |
| L-Methionine | 2 mg | |
| Mineral 3B solution | 5 mL | See recipe for Mineral 3B solution |
| L-Cysteine | 100 mg | |
| FeSO4 | 150 µL | 1.53 mg $mL^{-1}$ solution |
| Sodium bicarbonate | 2 mL | 10% solution |
| **Mineral 3B solution (100 mL)** | | |
| $KH_2PO_4$ | 1.8 g | |
| NaCl | 1.8 g | |
| $MgCl_2.6H_2O$ | 40 mg | |
| $CaCl_2$ | 39 mg | |
| $CoCl_2.6H_2O$ | 2 mg | |
| $MnCl_2.4H_2O$ | 20 mg | |
| $NH_4Cl$ | 1g | |
| $Na_2SO_4$ | 500 mg | |
| **BMM (1L)** | | |
| BMM-C | 1 L | |
| Glucose | 5 g | |
| **BMM-glycan**[##] | | |
| DM29 | 1 L | |
| Glycan | Variable (**Table S5**) | One glycan used for cell growth |

**Table S4. Media recipes for *Bacteroides* minimal media (BMM)[43], BMM-C and BMM-glycan[#].** Media was sterilized by filteration to a final pH of 7.1. [##]The concentration and preparation of the glycans are described in **Table S5**.

| Glycans | Vendor | Item Number | Concentration (g $L^{-1}$) | Sterilization |
|---|---|---|---|---|
| Arabinogalactan | Sigma | 10830-25G | 5 | filtered# |
| Arabinoxylan | Megazyme | P-WAXYL | 5 | filtered |
| Amylopectin (maize) | Sigma | 10120-250G | 5 | autoclaved## |
| Agarose | Sigma | A9539-10G | <2 | autoclaved |
| Chondroitin sulfate | Sigma | C9819-5G | 5 | filtered |
| Dextran (*Leuconostoc* spp) | Sigma | 31392-10G | 5 | filtered |
| Galactan (potato) | Sigma | 50009-1G | 5 | autoclaved |
| Galactomannan (carob) | Sigma | 91783-1G | 5 | autoclaved |
| Glucomannan | Megazyme | P-GLCML | <2 | autoclaved |
| Glycogen | Sigma | G8751-5G | 5 | filtered |
| Gum Arabic (acacia tree) | Sigma | G9752-500G | 5 | filtered |
| Inulin (chicory)### | Sigma | I2255-10G | 5 | filtered |
| Inulin (chicory) | Acros Organics | AC457100250 | 5 | filtered |
| Laminarin (*Laminaria digitata*) | Sigma | L9634-1G | 5 | filtered |
| Lichenin (Icelandic Moss) | Megazyme | P-LICHN | 2.5 | filtered |
| Levan | Megazyme | P-LEVAN | 5 | filtered |
| Pectic galactan (Potato) | Megazyme | P-PGAPT | 5 | filtered |
| Pectin | Gojira Fine Chemicals | PE1006 | 5 | autoclaved |
| Pullulan (*Aureobasidium pullulans*) | Sigma | P4516-5G | 5 | filtered |
| Starch | Sigma | S9765-100G | <2 | autoclaved |
| Type II mucin (porcine stomach) | Sigma | M2378-100G | 5 | autoclaved |
| Type III mucin (porcine stomach) | Sigma | M1778-100G | 2.5 | autoclaved |
| Xylan (Beechwood) | Megazyme | P-XYLNBE-10G | 5 | filtered |
| Xyloglucan (Tamarind) | Megazyme | P-XYGLN | 5 | autoclaved |

**Table S5. Glycans used in this study.**

#0.2 μm filter (Whatman) was used to sterilize the media.

##Supernatant of the autoclaved glycan media was used for cell growth.

###Type of inulin used for all experiments unless otherwise noted.

| Component | Concentration (mM) | Note |
|---|---|---|
| **DM29 (1 L)** | | |
| $Na_2SeO_3$ | 0.000578243 | |
| $Na_2WO_4$ | 0.003403444 | |
| $AlK(SO_4)_2$ | 0.003872891 | |
| $CaCl_2$ | 1.290106325 | |
| $CuSO_4$ | 0.021265311 | |
| $Na_2MoO_4$ | 0.004856255 | |
| $NiCl_2$ | 0.01543217 | |
| $H_3BO_3$ | 0.016172589 | |
| $Co(NO_3)_2$ | 0.054661835 | |
| $ZnSO_4$ | 0.061931009 | |
| $FeSO_4$ | 0.065829318 | |
| EDTA | 0.171092253 | |
| $MnSO_4$ | 0.331123635 | |
| NaCl | 1.711075639 | |
| $MgSO_4$ | 2.741638004 | |
| L-Cysteine | 8.4 | |
| L-Aspartic acid | 0.4 | |
| L-Glutamic acid | 0.662721893 | |
| L-Tryptophan | 0.73 | |
| L-Methionine | 0.84 | |
| L-Histidine | 1 | |
| L-Isoleucine | 1.6 | |
| L-Threonine | 1.9 | |
| L-Lysine | 2.4 | |
| L-Asparagine | 2.6 | |
| L-Glutamine | 2.7 | |
| L-Valine | 2.8 | |
| L-Leucine | 3.6 | |
| L-Phenylalanine | 4.5 | |
| L-Glycine | 4.7 | |
| L-Alanine | 5.3 | |
| L-Proline | 5.9 | |
| L-Serine | 6.4 | |
| L-Arginine | 21.81400689 | |
| Potassium Hydroxide | 0.000449153 | TekNova 10x AGCU Mix |
| Uracil | 5.88829E-05 | |
| Cytosine | 5.94059E-05 | |
| Guanine | 5.95514E-05 | |
| L-Adenine | 5.99423E-05 | |
| Orotic Acid | 0.064061499 | |

| | | |
|---|---|---|
| Magnesium Chloride | 1.680495746 | |
| Potassium Phosphate | 6.61327063 | |
| Nicotinamide | 0.034392401 | |
| Xanthine | 0.024981921 | |
| Folic Acid | 0.002402832 | |
| Pyrixodal | 0.009822218 | |
| Thymidine | 0.021 | |
| Biotin | 0.040952069 | |
| Ca-D-pantothenate | 0.000419695 | |
| Cobalamin | 1.47561E-06 | |
| Tetrahydrofolic Acid | 2.80628E-06 | |
| p-Aminobenzoic Acid | 0.073072918 | |
| Pyridoxine HCl | 0.000972583 | |
| Riboflavin | 0.000531406 | |
| Thiamine HCl | 0.000593064 | |
| Pyridoxamine | 0.021 | |
| Haemin | 0.015338835 | |
| Ammonia Chloride | 9.347366847 | |
| MOPS | 71.68003181 | |
| Sodium Bicarbonate | 47.6150797 | |
| L-Tyrosine | 3.201059661 | |
| Sodium Sulfate | 7.040270346 | |
| **Component** | **Concentration** | **Note** |
| **DM29-glucose (1 L)** | | |
| DM29 | 1 L | |
| Glucose | 5 g | |
| **DM29-glycan (1 L)** ## | | |
| DM29 | 1 L | |
| Glycan | Variable | One specific glycan used for cell growth |

**Table S6. Recipe of defined media 29 (DM29), DM29-glucose and DM29-glycan#.**
#DM29 is a media without a carbon source. This media was sterilized by filtration to a final pH of 6.7.
##Information about glycans can be found in **Table S5**.

| **Primer name** | **Index** | **Sequence** |
|---|---|---|
| BU-barcode_fwd_1 | ATCACG | AATGATACGGCGACCACCGAGATCTACACATCAC<br>GACACTCTTTCCCTACACGACGCTCTTCCGATCTT<br>GTCTAAATTCGGCACATTAAACC |
| BU-barcode_fwd_2 | CGATGT | AATGATACGGCGACCACCGAGATCTACACCGATG<br>T<br>ACACTCTTTCCCTACACGACGCTCTTCCGATCTTT<br>GTCTAAATTCGGCACATTAAACC |
| BU-barcode_fwd_3 | TTAGGC | AATGATACGGCGACCACCGAGATCTACACTTAGG<br>C<br>ACACTCTTTCCCTACACGACGCTCTTCCGATCTGT<br>TGTCTAAATTCGGCACATTAAACC |
| BU-barcode_fwd_4 | TGACCA | AATGATACGGCGACCACCGAGATCTACACTGACC<br>A<br>ACACTCTTTCCCTACACGACGCTCTTCCGATCTCG<br>A TGTCTAAATTCGGCACATTAAACC |
| BU-barcode_fwd_5 | ACAGTG | AATGATACGGCGACCACCGAGATCTACACACAGT<br>G<br>ACACTCTTTCCCTACACGACGCTCTTCCGATCTAT<br>GATGTCTAAATTCGGCACATTAAACC |
| BU-barcode_fwd_6 | GCCAAT | AATGATACGGCGACCACCGAGATCTACACGCCAA<br>T<br>ACACTCTTTCCCTACACGACGCTCTTCCGATCTTG<br>CGATGTCTAAATTCGGCACATTAAACC |
| BU-barcode_fwd_7 | CAGATC | AATGATACGGCGACCACCGAGATCTACACCAGAT<br>C<br>ACACTCTTTCCCTACACGACGCTCTTCCGATCTGA<br>GTGGTGTCTAAATTCGGCACATTAAACC |
| BU-barcode_fwd_8 | ACTTGA | AATGATACGGCGACCACCGAGATCTACACACTTGA<br>ACACTCTTTCCCTACACGACGCTCTTCCGATCTCC<br>TGGAGTGTCTAAATTCGGCACATTAAACC |
| BU-barcode_rev_1 | ATCACGAG | CAAGCAGAAGACGGCATACGAGATATCACGAGGT<br>GACTGGAGTTCAGACGTGTGCTCTTCCGATCTAGA<br>ATATCGTACGGTGGAGAAAC |
| BU-barcode_rev_2 | CGATGTTC | CAAGCAGAAGACGGCATACGAGATCGATGTTCGT<br>GACTGGAGTTCAGACGTGTGCTCTTCCGATCTAAG<br>AATATCGTACGGTGGAGAAAC |
| BU-barcode_rev_3 | TTAGGCGA | CAAGCAGAAGACGGCATACGAGATTTAGGCGAGT<br>GACTGGAGTTCAGACGTGTGCTCTTCCGATCTTCA<br>GAATATCGTACGGTGGAGAAAC |
| BU-barcode_rev_4 | TGACCAAT | CAAGCAGAAGACGGCATACGAGATTGACCAATGT<br>GACTGGAGTTCAGACGTGTGCTCTTCCGATCTCTA<br>AGAATATCGTACGGTGGAGAAAC |

| | | |
|---|---|---|
| BU-barcode_rev_5 | ACAGTGCT | CAAGCAGAAGACGGCATACGAGATACAGTGCTGTGACTGGAGTTCAGACGTGTGCTCTTCCGATCTGATAAGAATATCGTACGGTGGAGAAAC |
| BU-barcode_rev_6 | GCCAATGT | CAAGCAGAAGACGGCATACGAGATGCCAATGTGTGACTGGAGTTCAGACGTGTGCTCTTCCGATCTACTCAAGAATATCGTACGGTGGAGAAAC |
| BU-barcode_rev_7 | CAGATCGA | CAAGCAGAAGACGGCATACGAGATCAGATCGAGTGACTGGAGTTCAGACGTGTGCTCTTCCGATCTTTCTCTAGAATATCGTACGGTGGAGAAAC |
| BU-barcode_rev_8 | ACTTGAAA | CAAGCAGAAGACGGCATACGAGATACTTGAAAGTGACTGGAGTTCAGACGTGTGCTCTTCCGATCTCACTTCTAGAATATCGTACGGTGGAGAAAC |
| BU-barcode_rev_9 | GATCAGTG | CAAGCAGAAGACGGCATACGAGATGATCAGTGGTGACTGGAGTTCAGACGTGTGCTCTTCCGATCTAGAATATCGTACGGTGGAGAAAC |
| BU-barcode_rev_10 | TCTACCTC | CAAGCAGAAGACGGCATACGAGATTCTACCTCGTGACTGGAGTTCAGACGTGTGCTCTTCCGATCTAAGAATATCGTACGGTGGAGAAAC |
| BU-barcode_rev_11 | CTTGTATG | CAAGCAGAAGACGGCATACGAGATCTTGTATGGTGACTGGAGTTCAGACGTGTGCTCTTCCGATCTTCAGAATATCGTACGGTGGAGAAAC |
| BU-barcode_rev_12 | TAGCTTCC | CAAGCAGAAGACGGCATACGAGATTAGCTTCCGTGACTGGAGTTCAGACGTGTGCTCTTCCGATCTCTAAGAATATCGTACGGTGGAGAAAC |

**Table S7. Primers used for next-generation sequencing of barcoded *B. uniformis* strains.**

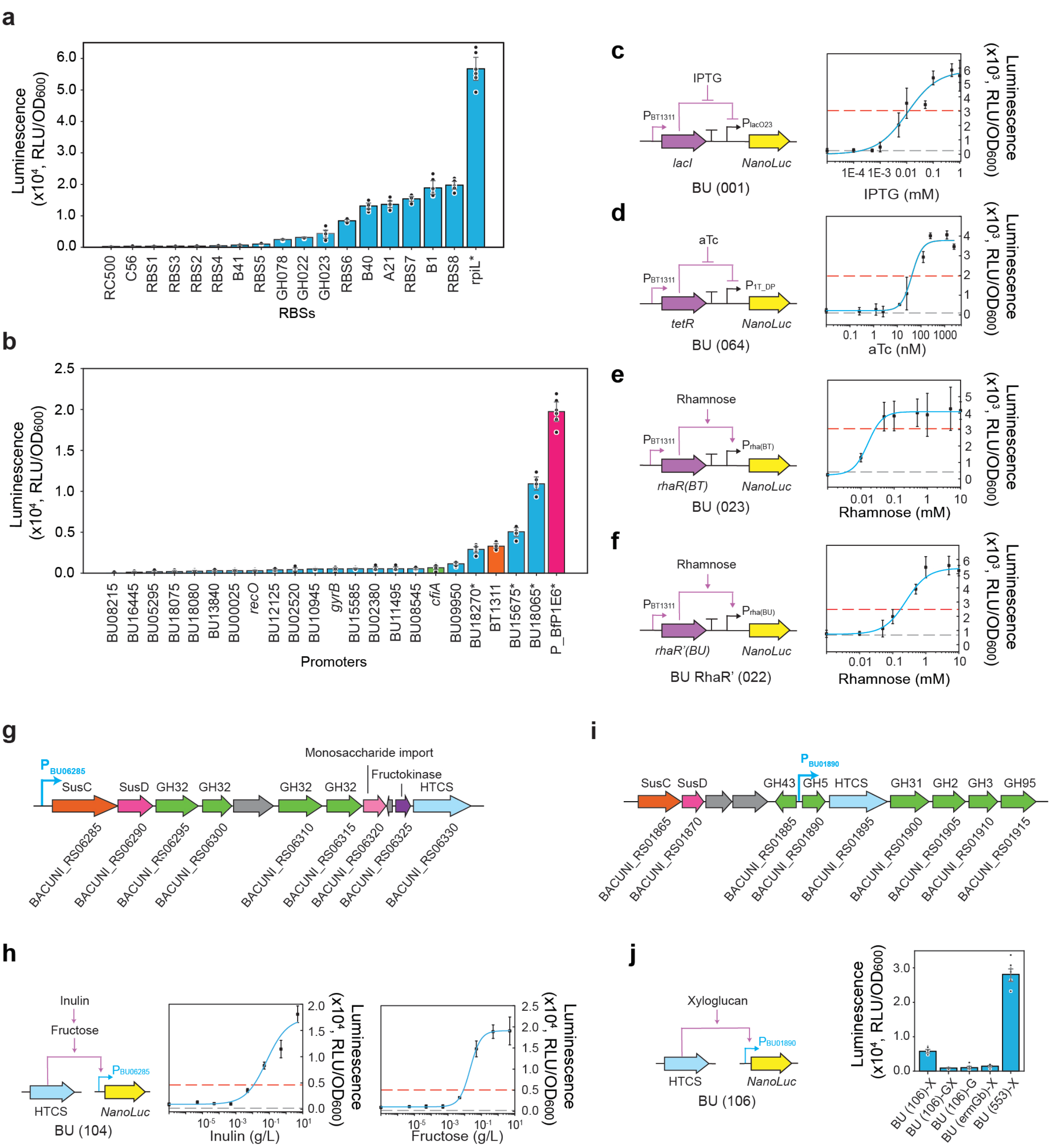

**Figure S1. Characterization of libraries of ribosome binding sites and promoters on gene expression in *B. uniformis*. (a)** Bar plot of the NanoLuc luciferase reporter activity divided by absorbance at 600 nm ($OD_{600}$) for a set of ribosome binding sites (RBSs) in *B. uniformis*. All RBSs were controlled by the strong $P_{BfP1E6}$ promoter. Data points represent 6 values from 2 biological replicates (3 technical replicates for each). Data points represent the mean and error bars denote 1 s.d. from the mean. **(b)** Bar plot of NanoLuc luciferase reporter activity divided by $OD_{600}$ for a set of promoters in *B. uniformis*. The green, orange and pink colored bars represent three previously reported strong promoters in *B. thetaiotaomicron*[3,5]. An asterisk (*) indicates RBS8 was placed under those promoters. The original RBS was used for those promoters without an asterisk. Data points represent 6 values from 2 biological replicates (3 technical replicates for each). Data points represent the mean and error bars denote 1 s.d. from the mean. **(c)** Dose response of the NanoLuc luciferase reporter activity divided by $OD_{600}$ driven by an IPTG inducible promoter as a function of IPTG concentration in *B. uniformis*. The gray dashed line indicates the background signal of the control empty plasmid (BU (ermGb)) and the red dashed lines indicates 10% of the signal obtained from the constitutive promoter $P_{BT1311}$ (BU (553)) in panels **(c)-(f).** The blue line represents a Hill function fit to the data (Hill function: $y = b + a\left(\frac{x^n}{k^n+x^n}\right)$ where $b,\ a,\ n,\ k$ represent model parameters and $y$ and $x$ denote the normalized luminescence (RLU $OD600^{-1}$) and concentration of inducer, respectively. Data points represent the mean and error bars denote 1 s.d. from the mean (n=3). **(d)** Dose response of the NanoLuc luciferase reporter activity divided by $OD_{600}$ driven by an aTc inducible promoter as a function of aTc concentration in *B. uniformis*. The blue line indicates a Hill function fit to the data. Data points represent the mean and error bars denote 1 s.d. from the mean (n=3). **(e)** Dose response of the NanoLuc luciferase reporter activity divided by $OD_{600}$ driven by a rhamnose inducible promoter ($P_{rha(BT)}$) as a function of rhamnose concentration in *B. uniformis*. The *rhaR* transcription regulator gene was cloned from *B. thetaiotaomicron*. The blue line indicates a Hill function fit to the data. Data points represent the mean and error bars denote 1 s.d. from the mean (n=3). **(f)** Dose response of the NanoLuc luciferase reporter activity divided by $OD_{600}$ driven by a rhamanose inducible promoter ($P_{rha(BU)}$) as a function of rhamnose concentration in *B. uniformis*. The inactive *rhaR* gene from *B. uniformis* was modified to generate a functional *rhaR'* gene. The blue line indicates a Hill function fit to the data. Data points represent the mean and error bars denote 1 s.d. from the mean (n=6). **(g)** Schematic of the gene organization the inulin utilization pathway in *B. uniformis*. **(h)** Dose response of the NanoLuc luciferase reporter activity divided by $OD_{600}$ driven by an inulin/fructose inducible promoter as a function of inulin (left) or fructose (right) in *B. uniformis*. The gray dashed line indicates the signal obtained from the control empty plasmid (BU (ermGb)). The red dashed line indicates the signal obtained from the constitutive promoter $P_{BT1311}$ (BU (553)). The blue line indicates a Hill function fit to the data. Data points represent the mean and error bars denote 1 s.d. from the mean (n=6). **(i)** Schematic of the gene organization of *B. uniformis* xyloglucan utilization pathway. **(j)** Dose response of the NanoLuc luciferase reporter activity divided by $OD_{600}$ driven by a xyloglucan inducible promoter as a function of xyloglucan in *B. uniformis*. BU (106)-X, BU (ermGb)-X, BU (553)-X indicate strains in BMM-xyloglucan media; BU (106)-G indicates BU (106) in BMM media; BU (106)-GX indicates BU (106) in media with both glucose and xyloglucan. Data points represent 9 values from 3 independent biological replicates (3 technical replicates for each). Data points represent the mean and error bars denote 1 s.d. from the mean.

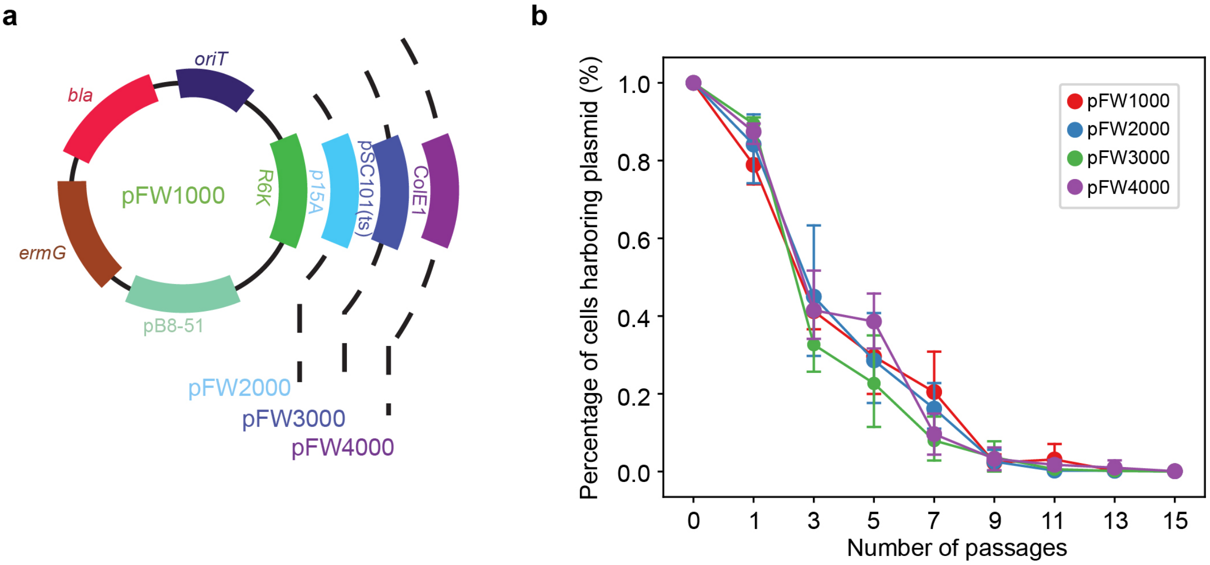

**Figure S2. Construction and characterization of *E. coli–Bacteroides* shuttle plasmids. (a)** Schematic of *E. coli-Bacteroides* shuttle plasmids. The four shuttle plasmids were designed with *ermG* (erythromycin resistant gene) for *Bacteroides* selection, pB8-51[8] as the replication origin from *Bacteroides*, *oriT* for conjugation, *bla* (carbenicillin resistant gene) for antibiotic selection in *E. coli* and one of four different *E. coli* origins of replication. **(b)** Line plot of the number of passages versus the percent of cells harboring plasmid. Data points denote the mean and error bars are 1 s.d. from the mean (n=4).

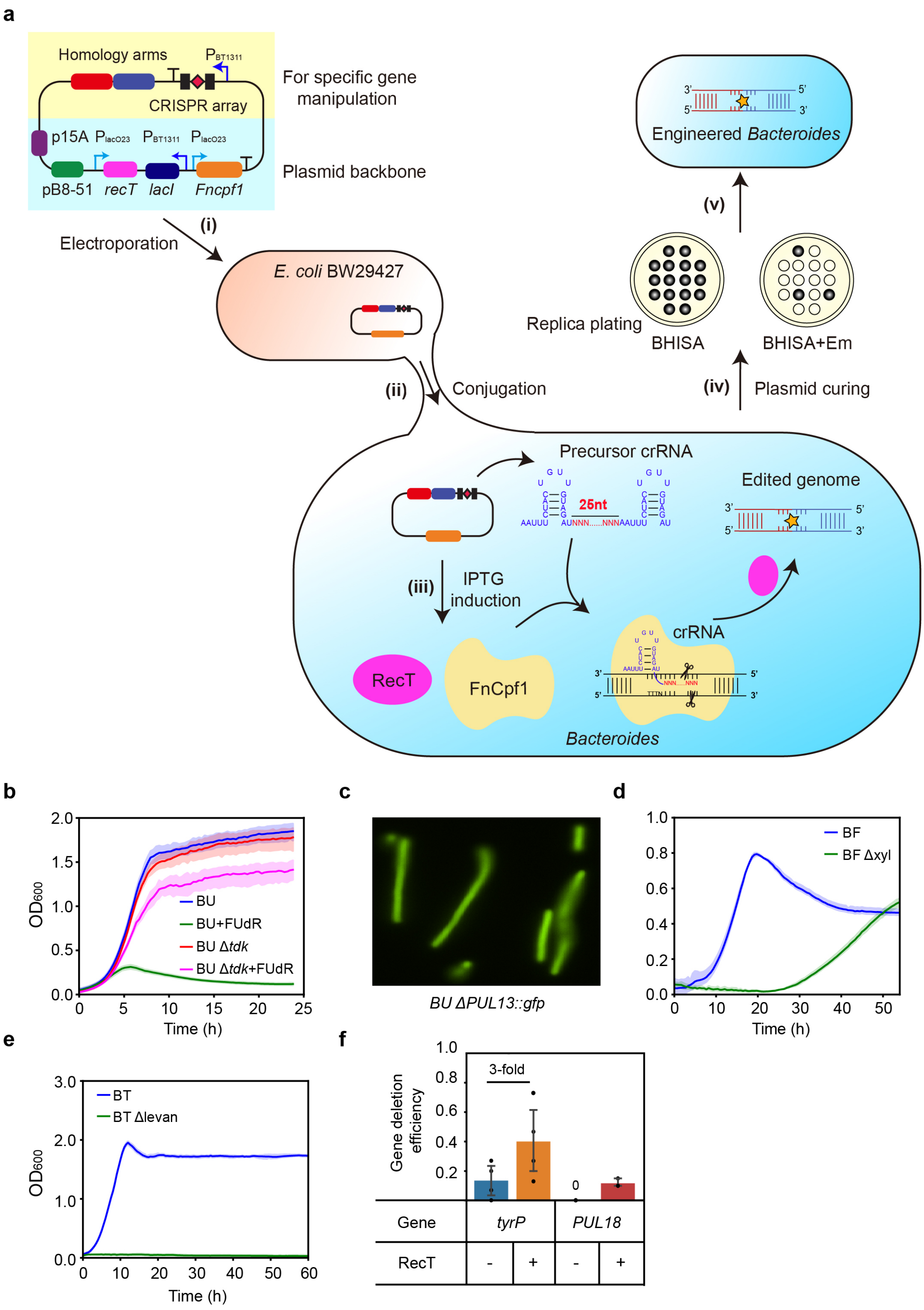

**Figure S3. Genome editing in *Bacteroides* with CRISPR-FnCpf1. (a)** Schematic of gene-manipulation procedures in *Bacteroides* (using gene deletion with pFW2500 as an example). (i) The two homology arms with crRNA expression cassette were ligated into the pFW2500 plasmid, which contains an IPTG inducible *Fncpf1* and *recT*. The sequence confirmed plasmid was transformed into *E. coli* BW29427. (ii) The plasmid was subsequently transformed into the *Bacteroides* host strain via conjugation with *E. coli* BW29427. (iii) The strain harboring the gene deletion plasmid was then induced by IPTG via plating onto the BHISA+Em+IPTG agar plate. The mutants were confirmed by colony PCR. (iv) The correct mutants are passaged in anaerobic basal broth (ABB) 3 times and then spread on BHISA agar plates. Colonies were picked and spotted on BHISA and BHISA+Em plates. The mutants that were unable to grow on the BHISA+Em plates are selected as the correct mutants. **(b)** Absorbance at 600 nm ($OD_{600}$) as a function of time for *B. uniformis* wild-type (BU) and BU Δ*tdk* in the presence and absence of FUdR (5-fluorodeoxyuridine, 200 µg/mL). Lines denote the mean and shaded regions represent 95% confidence interval. **(c)** Fluorescent microscopy image of the *B. uniformis* Δ*PUL13*::*gfp* mutant. **(d)** $OD_{600}$ as a function of time for *B. fragilis* wild-type (BF) and BF Δ*xyl* in BMM-xylose media. Lines denote the mean and shaded regions represent 95% confidence interval. **(e)** $OD_{600}$ as a function of time for *B. thetaiotaomicron* wild-type (BT) and BT Δ*levan* in BMM-Levan media. Lines denote the mean and shaded regions represent 95% confidence interval. **(f)** Bar plot of gene deletion efficiency of *tyrP* and *PUL18* with and without RecT. Data points represent biological replicates, the height of the bar represents the mean and error bars are 1 s.d. from the mean (n=3-4).

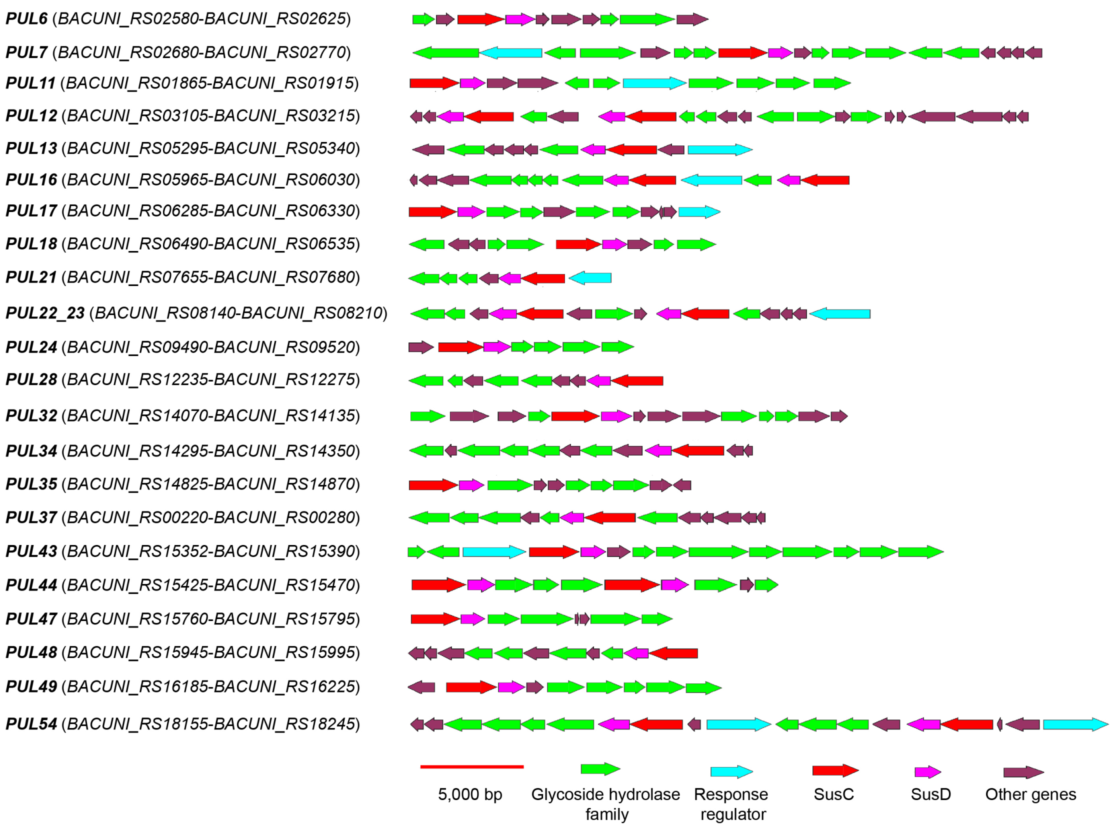

**Figure S4. Gene organization of 23 polysaccharide utilization loci targeted for deletion in *B. uniformis*.** Highlighted genes include glycoside hydrolases, response regulators, *susC* and s*usD*.

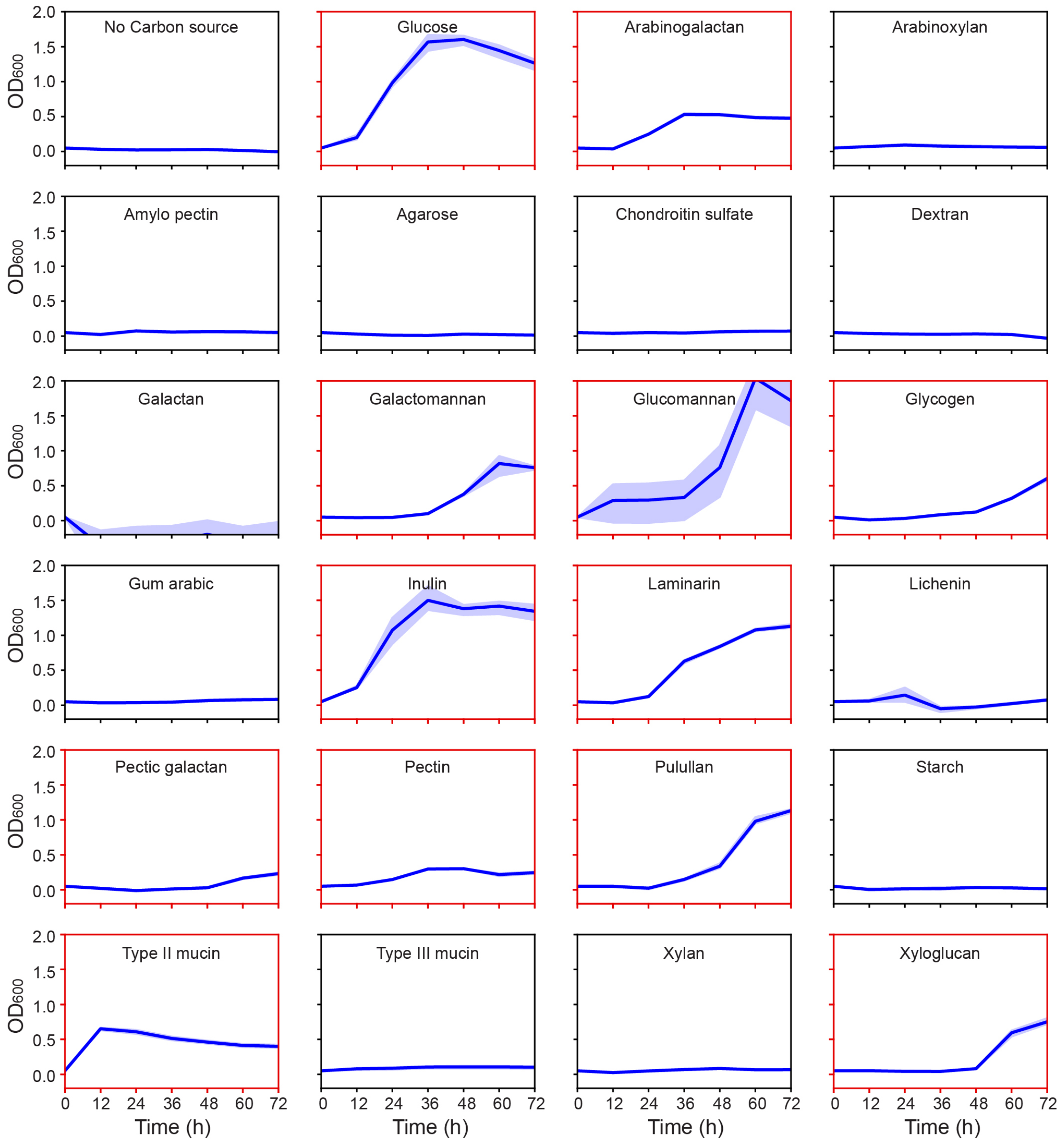

**Figure S5. *B. uniformis* growth over time in the presence of different carbon sources.** Each sub-plot represents absorbance at 600 nm ($OD_{600}$) as a function of time. Lines denote the mean and shaded regions represent 95% confidence interval (n=3). The sub-plots outlined in red indicate conditions with a maximum $OD_{600}$ greater than 0.2.

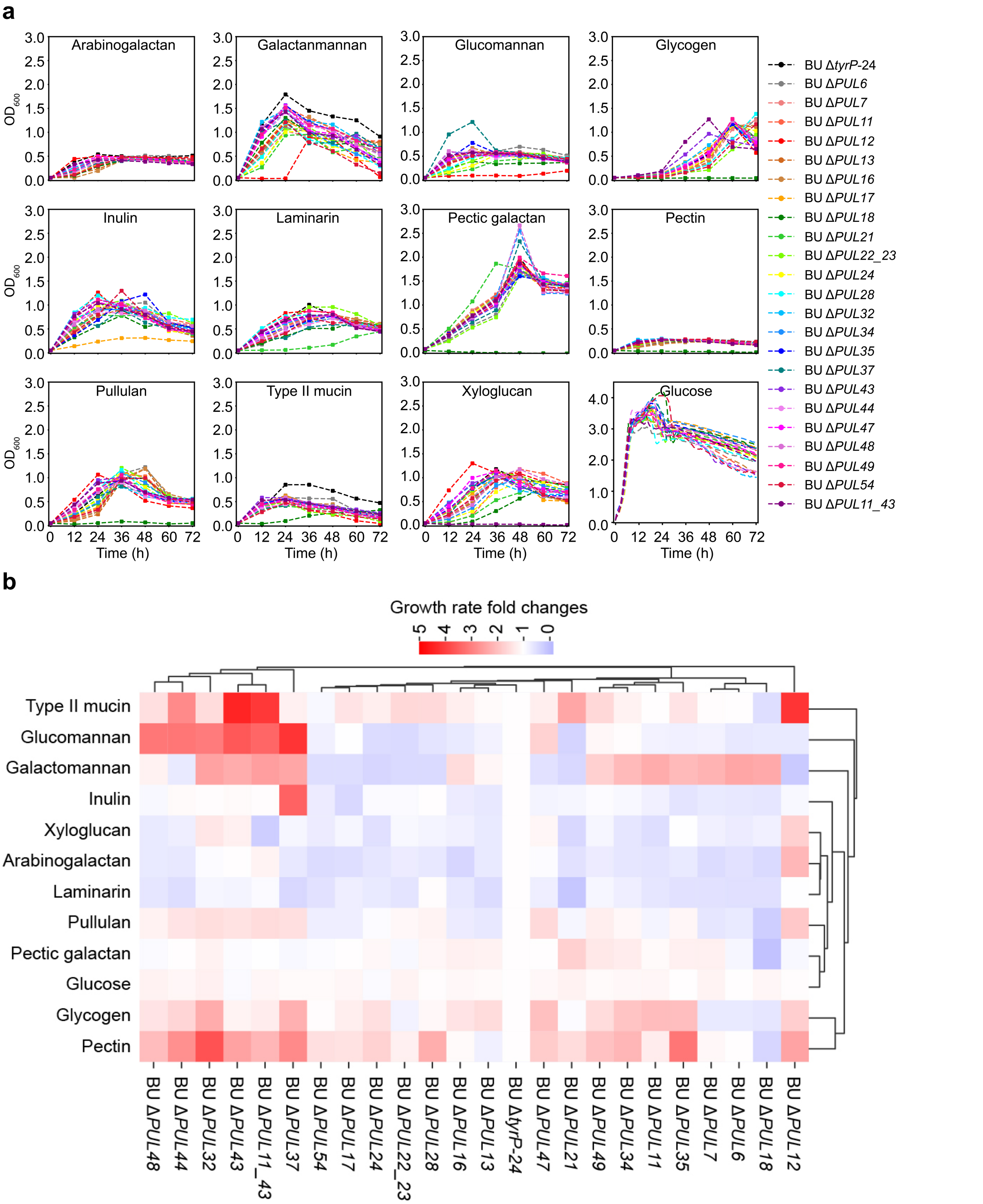

**Figure S6. Growth characterization of *B. uniformis* polysaccharide utilization loci (PULs) deletion mutants and the control strain in media supplemented with different carbon sources. (a)** Absorbance at 600 nm ($OD_{600}$) as a function of time of *B. uniformis* PUL deletion mutants and control strain Δ*tyrP*-24 in BMM-C (*Bacteroides* minimal media without carbon source) supplemented with different carbon sources. Data points represent the mean of 3 independent replicates. The inulin used in this experiment was from Acros Organics (Catalog number AC457100250). **(b)** Biclustered heatmap of the fold changes of inferred growth rates of each PUL deletion mutant compared to the control Δ*tyrP*-24. Growth rates were inferred by fitting the generalized Lotka Voterra (gLV) model to the time-series data in panel **(a)** (**Methods**, **Supplementary Data 2**).

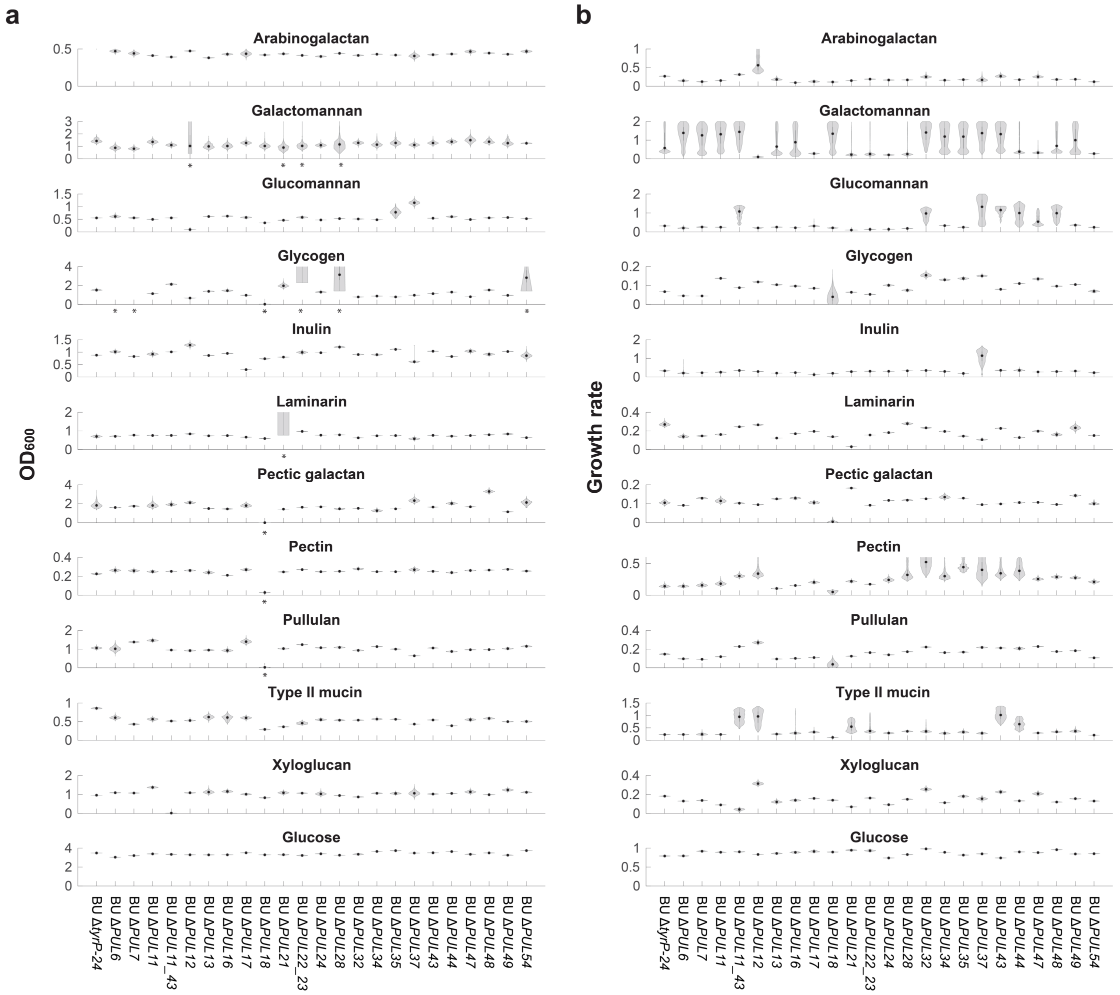

**Figure S7. Violin plots of the inferred distributions of carrying capacities and growth rates for *B. uniformis* polysaccharide utilization loci deletion mutants and the control strain in media containing different carbon sources.** Each subplot represents the violin plots for all *B. uniformis* mutants in media with a carbon source, which is indicated in the title of the subplot. Violin plots in panel **(a)** with an asterisk labeled on the x-axis represent carrying capacity distributions with coefficient of variation greater than 0.2 (**Supplementary Data 2**). The inferred carrying capacities in these conditions are replaced by the maximum $OD_{600}$ of the experiment in **Figure 2b** in the main text.

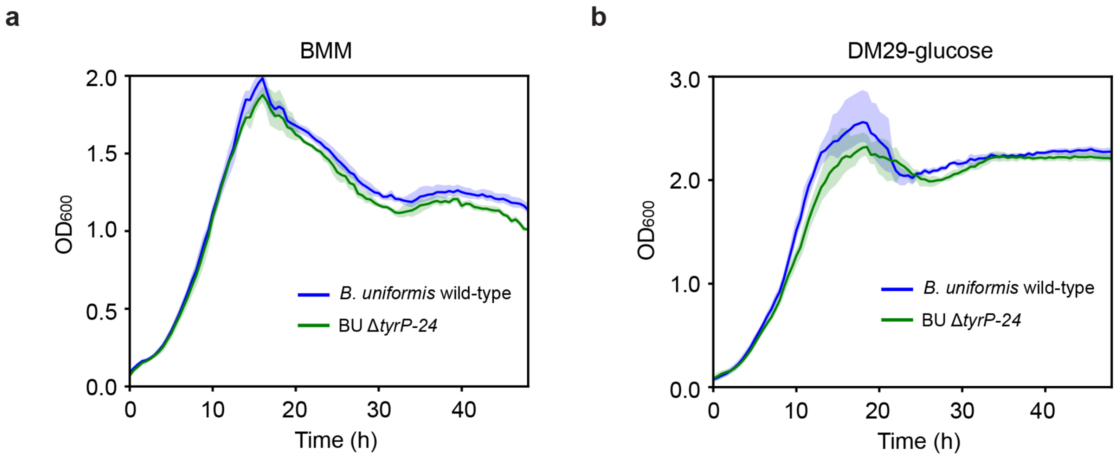

**Figure S8. Growth characterization of *B. uniformis* wild-type and the barcoded control strain Δ*tyrP-24* in (a) *Bacteroides* minimal media (BMM) or (b) DM29-glucose media (b).** Lines denote the mean and shaded regions represent the 95% confidence interval (n=3).

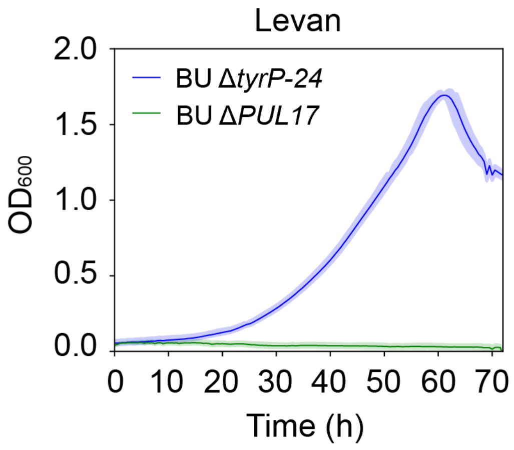

**Figure S9. Growth characterization of *B. uniformis PUL17* deletion (Δ*PUL17*) and the control strain Δ*tyrP-24*.** The mutant Δ*PUL17* and control strain Δ*tyrP-24* were grown in BMM-C (*Bacteroides* minimal media without carbon source) supplemented with levan. Lines denote the mean and the shaded regions represent 95% confidence interval (n=3).

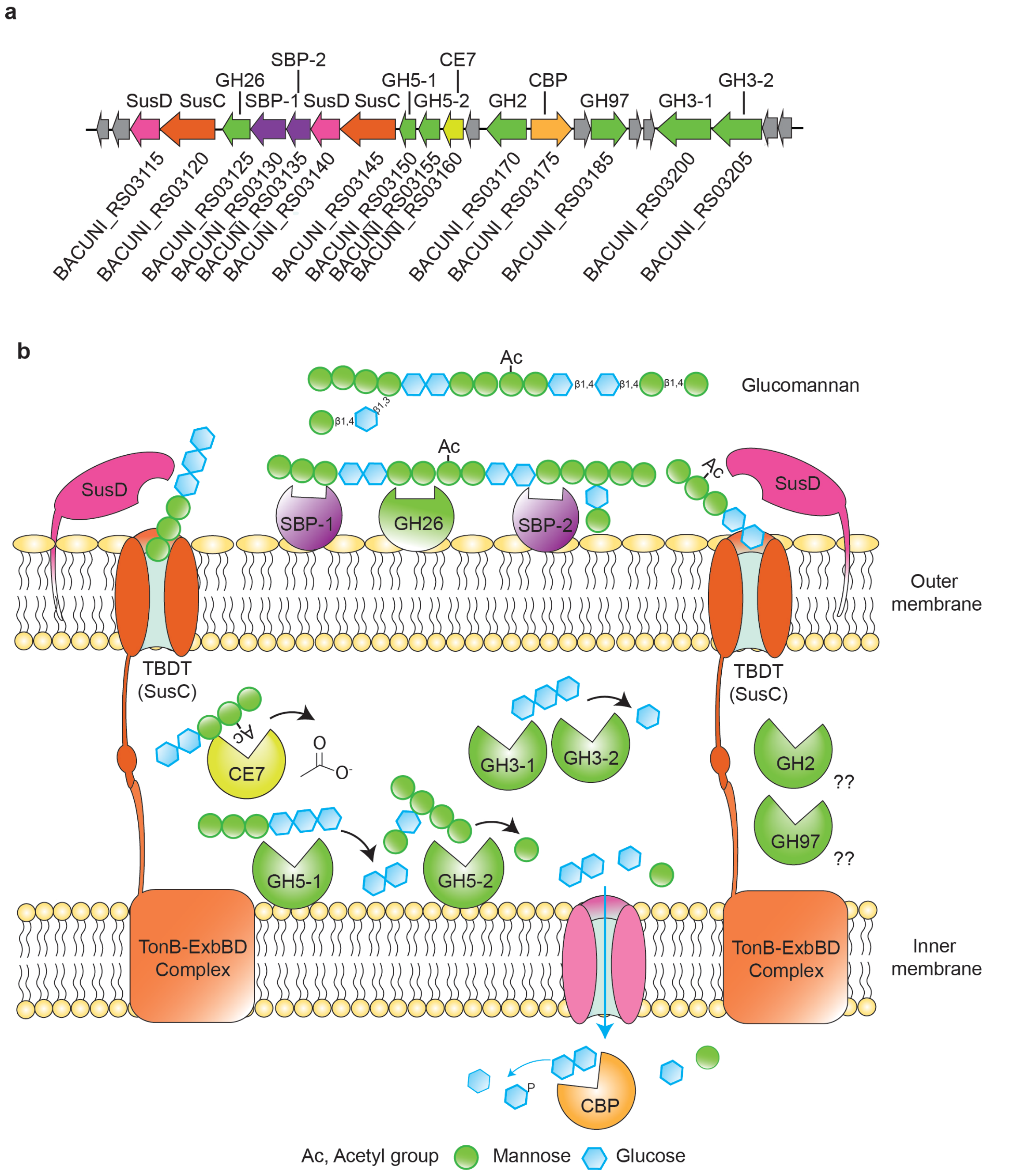

**Figure S10. Predicted biochemical model for glucomannan utilization by *PUL12* in *B. uniformis*. (a)** Schematic of gene organization of *PUL12*, colored by predicted protein function shown in subsequent panels. Genes with unknown predicted functions are represented in gray. GenBank locus tag numbers are provided below each gene. SusC: SusC-like TonB-dependent transporter (TBDT); SusD: SusD-like cell-surface glycan-binding protein; SBP, sugar binding protein; CBP: cellobiose phosphorylase. **(b)** Schematic of the predicted biochemical model of glucomannan utilization in *B. uniformis* based on sequence homology, protein structure and predicted subcellular localization of the enzymes (**Supplementary Data 1**). The detailed descriptions of the model can be found in **Supplementary note**.

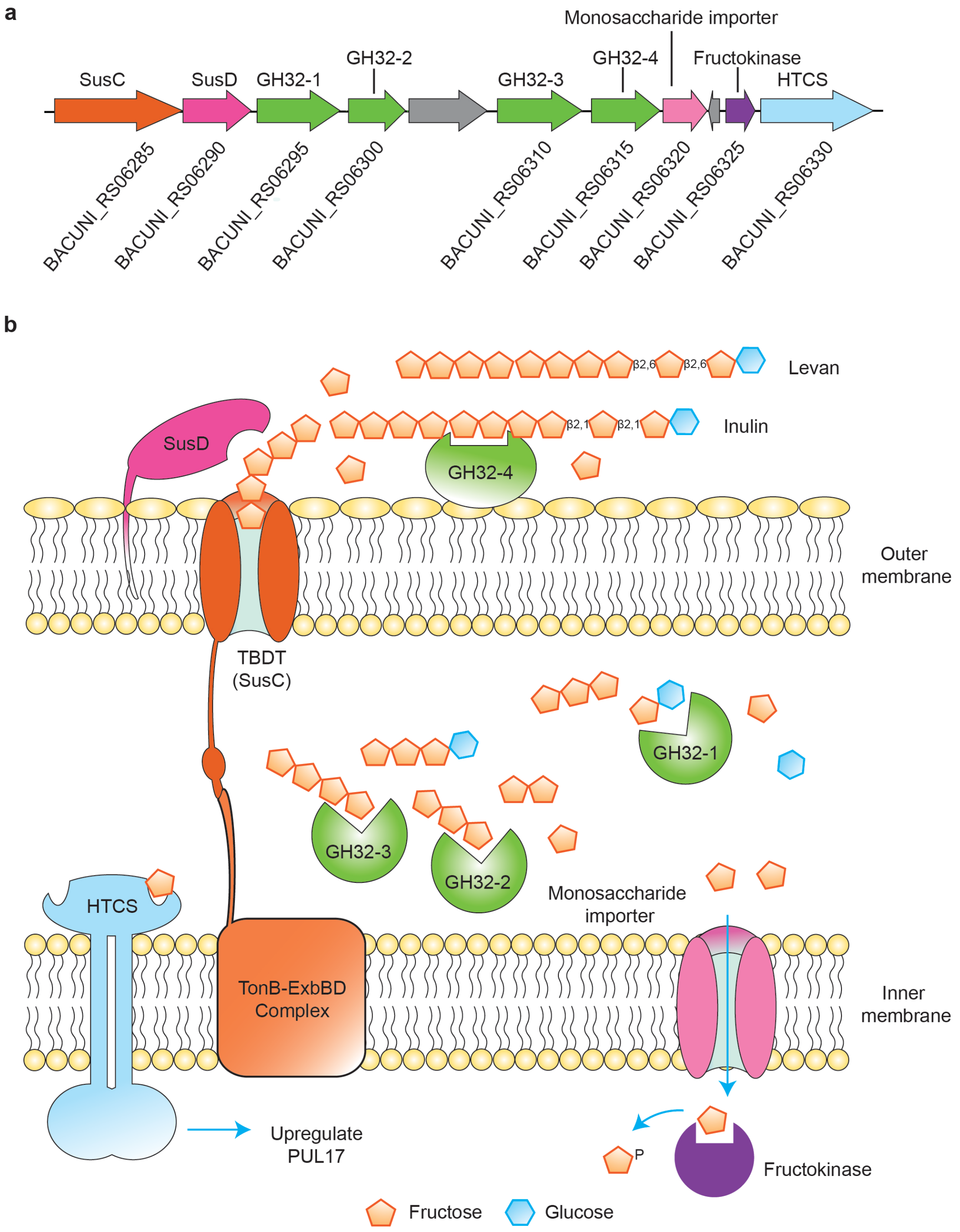

**Figure S11. Predicted biochemical model of inulin and levan utilization by *PUL17* in *B. uniformis*. (a)** Schematic of gene organization of *PUL17*, colored with reference to the proteins shown in subsequent panels. Genes with unknown functions are represented in gray. GenBank locus tag numbers are also provided below each gene. SusC: SusC-like TonB-dependent transporter (TBDT); SusD: SusD-like cell-surface glycan-binding protein; HTCS, hybrid two-component system. **(b)** Schematic of predicted biochemical model of inulin and levan utilization in *B. uniformis*. The prediction is based on a previously characterized levan utilization loci in *B. thetaiotaomicron*[22], sequence homology, protein structure and predicted subcellular localization of the enzymes (**Supplementary Data 1**). The detailed descriptions of the biochemical model can be found in **Supplementary note**.

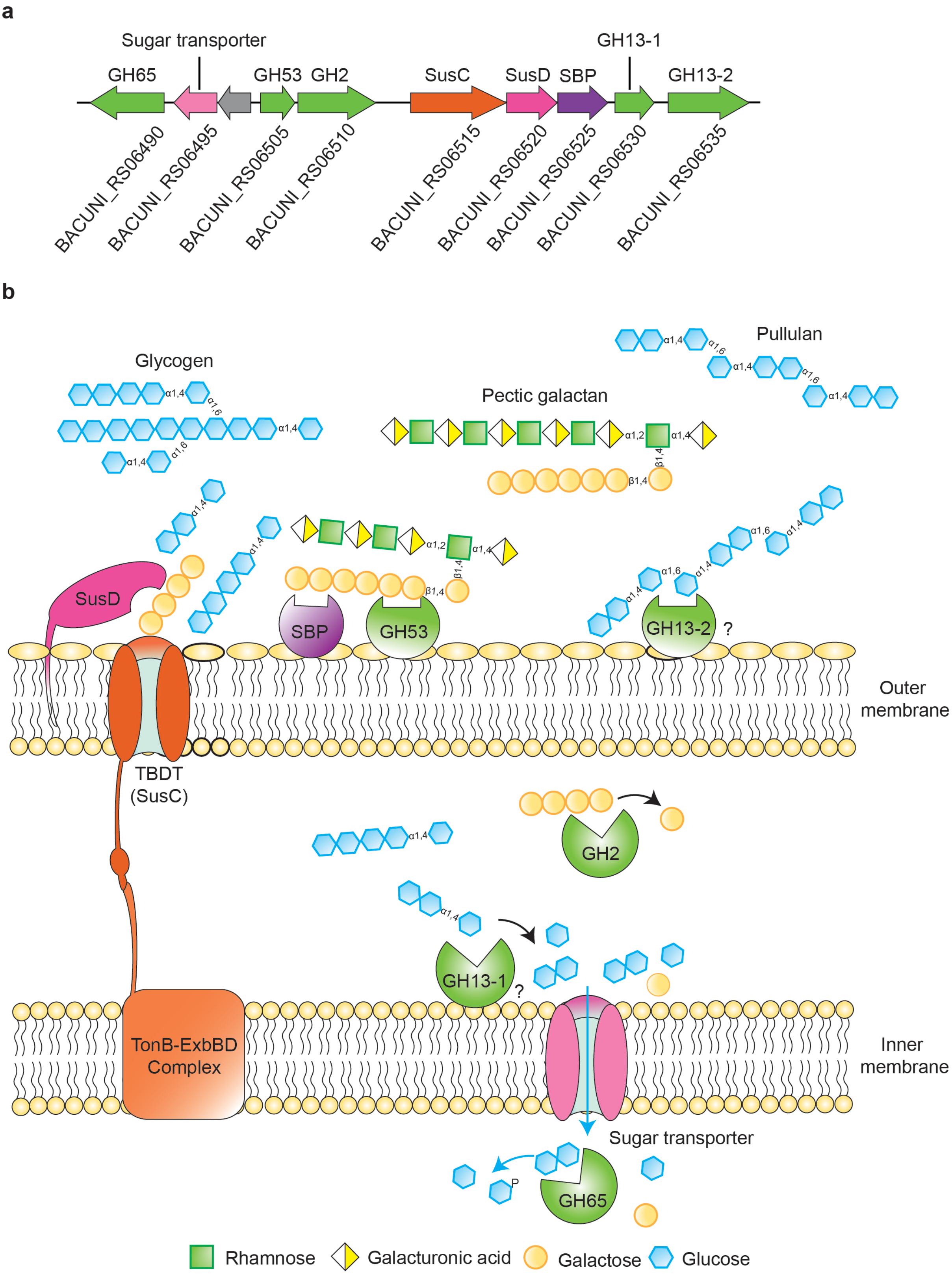

**Figure S12. Predicted biochemical model of glycogen, pullulan, pectic galactan (or pectin) utilization by *PUL18* in *B. uniformis*. (a)** Schematic of gene organization of *PUL18*, colored with reference to the proteins shown in subsequent panels. Genes with unknown functions are represented in gray. GenBank locus tag numbers are also provided below each gene. SusC: SusC-like TonB-dependent transporter (TBDT); SusD: SusD-like cell-surface glycan-binding protein; SBP: sugar binding protein. **(b)** Schematic of predicted biochemical model of glycogen, pullulan and pectic galactan utilization in *B. uniformis*. The prediction is based on sequence homology, protein structure and predicted subcellular localization of the enzymes (**Supplementary Data 1**). The detailed descriptions of the model can be found in **Supplementary note**. The symbol '?' indicates uncertainty in the prediction of subcellular localization.

a

b

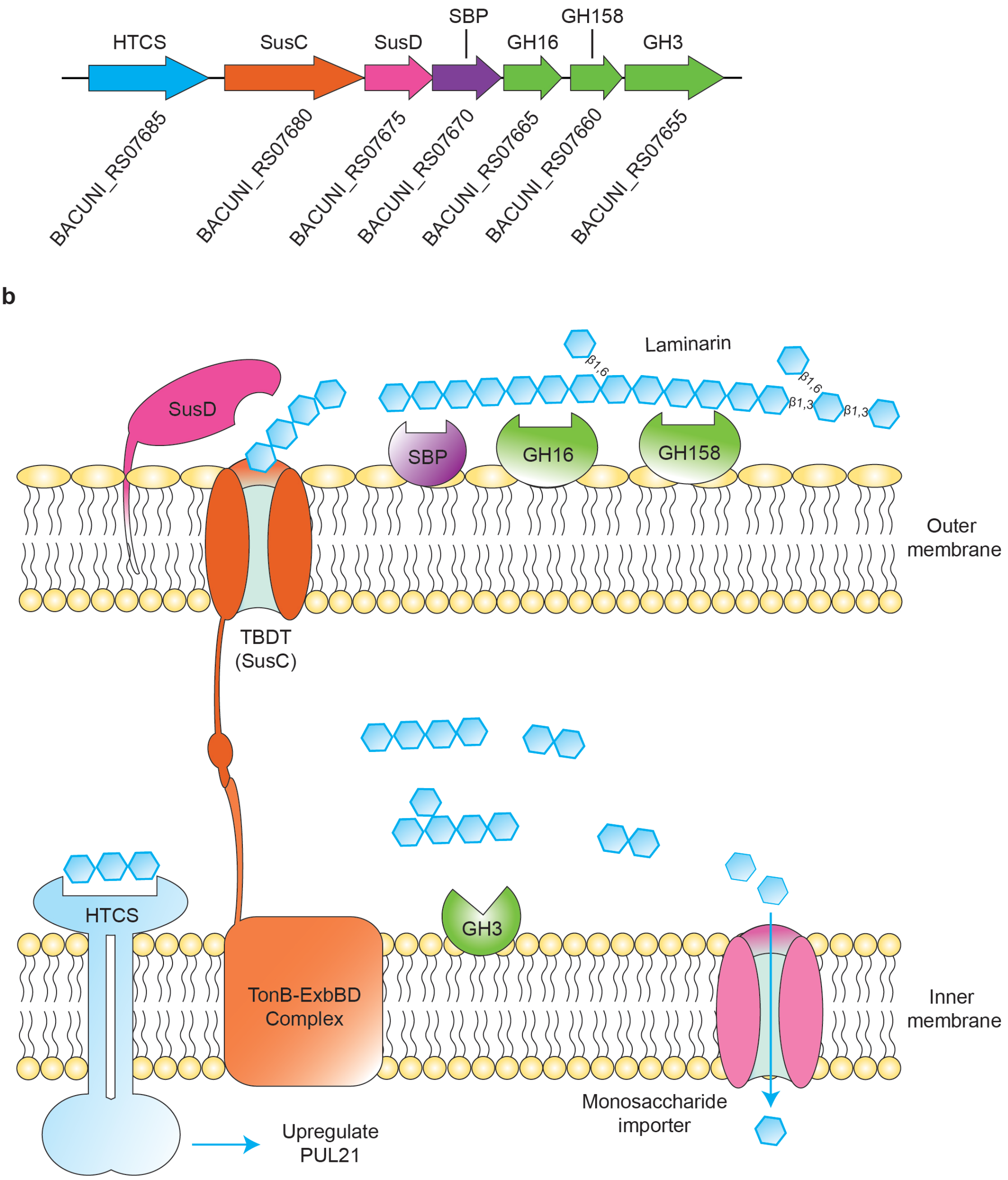

**Figure S13. Predicted biochemical model of laminarin utilization by *PUL21* in *B. uniformis*. (a)** Schematic of gene organization of *PUL21*, colored with reference to the proteins shown in subsequent panels. GenBank locus tag numbers are also provided below each gene. SusC: SusC-like TonB-dependent transporter (TBDT); SusD: SusD-like cell-surface glycan-binding protein; HTCS, hybrid two-component system. SBP: sugar binding protein. **(b)** Schematic of predicted biochemical model of laminarin utilization in *B. uniformis*[44].

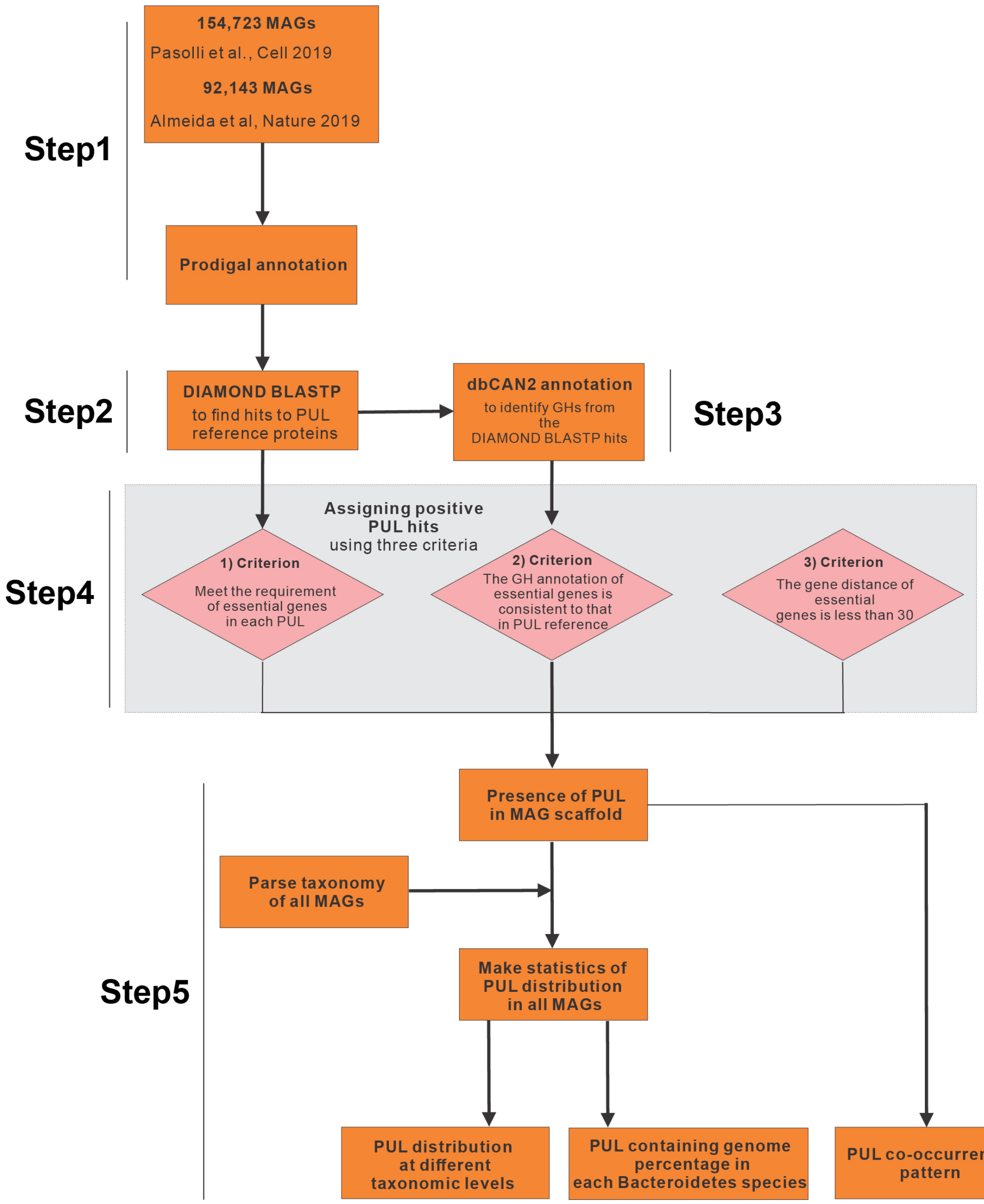

**Figure S14. Schematic of workflow for bioinformatic analysis of PULs in human gut microbiome metagenome-assembled genome datasets.** The detailed description of the workflow can be found in the **Methods**.

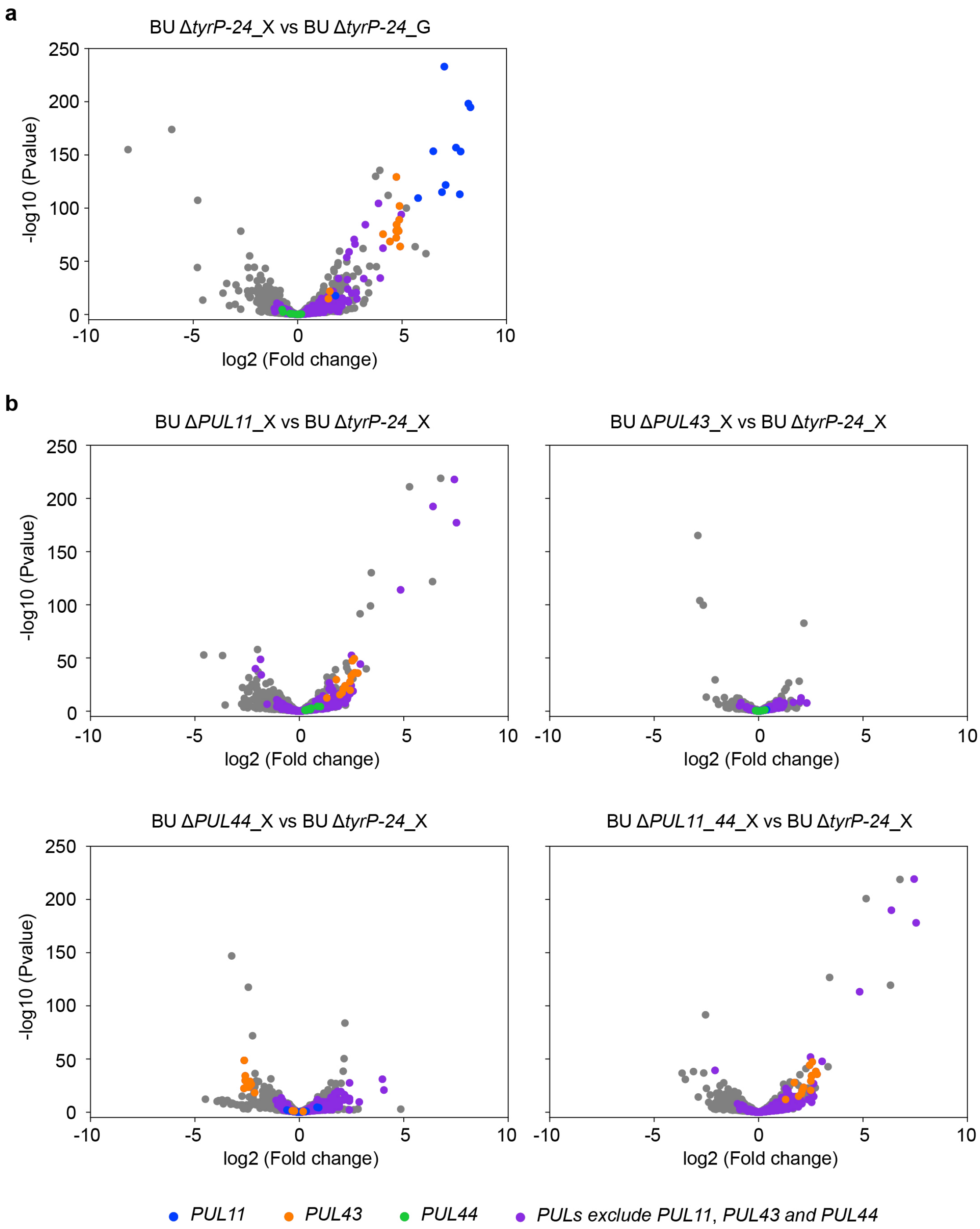

**Figure S15. Transcriptional fold changes of *B. uniformis* PUL mutants and the control strain Δ*tyrP-24*. (a)** Scatter plot of log2 fold changes of transcript abundance versus -log10(p-value) for Δ*tyrP-24* in xyloglucan media (X) compared to glucose media (G). In all sub-plots, gray, blue, orange, green, purple data points denote genes not associated with PULs, *PUL11* genes, *PUL43* genes, *PUL44* genes, or genes found in all PULs excluding *PUL11*, *PUL43* and *PUL44*, respectively. **(b)** Scatter plot of log2 fold change of transcript abundance versus -log10(p-value) for *B. uniformis* mutants Δ*PUL11*, Δ*PUL43*, Δ*PUL44* or Δ*PUL11_44* compared to Δ*tyrP-24* in xyloglucan media. Each RNA-seq condition consists of 2 biological replicates (**Supplementary Data 6**).

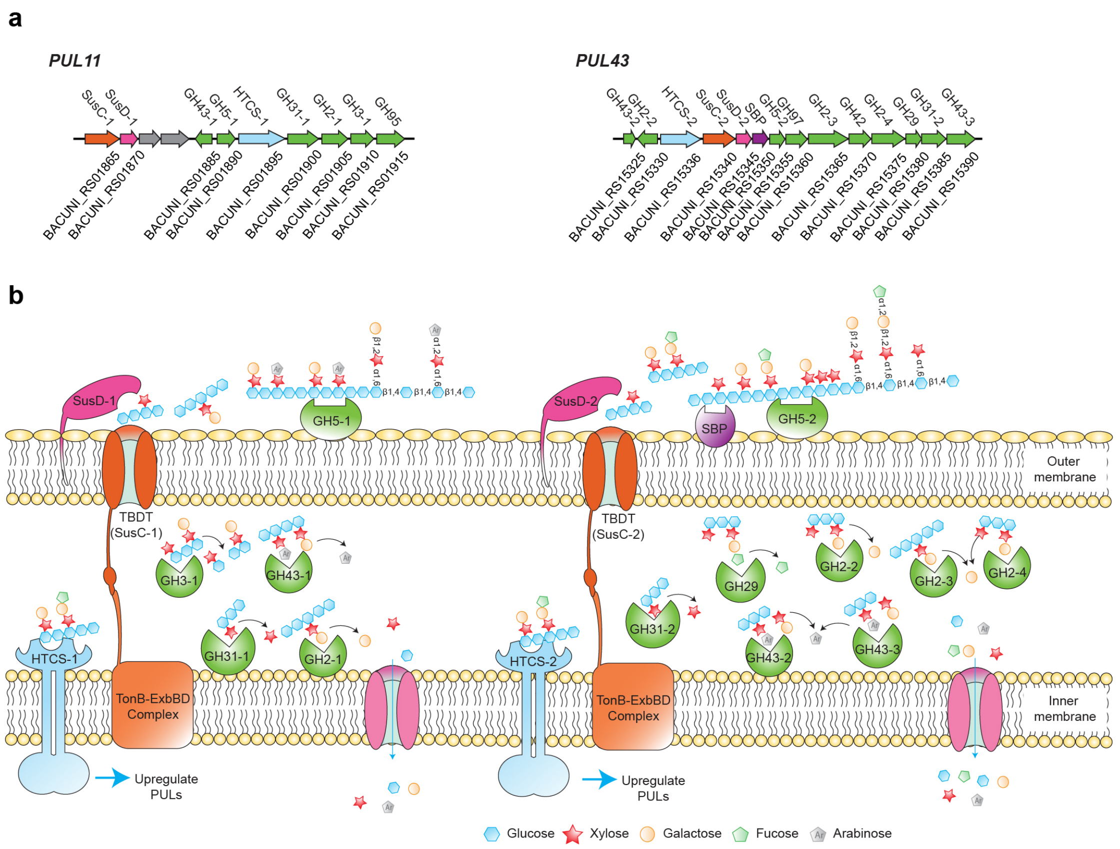

**Figure S16. Predicted biochemical model of xyloglucan utilization by *PUL11* and *PUL43* in *B. uniformis*. (a)** Schematic of gene organization of *PUL11* and *PUL43*, colored with reference to the proteins shown in subsequent panels. Genes with unknown functions are represented in gray. GenBank locus tag numbers are also provided below each gene. SusC: SusC-like TonB-dependent transporter (TBDT); SusD: SusD-like cell-surface glycan-binding protein; HTCS, hybrid two-component system; SBP: sugar binding protein. **(b)** Schematic of predicted biochemical model of xyloglucan utilization in *B. uniformis*. The prediction is based on a previous studied xyloglucan utilization loci in *B. ovatus*[20,34], sequence homology, structure similarities and subcellular localization of the enzymes (**Supplementary Data 1**). The detailed descriptions of the model can be found in **Supplementary note**.

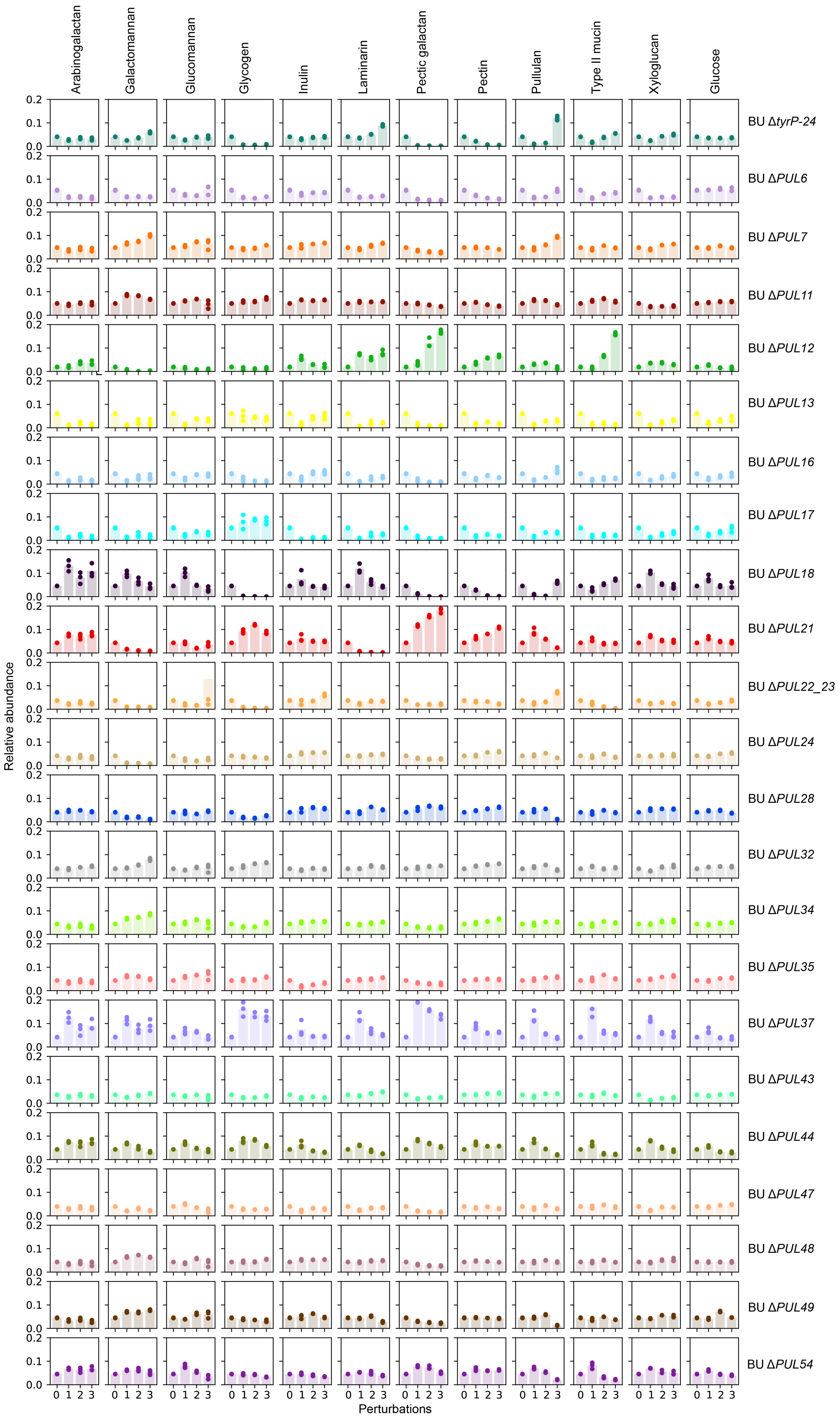
Arabinogalactan
Galactomannan
Glucomannan
Glycogen
Inulin
Laminarin
Pectic galactan
Pectin
Pullulan
Type II mucin
Xyloglucan
Glucose
BU ΔtyrP-24
BU ΔPUL6
BU ΔPUL7
BU ΔPUL11
BU ΔPUL12
BU ΔPUL13
BU ΔPUL16
BU ΔPUL17
BU ΔPUL18
BU ΔPUL21
BU ΔPUL22_23
BU ΔPUL24
BU ΔPUL28
BU ΔPUL32
BU ΔPUL34
BU ΔPUL35
BU ΔPUL37
BU ΔPUL43
BU ΔPUL44
BU ΔPUL47
BU ΔPUL48
BU ΔPUL49
BU ΔPUL54
Relative abundance
Perturbations

**Figure S17. Relative abundance of *B. uniformis* mutants in the mutant pool for different dilution perturbation conditions.** Bar plots show the mean value of relative abundance of each PUL deletion mutant or control Δ*tyrP-24* strain in different dilution perturbation conditions described in **Fig. 4a** across a range of media supplemented with different carbon sources. Data points indicate the relative abundance of 3 biological replicates.

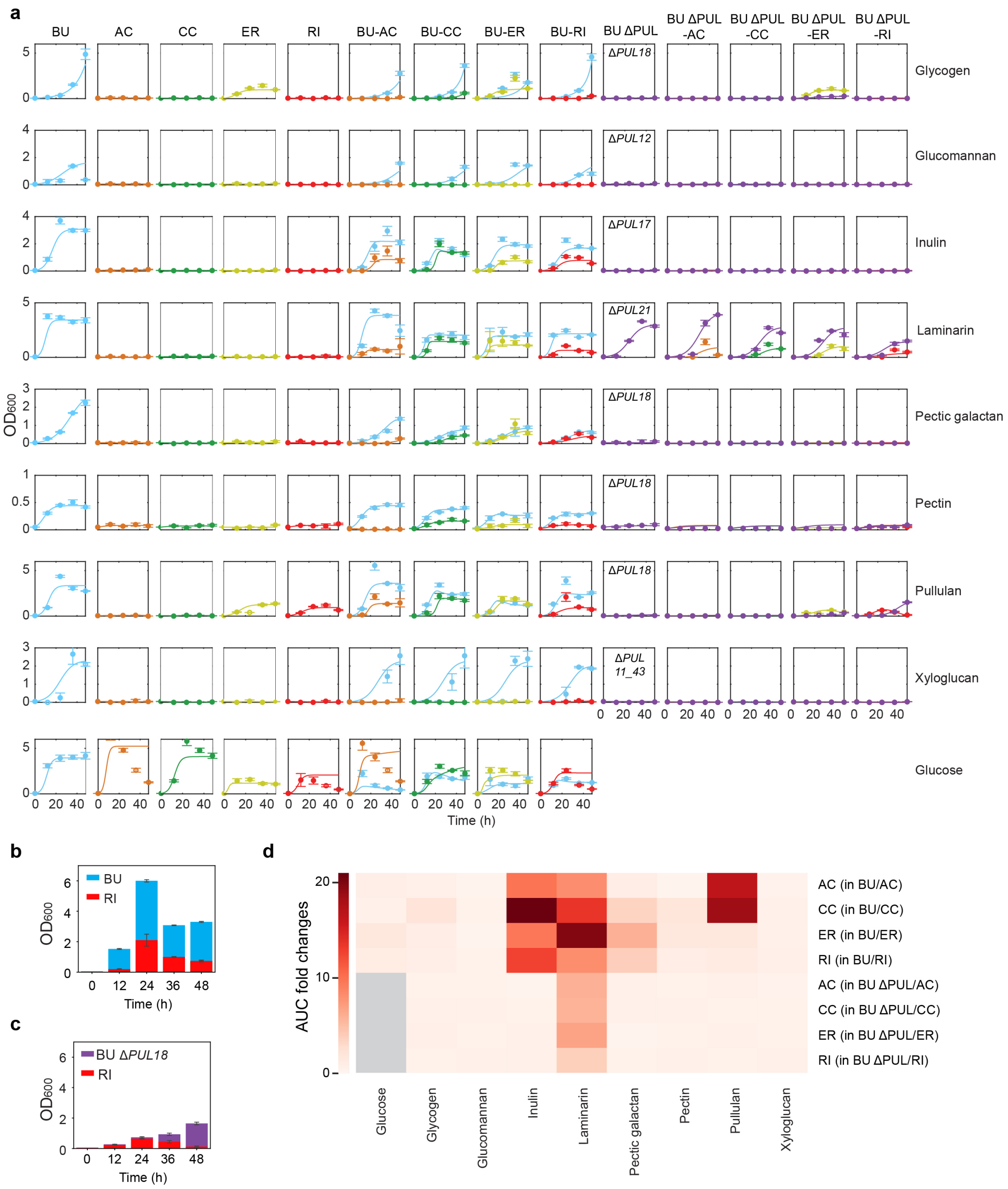

**Figure S18. Investigating inter-species interactions between *B. uniformis* wild-type, PUL deletion mutants and butyrate producers. (a)** Time-series measurements of species absolute abundance determined by 16S rDNA sequencing multiplied by $OD_{600}$ of *B. uniformis* wild-type, PUL deletion mutants and butyrate producers including *A. caccae* (AC), *C. comes* (CC), *E. rectale* (ER) and *R. intestinalis* (RI) in monoculture and co-culture conditions across media supplemented with different carbon sources. **(b)** Stacked bar plot of the absolute abundance of *B. uniformis* wild-type and *R. intestinalis* in co-culture in pullulan media for easier visualization (same data as shown in panel **(a)**). The height of the bars denotes the mean and error bars are +/- 1 s.d. (n = 3). **(c)** Stacked bar plot of the absolute abundance of Δ*PUL18* and *R. intestinalis* in co-culture in pullulan media for easier visualization (same data as shown in panel **(a)**). The height of the bars denotes the mean and error bars are +/- 1 s.d. (n = 3). **(d)** Heatmap of the total growth (area under the curve or AUC) fold changes of butyrate producers in co-culture with *B. uniformis* compared to the AUC of each butyrate producer in monoculture. For AUC calculation, when the measured $OD_{600}$ value was lower than the initial $OD_{600}$, we set the $OD_{600}$ to the value of the initial $OD_{600}$. The gray color represents no experiment performed in those conditions.

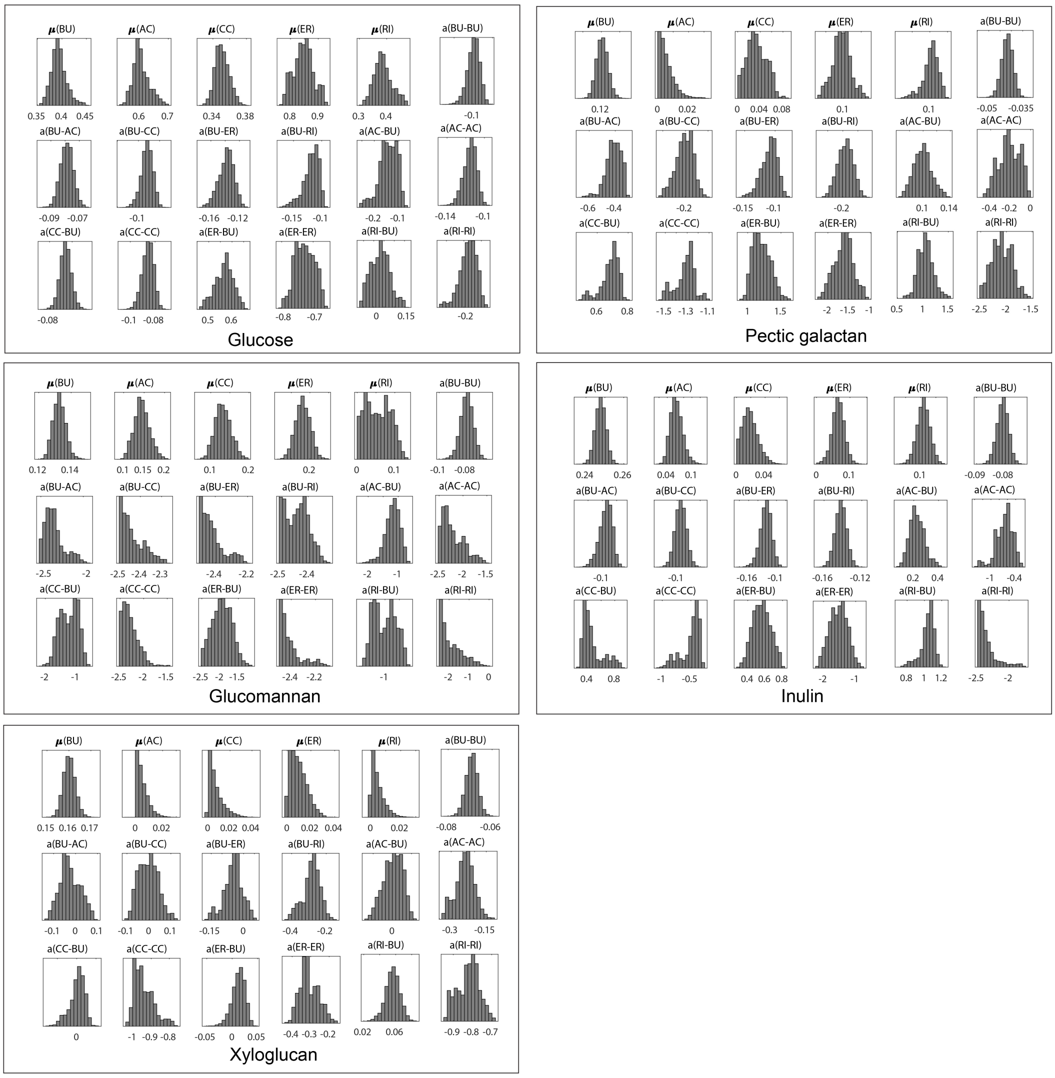

**Figure S19. Marginal distributions of growth rates (μ) and inter-species interaction coefficients inferred from the experiments described in Fig. 6a using Markov chain Monte Carlo in media with different carbon sources.** Each parameter $a_{ij}$ represents the inter-species interaction coefficient of species $j$ on the growth of species $i$, where $i$ and $j$ represent *B. uniformis* (BU), *A. caccae* (AC), *C. comes* (CC), *E. rectale* (ER) or *R. intestinalis* (RI).

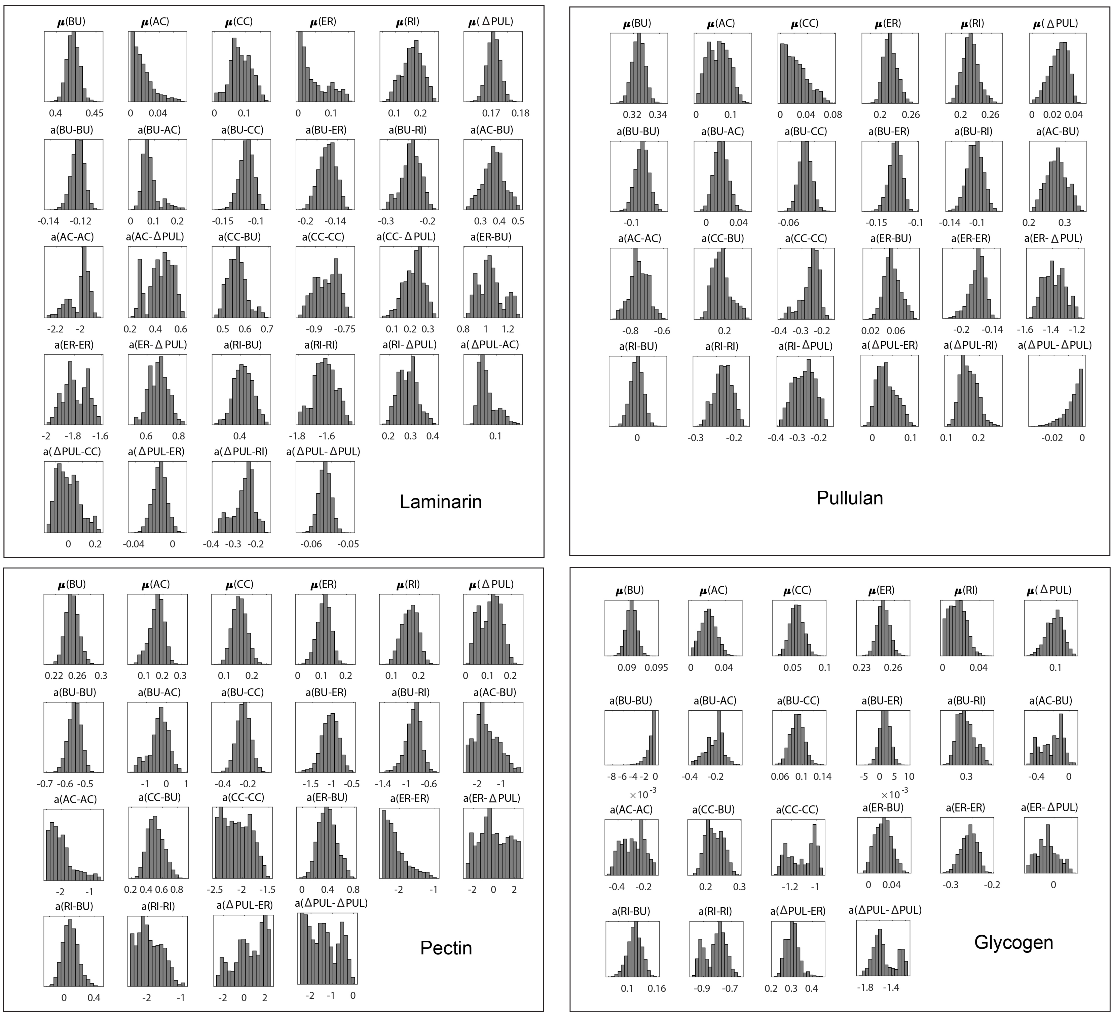

**Figure S20. Marginal distributions of inferred growth rates (μ) and inter-species interaction coefficients based on experiments described in Fig. 6a using Markov chain Monte Carlo in media with different carbon sources.** The *B. uniformis* mutants ΔPUL represents Δ*PUL21* in laminarin and Δ*PUL18* in pullulan, pectin and glycogen. Each parameter $a_{ij}$ denotes the inter-species interaction coefficient of species $j$ on the growth of species $i$, where $i$ and $j$ represent *B. uniformis* (BU), *A. caccae* (AC), *C. comes* (CC), *E. rectale* (ER), *R. intestinalis* (RI) or a given PUL deletion mutant.

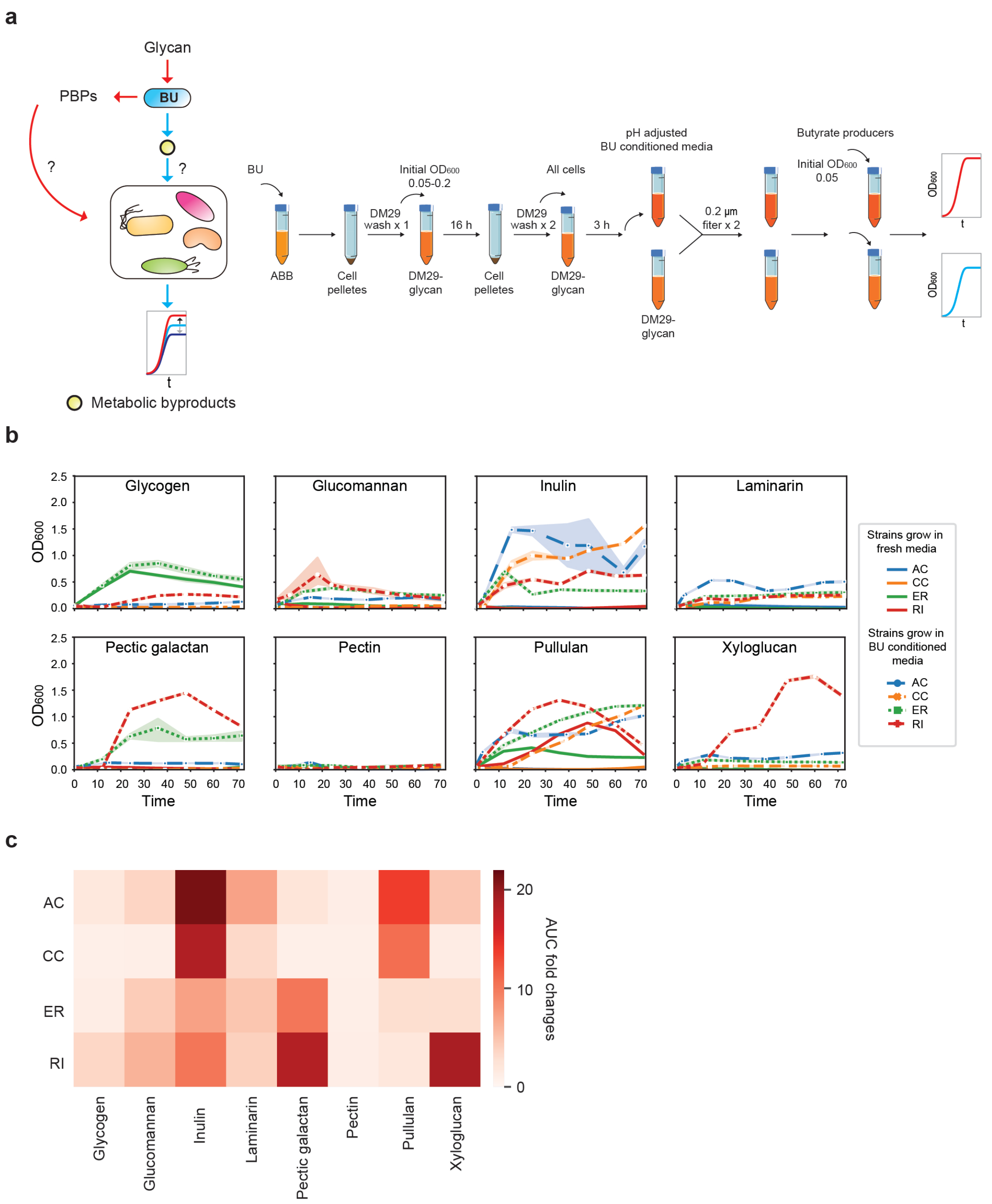

**Figure S21. Effects of *B. uniformis* conditioned glycan media on the growth of butyrate producers. (a)** Schematic of experimental design. BU represents *B. uniformis* wild-type strain and PBPs denotes the polysaccharide breakdown products. **(b)** Time-series measurements of absorbance at 600 nm ($OD_{600}$) for the butyrate producers *A. caccae* (AC), *C. comes* (CC), *E. rectale* (ER) and *R. intestinalis* (RI) in *B. uniformis* conditioned glycan media or fresh glycan media. Lines denote the mean and shaded regions represent 95% confidence interval (n=2-3). **(c)** Heatmap of the total growth (area under the curve or AUC) fold changes of each butyrate producer in *B. uniformis* conditioned glycan media compared to the AUC of each butyrate producer in fresh glycan media. In cases where the measured $OD_{600}$ was lower than the initial $OD_{600}$, we set the $OD_{600}$ to the initial $OD_{600}$.

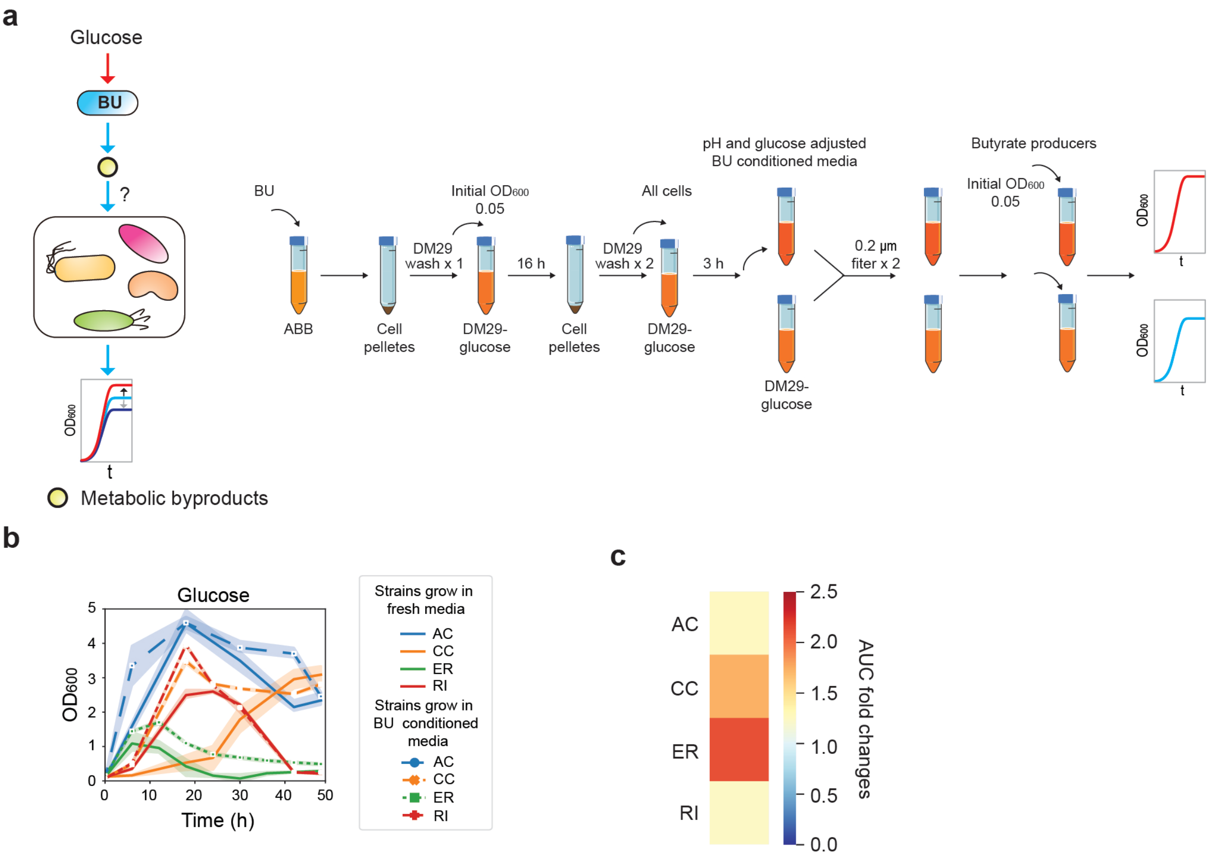

**Figure S22. Effects of *B. uniformis* metabolic byproducts on the growth of butyrate producers. (a)** Schematic of the experimental design. BU represents *B. uniformis* wild-type strain. **(b)** Time-series measurements of absorbance at 600 nm ($OD_{600}$) of the butyrate producers including *A. caccae* (AC), *C. comes* (CC), *E. rectale* (ER) and *R. intestinalis* (RI) in pH and glucose adjusted *B. uniformis* (BU) conditioned glucose media or fresh glucose media. Lines denote the mean and shaded regions represent 95% confidence interval (n=2-3). **(c)** Heatmap of the total growth (area under the curve or AUC) fold changes of each butyrate producer in *B. uniformis* conditioned glucose media compared to the AUC of each butyrate producer in fresh glucose media. In cases where the measured $OD_{600}$ value was lower than the initial $OD_{600}$, we set the $OD_{600}$ to the initial $OD_{600}$.

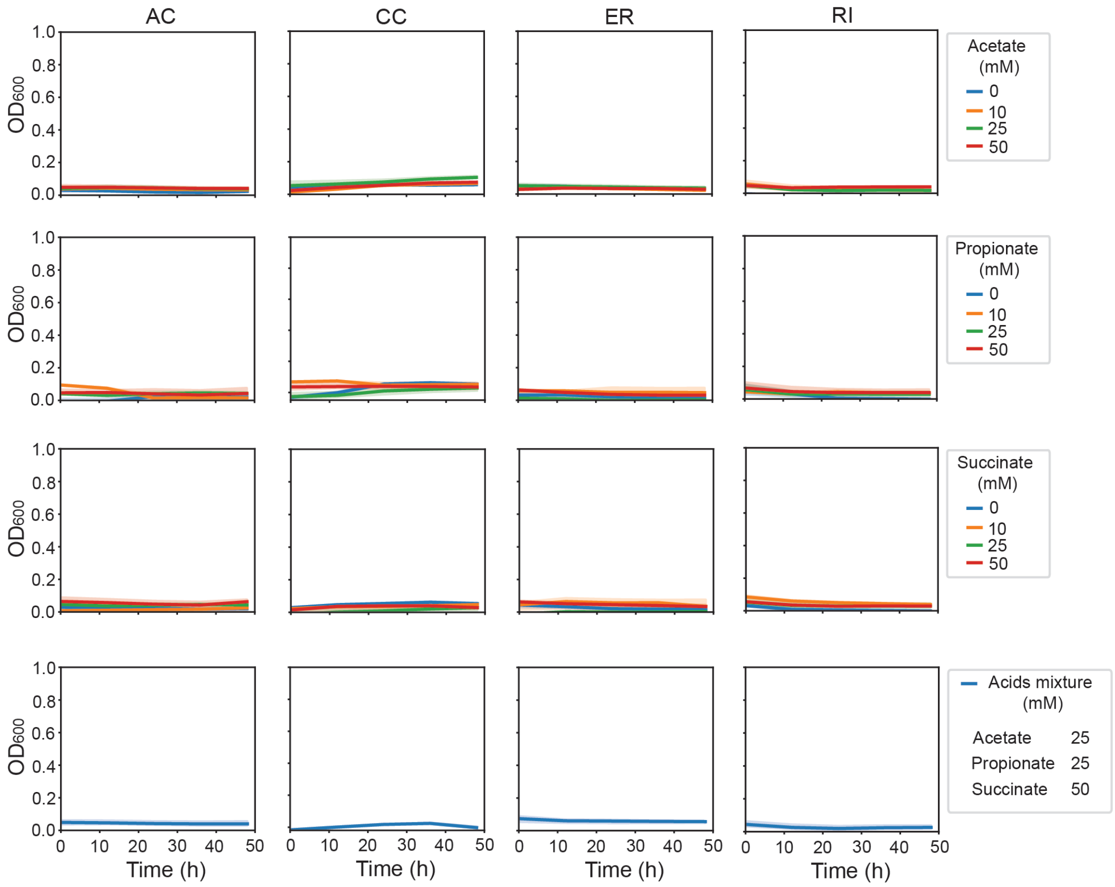

**Figure S23. Growth responses of butyrate producers in media with fermentation end products as the primary carbon source.** Time-series measurements of absorbance at 600 nm ($OD_{600}$). The butyrate producers including *A. caccae* (AC), *C. comes* (CC), *E. rectale* (ER) and *R. intestinalis* (RI) were grown in DM29 media with addition of 0, 10, 25 or 50 mM acetate or propionate or succinate or the mixture of these fermentation end products (25 mM of acetate, 25 mM of propionate and 50 mM of succinate). Lines denote the mean and shaded regions represent 95% confidence interval (n=3).

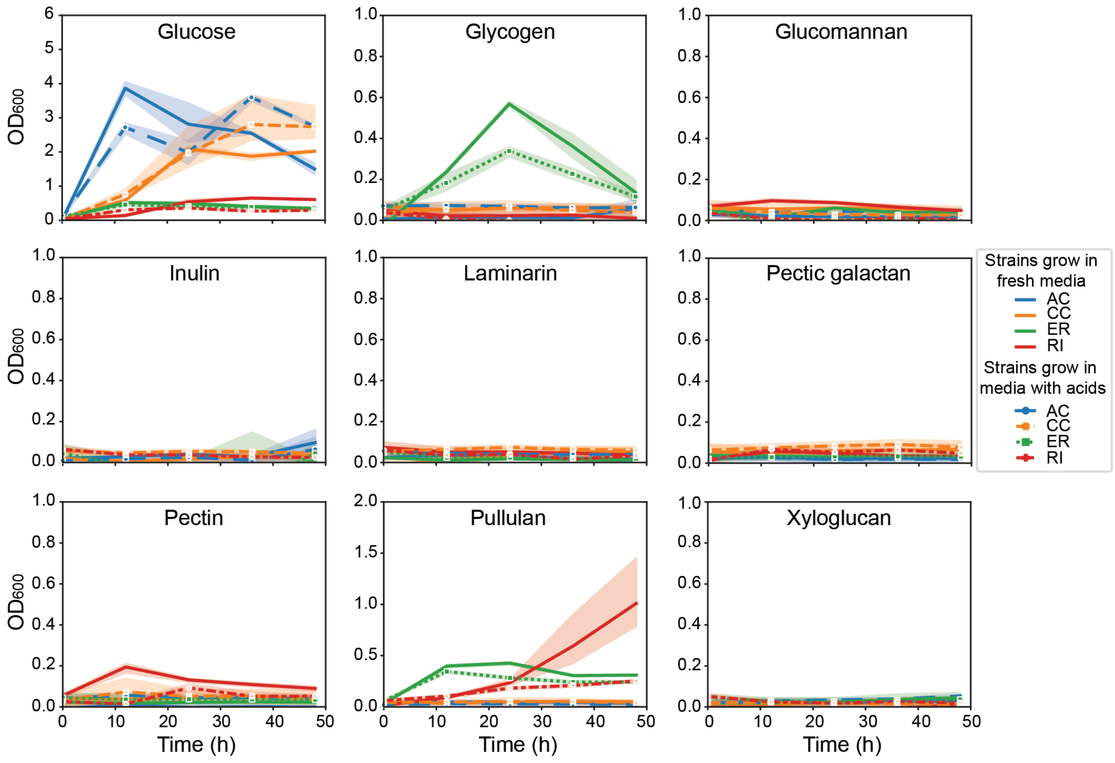

**Figure S24. Growth response of butyrate producers in DM29-glucose or DM29-glycan media supplemented with fermentation end products.** Time-series measurements of absorbance at 600 nm ($OD_{600}$). The butyrate producers *A. caccae* (AC), *C. comes* (CC), *E. rectale* (ER) or *R. intestinalis* (RI) were grown in DM29-glucose or DM29-glycan media with or without the addition of a mixture of 25 mM of acetate, 25 mM of propionate and 50 mM of succinate. Lines denote the mean and shaded regions represent 95% confidence interval (n=3).

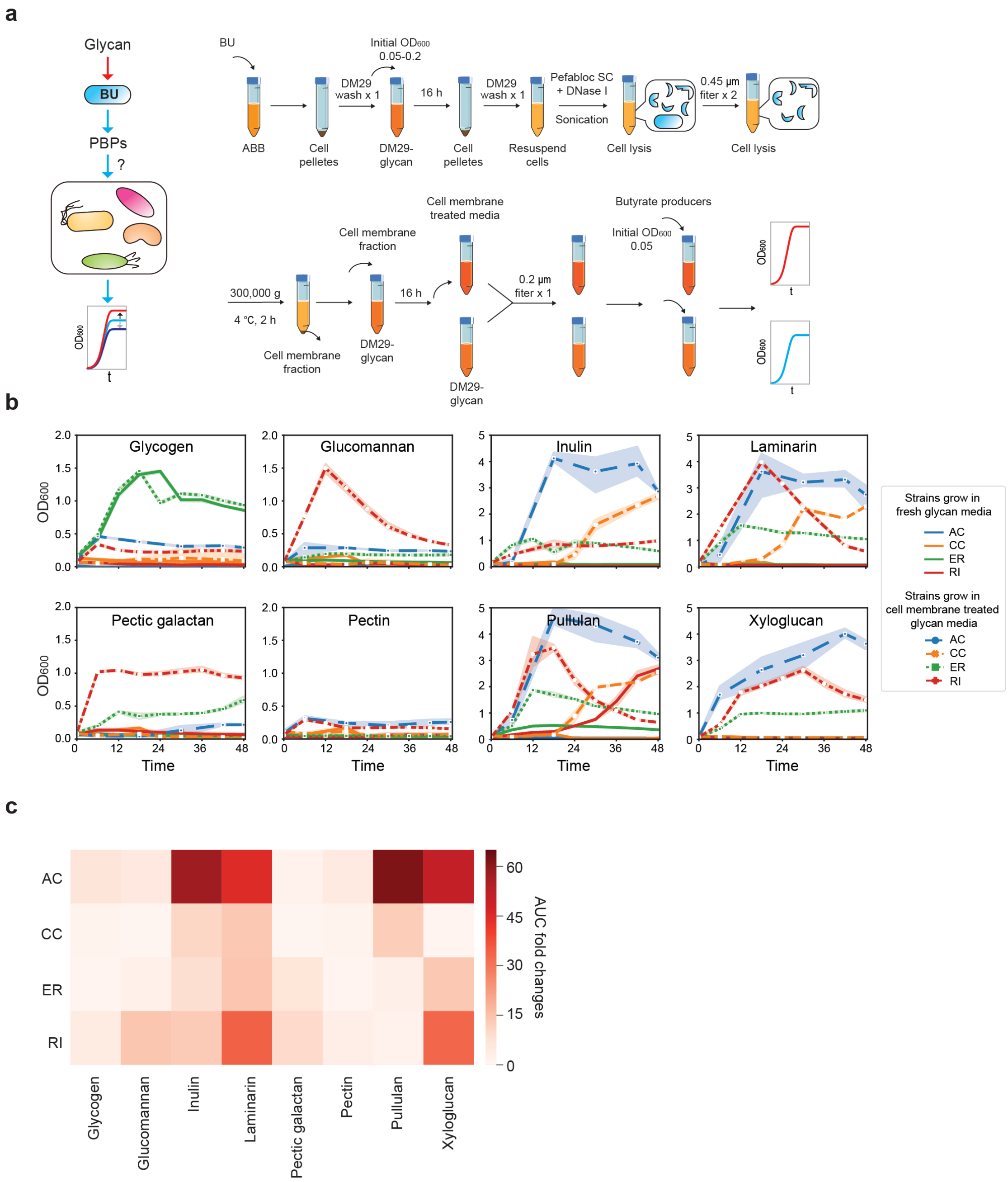

**Figure S25. Effects of *B. uniformis* (BU) cell membrane treated glycan media on the growth of butyrate producers. (a)** Schematic of the experimental design. PBPs denote polysaccharide breakdown products. **(b)** Time-series measurements of absorbance at 600 nm ($OD_{600}$). Butyrate producers including *A. caccae* (AC), *C. comes* (CC), *E. rectale* (ER) and *R. intestinalis* (RI) were grown in BU cell membrane treated glycan media or fresh glycan media. Lines denote the mean and shaded regions represent 95% confidence interval (n=3). **(c)** Heatmap of the total growth (area under the curve or AUC) fold changes of butyrate producers in BU cell membrane treated glycan media compared to the AUC of each butyrate producer in fresh glycan media. In cases where the measured $OD_{600}$ value was lower than the initial $OD_{600}$, we set the $OD_{600}$ value to equal to the initial OD.

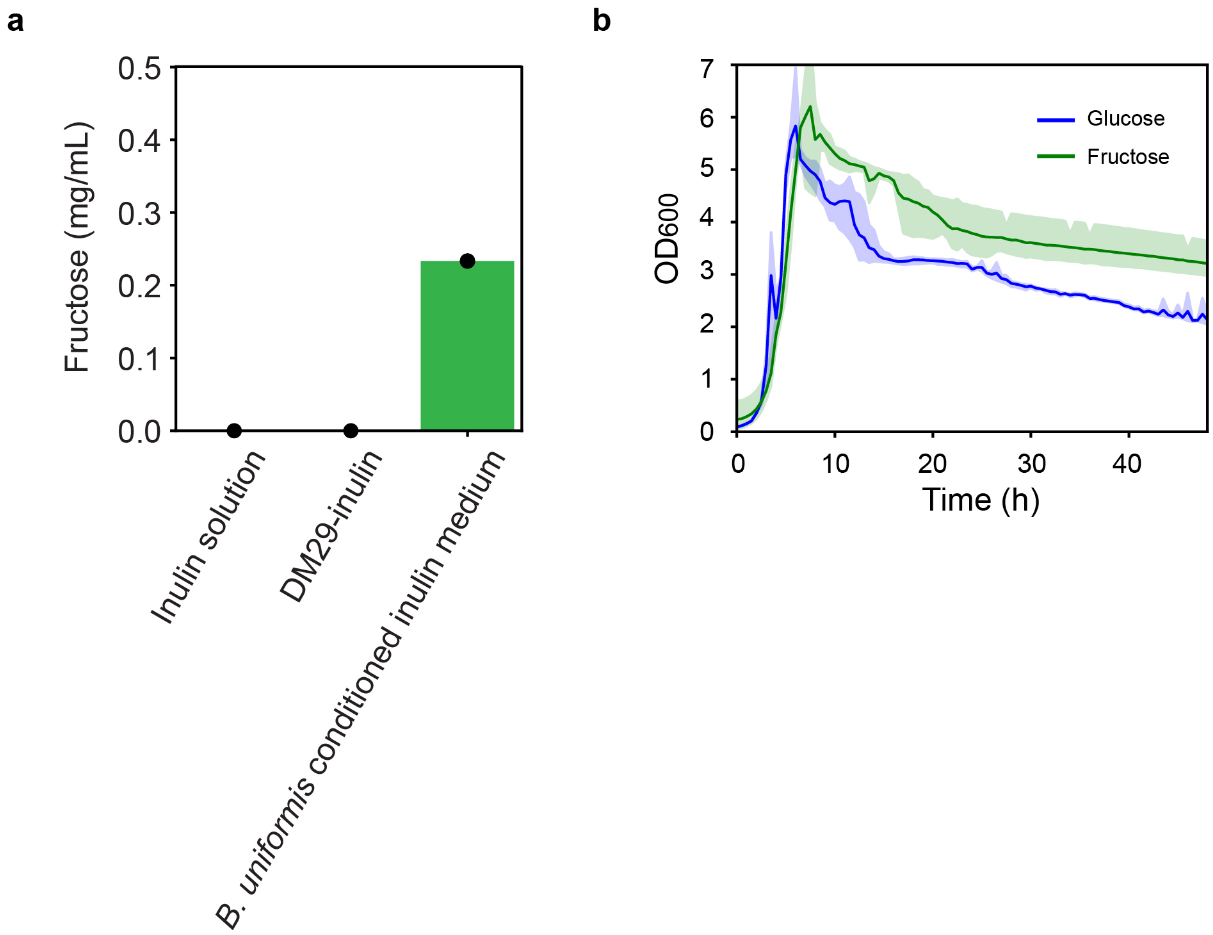

**Figure S26. *A. caccae* can utilize inulin breakdown products generated by *B. uniformis* (BU). (a)** Bar plot of fructose concentrations in inulin solution, DM29-inulin or BU conditioned inulin media. The height of the bar indicates the mean and data points represent individual biological replicates. Inulin solution contained 5 g $L^{-1}$ of inulin in dd$H_2O$ and DM29-inulin media contained 5 g $L^{-1}$ of inulin. The BU conditioned inulin media was prepared as described in **Fig. S21**. (n=1 for inulin solution and DM29-inulin media; n=3 for BU conditioned inulin media) **(b)** Time-series measurements of absorbance at 600 nm ($OD_{600}$) of *A. caccae* in DM29-glucose (5 g $L^{-1}$) or DM29-fructose (5 g $L^{-1}$) media. Lines denote the mean and shaded regions represent 95% confidence interval (n=3).

**References**

1 Bayley, D. P., Rocha, E. R. & Smith, C. J. Analysis of *cepA* and other *Bacteroides fragilis* genes reveals a unique promoter structure. *FEMS Microbiology Letters* **193**, 149-154 (2000).

2 Vingadassalom, D. *et al.* An unusual primary sigma factor in the *Bacteroidetes* phylum. *Molecular Microbiology* **56**, 888-902 (2005).

3 Mimee, M., Tucker, A. C., Voigt, C. A. & Lu, T. K. Programming a human commensal bacterium, *Bacteroides thetaiotaomicron*, to sense and respond to stimuli in the murine gut microbiota. *Cell systems* **1**, 62-71 (2015).

4 Smith, C. J., Rogers, M. B. & Mckee, M. L. Heterologous gene expression in *Bacteroides fragilis*. *Plasmid* **27**, 141-154 (1992).

5 Whitaker, W. R., Shepherd, E. S. & Sonnenburg, J. L. Tunable expression tools enable single-cell strain distinction in the gut microbiome. *Cell* **169**, 538-546 (2017).

6 Parker, A. C. & Smith, C. J. Genetic and biochemical analysis of a novel Ambler class A beta-lactamase responsible for cefoxitin resistance in *Bacteroides* species. *Antimicrobial Agents and Chemotherapy* **37**, 1028-1036.

7 Lim, B., Zimmermann, M., Barry, N. A. & Goodman, A. L. Engineered regulatory systems modulate gene expression of human commensals in the gut. *Cell* **169**, 547-558 (2017).

8 Tagawa, J. *et al.* Development of a novel plasmid vector pTIO-1 adapted for electrotransformation of *Porphyromonas gingivalis*. *Journal of Microbiological Methods* **105**, 174-179 (2014).

9 Almagro Armenteros, J. J. *et al.* SignalP 5.0 improves signal peptide predictions using deep neural networks. *Nature Biotechnology* **37**, 420-423 (2019).

10 Yu, C. S. *et al.* CELLO2GO: a web server for protein subCELlular LOcalization prediction with functional gene ontology annotation. *PloS one* **9**, e99368 (2014).

11 Yu, N. Y. *et al.* PSORTb 3.0: improved protein subcellular localization prediction with refined localization subcategories and predictive capabilities for all prokaryotes. *Bioinformatics* **26**, 1608-1615 (2010).

12 Bhasin, M., Garg, A. & Raghava, G. P. PSLpred: prediction of subcellular localization of bacterial proteins. *Bioinformatics* **21**, 2522-2524 (2005).

13 Terrapon, N. *et al.* PULDB: the expanded database of polysaccharide utilization Loci. *Nucleic Acids Research* **46**, D677-D683 (2017).

14 Kelley, L. A., Mezulis, S., Yates, C. M., Wass, M. N. & Sternberg, M. J. The Phyre2 web portal for protein modeling, prediction and analysis. *Nature Protocols* **10**, 845-858 (2015).

15 Bagenholm, V. *et al.* A surface-exposed GH26 beta-mannanase from *Bacteroides ovatus*: structure, role, and phylogenetic analysis of BoMan26B. *Journal of Biological Chemistry* **294**, 9100-9117 (2019).

16 Zhu, Y. *et al.* Periplasmic *Cytophaga hutchinsonii* endoglucanases are required for use of crystalline cellulose as the sole source of carbon and energy. *Applied and Environmental Microbiology* **82**, 4835-4845 (2016).

17 Vincent, F. *et al.* Common inhibition of both beta-glucosidases and beta-mannosidases

by isofagomine lactam reflects different conformational itineraries for pyranoside hydrolysis. *Chembiochem* **5**, 1596-1599 (2004).

18 Fiyinfoluwa A. Adesioye, T. P. M., Surendra Vikram, Bryan T. Sewell, Wolf-Dieter Schubert, Don A. Cowan. Structural characterization and directed evolution of a novel acetyl xylan esterase reveals thermostability determinants of the carbohydrate esterase 7 family. *Applied and Environmental Microbiology* **84**, 1-16 (2018).

19 Razeq, F. M. *et al.* A novel acetyl xylan esterase enabling complete deacetylation of substituted xylans. *Biotechnology for Biofuels* **11**, 74 (2018).

20 Hemsworth, G. R. *et al.* Structural dissection of a complex *Bacteroides ovatus* gene locus conferring xyloglucan metabolism in the human gut. *Open Biology* **6**, 160142 (2016).

21 Okuyama, M. *et al.* Catalytic mechanism of retaining alpha-galactosidase belonging to glycoside hydrolase family 97. *Journal of Molecular Biology* **392**, 1232-1241 (2009).

22 Sonnenburg, E. D. *et al.* Specificity of polysaccharide use in intestinal *Bacteroides* species determines diet-induced microbiota alterations. *Cell* **141**, 1241-1252 (2010).

23 Bujacz, A., Jedrzejczak-Krzepkowska, M., Bielecki, S., Redzynia, I. & Bujacz, G. Crystal structures of the apo form of beta-fructofuranosidase from *Bifidobacterium longum* and its complex with fructose. *The FEBS Journal* **278**, 1728-1744 (2011).

24 Nagem, R. A. *et al.* Crystal structure of exo-inulinase from *Aspergillus awamori*: the enzyme fold and structural determinants of substrate recognition. *Journal of Molecular Biology* **344**, 471-480 (2004).

25 Pouyez, J. *et al.* First crystal structure of an endo-inulinase, INU2, from *Aspergillus ficuum*: discovery of an extra-pocket in the catalytic domain responsible for its endo-activity. *Biochimie* **94**, 2423-2430 (2012).

26 Koropatkin, N. M., Martens, E. C., Gordon, J. I. & Smith, T. J. Starch catabolism by a prominent human gut symbiont is directed by the recognition of amylose helices. *Structure* **16**, 1105-1115 (2008).

27 Martens, E. C., Koropatkin, N. M., Smith, T. J. & Gordon, J. I. Complex glycan catabolism by the human gut microbiota: the *Bacteroidetes* Sus-like paradigm. *Journal of Biological Chemistry* **284**, 24673-24677 (2009).

28 Marie-Pierre Egloff, J. U., Lutz Haalck, Herman van Tilbeurgh. Crystal structure of maltose phosphorylase from *Lactobacillus brevis*: unexpected evolutionary relationship with glucoamylases. *Structure* **9**, 689-697 (2001).

29 Boger, M., Hekelaar, J., van Leeuwen, S. S., Dijkhuizen, L. & Lammerts van Bueren, A. Structural and functional characterization of a family GH53 beta-1,4-galactanase from *Bacteroides thetaiotaomicron* that facilitates degradation of prebiotic galactooligosaccharides. *Journal of Structural Biology* **205**, 1-10 (2019).

30 Luis, A. S. *et al.* Dietary pectic glycans are degraded by coordinated enzyme pathways in human colonic *Bacteroides*. *Nature Microbiology* **3**, 210-219 (2018).

31 Pluvinage, B. *et al.* Molecular basis of an agarose metabolic pathway acquired by a human intestinal symbiont. *Nature Communications* **9**, 1043 (2018).

32 Lee, H. S. *et al.* Cyclomaltodextrinase, neopullulanase, and maltogenic amylase are nearly indistinguishable from each other. *Journal of Biological Chemistry* **277**, 21891-21897 (2002).

33 Turkenburg, J. P. *et al.* Structure of a pullulanase from *Bacillus acidopullulyticus*. *Proteins* **76**, 516-519 (2009).
34 Larsbrink, J. *et al.* A discrete genetic locus confers xyloglucan metabolism in select human gut *Bacteroidetes*. *Nature* **506**, 498-502 (2014).
35 Matsuzawa, T., Watanabe, M., Nakamichi, Y., Fujimoto, Z. & Yaoi, K. Crystal structure and substrate recognition mechanism of *Aspergillus oryzae* isoprimeverose-producing enzyme. *Journal of Structural Biology* **205**, 84-90 (2019).
36 Rogowski, A. *et al.* Glycan complexity dictates microbial resource allocation in the large intestine. *Nature Communications* **6**, 7481 (2015).
37 Kitamura, M. *et al.* Structural and functional analysis of a glycoside hydrolase family 97 enzyme from *Bacteroides thetaiotaomicron*. *Journal of Biological Chemistry* **283**, 36328-36337 (2008).
38 Dugdale, M. L. *et al.* Importance of Arg-599 of beta-galactosidase (*Escherichia coli*) as an anchor for the open conformations of Phe-601 and the active-site loop. *Biochemistry and Cell Biology* **88**, 969-979 (2010).
39 Godoy, A. S. *et al.* Crystal structure of beta1-->6-galactosidase from *Bifidobacterium bifidum* S17: trimeric architecture, molecular determinants of the enzymatic activity and its inhibition by alpha-galactose. *The FEBS Journal* **283**, 4097-4112 (2016).
40 Miguez Amil, S. *et al.* The cryo-EM Structure of *Thermotoga maritima* beta-galactosidase: quaternary structure guides protein engineering. *ACS Chemical Biology* **15**, 179-188 (2020).
41 Cao, H., Walton, J. D., Brumm, P. & Phillips, G. N., Jr. Structure and substrate specificity of a eukaryotic fucosidase from *Fusarium graminearum*. *Journal of Biological Chemistry* **289**, 25624-25638 (2014).
42 Vandermarliere, E. *et al.* Structural analysis of a glycoside hydrolase family 43 arabinoxylan arabinofuranohydrolase in complex with xylotetraose reveals a different binding mechanism compared with other members of the same family. *Biochemical Journal* **418**, 39-47 (2009).
43 Bacic, M. K. & Smith, C. J. Laboratory maintenance and cultivation of *Bacteroides* species. *Current Protocols in Microbiology* **9**, 13C-11 (2008).
44 Déjean, G. *et al.* Synergy between cell surface glycosidases and glycan-binding proteins dictates the utilization of specific Beta(1,3)-glucans by human gut *Bacteroides*. *mBio* **11**, e00095 (2020).
